# Therapeutic assessment of a novel mitochondrial complex I inhibitor in *in vitro* and *in vivo* models of Alzheimer’s disease

**DOI:** 10.1101/2025.02.12.637918

**Authors:** Sergey Trushin, Thi Kim Oanh Nguyen, Andrea Stojacovic, Mark Ostroot, J. Trey Deason, Su-Youne Chang, Liang Zhang, Slobodan I. Macura, Toshihiko Nambara, Wenyan Lu, Takahisa Kanekiyo, Eugenia Trushina

**Author notes:** Corresponding author: Eugenia Trushina, PhD Mayo Clinic 200 First Street SW, Guggenheim Bld., Rochester, MN 55905 USA. E-mail addresses (Author 1); (Author 2), (Author 3), (Author 4), (Author 5), (Author 6), (Author 7).

## Abstract

Despite recent approval of monoclonal antibodies that reduce amyloid (Aβ) accumulation, the development of disease-modifying strategies targeting the underlying mechanisms of Alzheimer’s disease (AD) is urgently needed. We demonstrate that mitochondrial complex I (mtCI) represents a druggable target, where its weak inhibition activates neuroprotective signaling, benefiting AD mouse models with Aβ and p-Tau pathologies. Rational design and structure‒activity relationship studies yielded novel mtCI inhibitors profiled in a drug discovery funnel designed to address their safety, selectivity, and efficacy. The new lead compound C458 is highly protective against Aβ toxicity, has favorable pharmacokinetics, and has minimal off-target effects. C458 exhibited excellent brain penetrance, activating neuroprotective pathways with a single dose. Preclinical studies in APP/PS1 mice were conducted via functional tests, metabolic assessment, *in vivo* ^31^P- NMR spectroscopy, blood cytokine panels, *ex vivo* electrophysiology, and Western blotting. Chronic oral administration improved long-term potentiation, reduced oxidative stress and inflammation, and enhanced mitochondrial biogenesis, antioxidant signaling, and cellular energetics. These studies provide further evidence that the restoration of mitochondrial function and brain energetics in response to mild energetic stress represents a promising disease-modifying strategy for AD.

## Introduction

Mitochondria are essential for supporting synaptic and cognitive functions. While their primary role involves generating adenosine triphosphate (ATP) through oxidative phosphorylation (OXPHOS), recent research highlights their broader involvement in regulating cellular processes such as energy status, calcium homeostasis, metabolite production, organelle dynamics, redox balance and immune responses, among others^1,2^. Understanding the complexity of mitochondrial signaling in cellular communication opens avenues for exploring novel therapeutic strategies that improve the fundamental mechanisms of healthy aging^3,4^. The OXPHOS machinery, which is located in the inner mitochondrial membrane, consists of four electron transport chain (ETC) complexes (I - IV) and ATP synthase (complex V). Mitochondrial dysfunction, characterized by reduced OXPHOS efficiency and increased reactive oxygen species (ROS) production, is linked to neurodegenerative diseases, including AD^5^. Interestingly, recent research has shown that genetic or pharmacological inhibition of OXPHOS complexes has beneficial effects on health and lifespan across various organisms, including Drosophila, *C. elegans*, mice, and humans^6–8^. However, only mild inhibition of activity was beneficial, as the complete ablation of major OXPHOS subunits resulted in a severe phenotype and a shorter lifespan. Life-extending mechanisms include adaptive responses to energetic stress mediated by AMP-activated protein kinase (AMPK)^9^. This paradoxical approach is supported by the use of metformin, an FDA-approved mtCI inhibitor^10,11^ broadly prescribed to the aging population to treat type II diabetes mellitus^12^.

Previously, we identified the small molecule tricyclic pyrone compound CP2 as a mtCI inhibitor and demonstrated its safety and neuroprotective effects in mouse models with Aβ (APP/PS1, APP, PS1, 5xTgAD) and p-Tau (3xTgAD) pathologies^13–17^. In all the studies, CP2 reduced Aβ and p-Tau accumulation, inflammation, oxidative stress, and cognitive dysfunction while improving mitochondrial function and energy homeostasis in the brain and periphery. Treatment is safe and efficacious when administered *in utero* at the pre- or symptomatic stages of the disease. Chronic administration of CP2 for 14 months to APP/PS1 mice was safe, resulting in blockade of ongoing neurodegeneration and cognitive protection^18,19^. However, its complex synthesis, multiple chiral centers, and limited blood‒brain barrier (BBB) penetrance pose challenges for its translation to the clinic. To address these limitations, we applied rational design and extensive structure‒activity relationship (SAR) studies to develop novel compounds, where C458 (cis-(N-(pyridin-4-ylmethyl)- 2-(3-(m-tolyloxy)cyclohexyl)propan-1-amine) has the best drug‒like properties. The results of the proof-of-concept studies presented below demonstrate the efficacy of C458 in multiple *in vitro* and *in vivo* AD models, further supporting the feasibility of the development of safe and efficacious mtCI inhibitors to treat neurodegenerative diseases associated with aging.

## Results

### Design and synthesis of C458

To improve the drug-like properties of the neuroprotective mtCI inhibitor CP2 (Fig. 1a), we used a rational design to develop a new series of small molecules. These compounds were screened via a drug discovery funnel developed and optimized in the laboratory to prioritize the best candidates (Fig. 2a). The primary assays evaluated the cytotoxicity, potency, selectivity as mtCI inhibitors, and target engagement across a broad concentration range (Fig. 2b-e). Among the 24 tested compounds, C458 (cis-(N-(pyridin-4-ylmethyl)-2-(3-(m-tolyloxy)cyclohexyl)propan-1- amine) (Fig. 1a) exhibited the most favorable drug-like properties (Table 1).

**Fig. 1:**
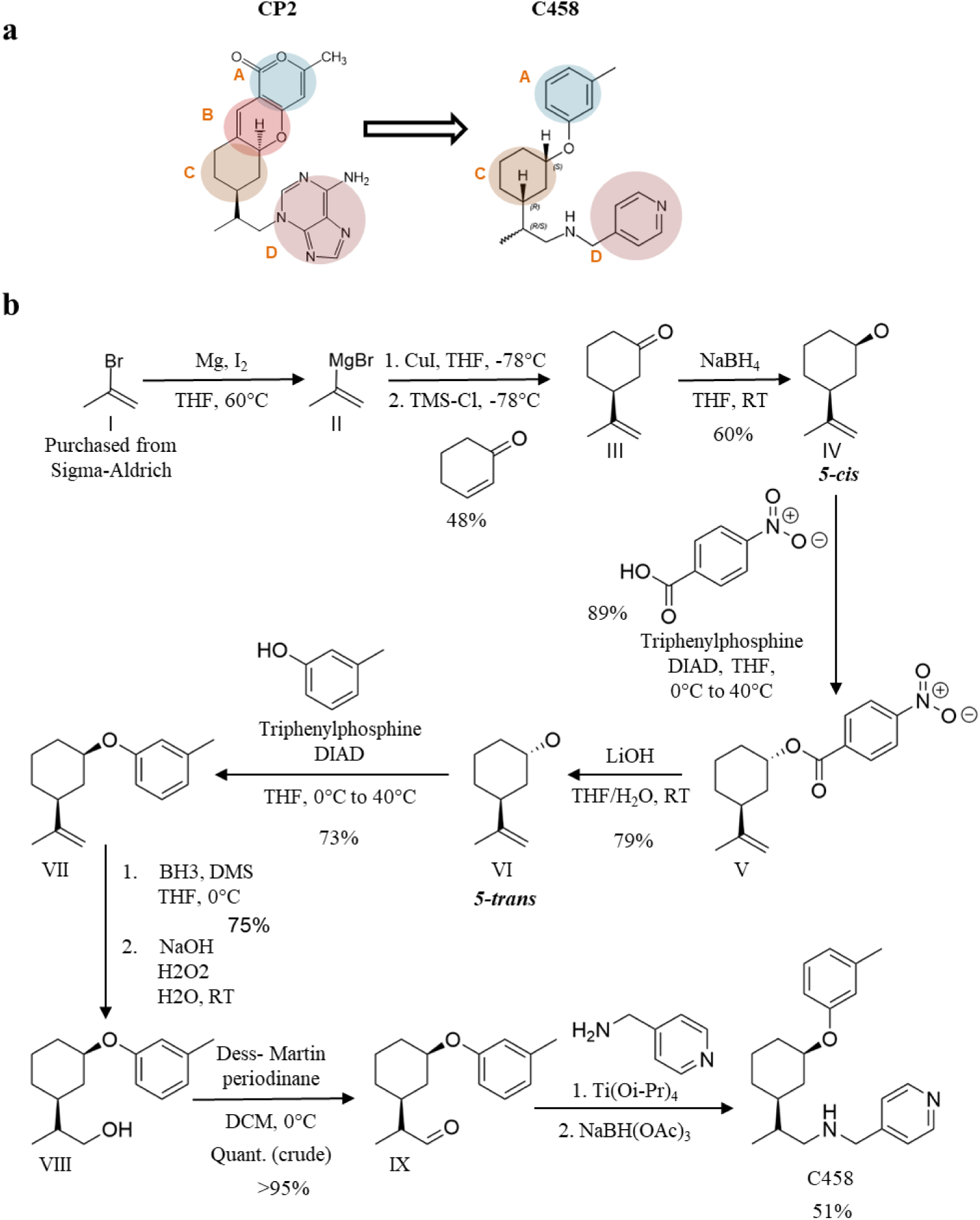
Rational design and chemical synthesis of C458. **a** The rational design of C458 was based on the CP2 structure, with structural modifications applied to rings A, B, and D. **b** The chemical synthesis of C458 involved the conversion of *5- cis*(IV) to 5-trans(VI), resulting in a cis isomer with an improved yield of 51%.

**Fig. 2:**
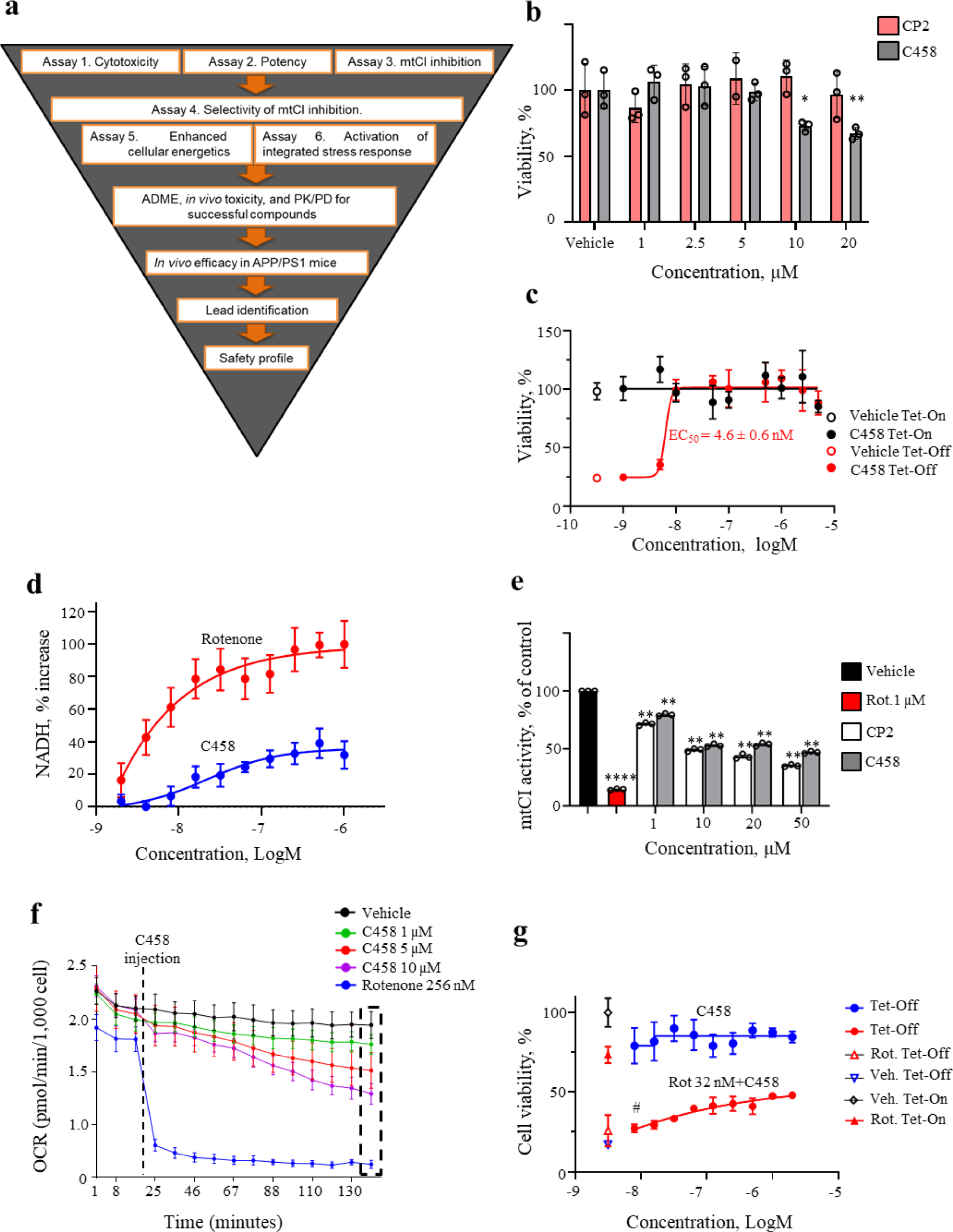
C458 is a nontoxic mild mtCI inhibitor effective against Aβ toxicity. **a** Drug Discovery Funnel for C458 identification. **b** C458 and CP2 do not cause cell death in WT mouse cortical neurons at concentrations less than 10 µM (24-hour MTT assay). *P* values were determined by one-way ANOVA. **c** C458 protects MC65 Tet-Off cells (Aβ expressed, red line) with an EC_50_ of 4.6 ± 0.6 nM. No toxicity was observed in MC65 Tet-On cells (Aβ not expressed, black line). EC_50_ values were calculated by fitting experimental data to calculated data for nonlinear regression via GraphPad Prism 10 software. The data are expressed as the means ± SDs, *n* = 3 technical replicates. **d** C458 and rotenone increase NADH levels in MC65 Tet-On cells, reflecting mtCI inhibition. NADH was measured over 24 hours. The data are expressed as the means ± SDs. **e** Compared with complete inhibition by 1 mM rotenone (NADH oxidation assay), CP2 and C458 mildly inhibited mtCI activity in isolated mouse brain mitochondria. The data are expressed as the means ± SDs, n = 4 technical replicates. f C458 reduces the oxygen consumption rate (OCR) in MC65 Tet-On cells less potently than rotenone does. The quantified data are shown in Supplementary Fig. 1h. **g**. Pretreatment with 32 nM rotenone eliminated C458 efficacy at the EC50 (#) in MC65 Tet-Off cells. The data are expressed as the means ± SDs, *n* = 4 technical replicates. **c-g** *P* values were calculated via unpaired Student’s *t* tests. Significance levels: \**P* < 0.05; \*\**P* < 0.01; \*\*\**P* < 0.001; \*\*\**P < 0.0001. All experiments were reproduced in two or more biological replicates*.

**Table 1.** Drug-like properties of CP2 and C458.

|  | **CP2** | **C458** |
| --- | --- | --- |
| Calculated log P | 3.0 | 4.4 |
| Calculated log D<sub>7.4</sub> | 2.0 | 2.0 |
| Total polar surface area | 109 | 34.2 |
| H-bond acceptors | 8 | 3 |
| H-bond donors | 2 | 1 |
| Molecular weight | 393.45 | 338.50 |

Compared with CP2, C458 has fewer hydrogen donors and acceptors, simplified stereochemistry with an open structure, fewer aliphatic carbons, and structural modifications to the A, B, and D rings (Fig. 1a). Additionally, its synthesis was simplified (Fig. 1b). By isolating the unwanted cis- isomer (IV) and converting it into the trans-isomer (VI) via a two-step sequence, the yield of the third synthetic step improved from 30% to 51%. All subsequent experiments utilized the cis-C458 isomer.

The cytotoxicity and potency of C458 (Fig. 2a-c) were evaluated in the human neuroblastoma MC65 cell line, which expresses the C99 fragment of the amyloid precursor protein (APP) under the control of a tetracycline-sensitive promoter^20^. When tetracycline is withdrawn (Tet-Off), MC65 cells express C99, which is converted to Aβ by γ-secretase. This leads to cell death within three days due to Aβ accumulation, causing oxytosis, ferroptosis, and mitochondrial dysfunction, key mechanisms involved in AD pathogenesis^21^. In the presence of tetracycline (Tet-On), MC65 cells behave normally and can be used to assess compound cytotoxicity (Fig. 2a, Assay 1). These cells provide an excellent phenotypic screening assay for drug discovery (Fig. 2a, Assay 2). The efficacy of CP2 against Aβ toxicity was initially demonstrated via this assay^22^. Like CP2, C458 did not induce cytotoxicity up to 10 μM in primary mouse cortical neurons (Fig. 2b) or in MC65 Tet-On cells (Fig. 2c, black line). Furthermore, treatment of MC65 Tet-Off cells with C458 prevented Aβ-induced cell death, with an EC_50_ of 4.6 ± 0.6 nM (Fig. 2c, red line), compared with 150 nM for CP2^22^. These results show that C458 provides superior protection against Aβ-induced mechanisms compared with CP2, with no cytotoxicity across a wide concentration range, demonstrating excellent safety.

### Target validation of C458

The target engagement (Fig. 2a, Assay 3) and selectivity of C458 for mtCI inhibition (Fig. 2a, Assay 4) were evaluated by measuring the enzymatic activities of complexes I - V in isolated mouse brain mitochondria (Fig. 2e, Supplementary Fig. 1a-c). At the efficacious concentrations established in Assay 2, C458 weakly inhibited the activity of mtCI (Fig. 2e, f) without affecting the activities of the other complexes (Assay 4, Supplementary Fig. 1a-c). Compared with rotenone, at 1 µM, C458 reduces mtCI activity by ∼15%, which completely inhibits mtCI at the same concentration. (Fig. 2e). Since mtCI inhibition by C458 at efficacious concentrations was very mild, we measured NADH accumulation in cells over time because of reduced NADH oxidation (Fig. 2d)^23^. NADH accumulation in MC65 Tet-On cells after C458 treatment occurred within the same concentration range required to rescue MC65 cells from Aβ toxicity, confirming target engagement (Fig. 2c, d). This inhibition resulted in a moderate ∼20% decrease in ATP production (Supplementary Fig. 1d). The weak inhibition of mtCI by C458 was further confirmed by a slower rate of NADH accumulation in MC65 cells than rotenone treatment (Fig. 2d). As a second measure, differences in the extent of mtCI inhibition were evaluated by measuring the oxygen consumption rate (OCR) in MC65 Tet-On cells following kinetic injections of C458 or rotenone via a Seahorse Extracellular Flux Analyzer (Fig. 2f). While rotenone immediately blocked respiration, C458 gradually and dose-dependently reduced the basal OCR (Fig. 2f, Supplementary Fig. 1f). At C458 concentrations of 1 µM or lower, changes in the OCR were minimal, and there was no increase in the extracellular acidification rate (ECAR), indicating the absence of increased glycolysis (Supplementary Fig. 1e). Only at concentrations above 10 µM did C458 significantly decrease the OCR and increase the ECAR (Supplementary Fig. 1e, f). Thus, at nanomolar concentrations, C458 weakly inhibits mtCI, protecting MC65 cells from Aβ toxicity without causing cytotoxicity or inducing glycolysis, closely mimicking the mechanisms of CP2^13^.

Our recent data (manuscript in preparation), generated via cryo-electron microscopy (cryo-EM) and purified ovine mtCI, confirmed that CP2 binds to the deep quinone site of mtCI (Qd site), one of three binding sites for rotenone^24,25^. Since C458 was designed on the basis of the CP2 structure (Fig. 1a), we hypothesize that it also binds to the quinone cavity. To test this hypothesis, we developed a competitive inhibition assay using rotenone to block the Qd site on mtCI. MC65 cells were pretreated with 32 nM rotenone for 30 minutes before C458 treatment. This rotenone concentration did not induce cell death in MC65 cells but effectively reduced the OCR by 50% within 30 minutes of rotenone addition (Supplementary Fig. 1g, h). Importantly, rotenone significantly reduced the protective activity of C458 against Aβ toxicity at a single nanomolar dose (EC_50_, Fig. 2g) and markedly diminished its effectiveness at higher concentrations (Fig. 2g). Treatment with any concentration of rotenone did not protect MC65 cells against Aβ toxicity (data not shown). These findings suggest that mild inhibition of mtCI by C458 because of its binding to the Qd-site of mtCI is crucial for its protective effect.

To further confirm C458 binding to mtCI, we developed a pull-down assay using C458 derivatives modified with spacers of varying lengths, each terminating with an NH_2_ group for immobilization onto functionalized agarose beads (Fig. 3a). Spacer-modified constructs (C458-8 with an 8-atom spacer and C458-2 with a 2-atom spacer) showed the same efficacy against Aβ toxicity in MC65 cells as unmodified C458, indicating that the spacers did not affect the activity of C458 (Supplementary Fig. 2a). In the pull-down assay, C458-8 immobilized at relatively high concentrations (1 mM) on agarose beads efficiently captured mtCI from mouse brain mitochondrial lysates. This efficiency was greater than that of C458-2 or C458-8 immobilized at a lower concentration (0.2 mM) (Supplementary Fig. 2b). The ability of C458-8-immobilized beads to capture mtCI was comparable to that of mtCI immunocapture via specific antibodies (Fig. 3b, IC). The specificity of the interaction was further demonstrated by the lack of nonspecific binding to control beads immobilized with ethanolamine (Fig. 3b, control).

**Fig. 3:**
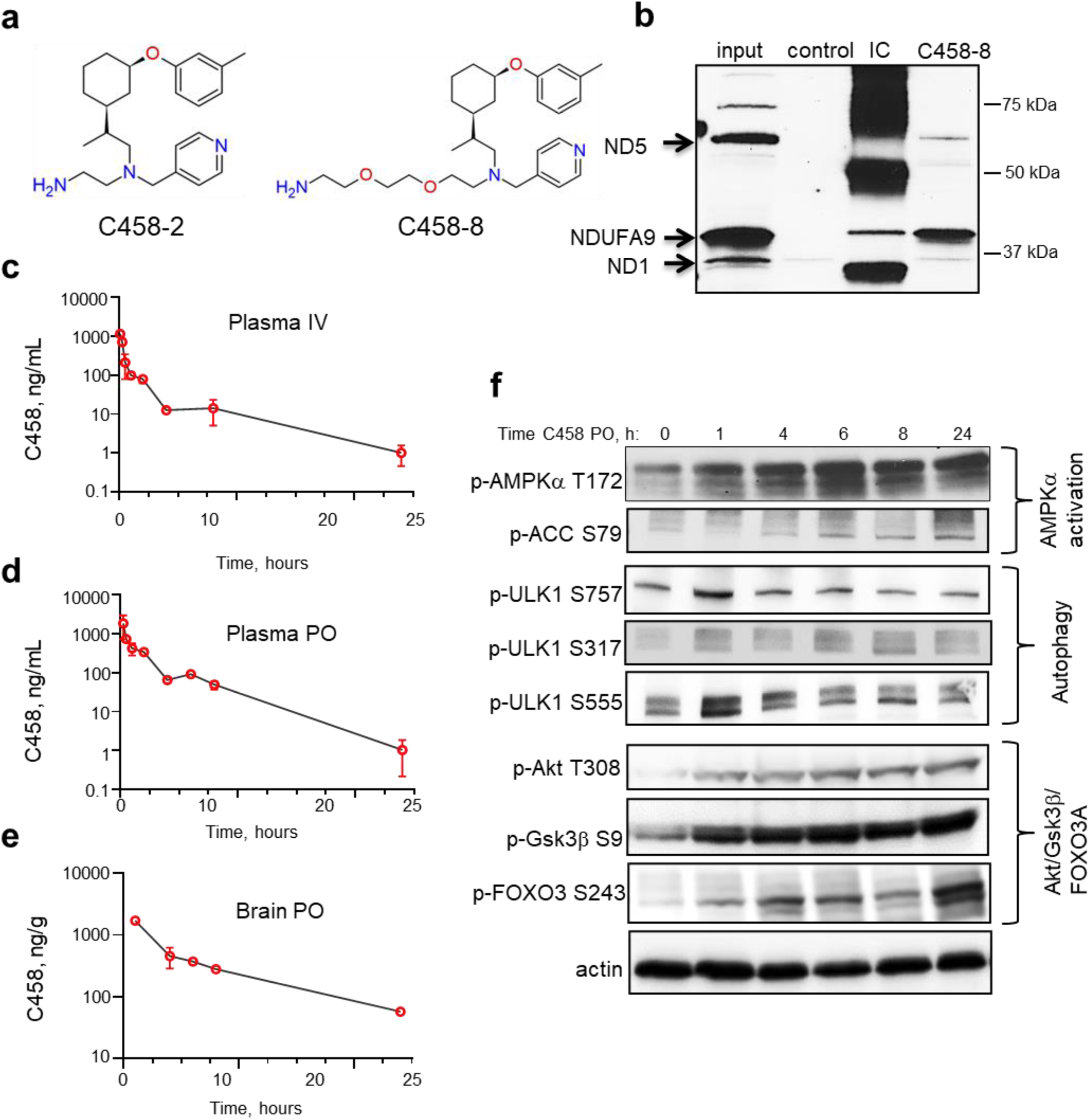
C458 penetrates the BBB, directly interacts with mtCI, and activates neuroprotective pathways in the mouse brain. **a** C458 analogs were designed with two- and eight-atom spacers on the basis of SAR studies. **b** Pull-down assay using C458-8 immobilized on agarose bead isolates mtCI from mitochondrial lysates. **c, d** Pharmacokinetics of C458 in plasma after intravenous (IV) and oral (PO) administration in C57BL/6 female mice (*n* = 3 *per* time point). **e** C458 levels in the brain after oral administration (25 mg/kg via gavage) in C57BL/6 female mice, demonstrating BBB penetration. **f** Activation of neuroprotective pathways in mouse brain tissue after a single oral dose of C458 (25 mg/kg). Western blot analysis was performed on brain tissues collected at various time points post administration. Each lane represents an individual mouse.

To assess off-target effects, C458 activity was evaluated via a 250-kinase panel provided by Nanosyn, Inc. Similar to CP2^14^, C458 did not inhibit any of the 250 kinases at concentrations of 1 µM or 10 µM (Supplementary Table 1). Taken together, these findings confirm that C458 is a specific mtCI inhibitor that is capable of protecting against Aβ toxicity in a cellular model of AD.

### C458 penetrates the BBB and activates neuroprotective mechanisms in the brain

The pharmacokinetic (PK) profile of C458 was evaluated in female wild-type mice (WT, C57BL/6) by administering 3 mg/kg intravenously (IV) or 25 mg/kg orally (PO) (Table 2).

**Table 2.** The pharmacokinetics and bioavailability of C458 in female C57BL6 mice.

| Summary | Dose | T (1/2)<sup>a</sup> | Tmax<sup>b</sup> | Cmax<sup>c</sup> | Clast<sup>d</sup> | AUClast<sup>e</sup> | AUCinf<sup>f</sup> | Ke<sup>g</sup> | Cl/F<sup>h</sup> | F (%)<sup>i</sup> | Brain/<br>Plasma<br>ratio |
| --- | --- | --- | --- | --- | --- | --- | --- | --- | --- | --- | --- |
|  | (mg/kg) | hr | hr | ng/mL | ng/mL | ng/mL*h | ng/mL*h |  | mL/hr/kg |  |  |
| IV<br>Plasma | 3 | 4.21 | 0.08 | 775.33 | 1.02 | 695 | 701 | 0.16 | 4280 |  |  |
| PO<br>Plasma | 25 | 2.61 | 0.25 | 1819.57 | 1.05 | 2768 | 2772 | 0.27 | 9030 | 47.39 |  |
| PO<br>Brain | 25 | 6.77 | 1.00 | 1705 | 57.02 | 8266 | 8823 | 0.10 | 3024 |  | 3.18 |

Summary of the PK studies of C458 delivered to C57BL/6 female mice ( $n = 3$ per time point) via the IV route (3 mg/kg) or orally (PO route) (25 mg/kg via gavage). a, Half-life; b, time to maximum concentration; c, maximum concentration; d, last measurable concentration; e, area under the curve from the last measurable concentration; f, area under the curve extrapolated to infinity; g, elimination rate constant; h, clearance adjusted for bioavailability; i, bioavailability.

Blood and brain tissues were collected from each mouse over a 24-hour period (Fig. 3c-e), and C458 levels in frozen plasma and brain samples were measured via LC‒MS/MS. C458 demonstrated high BBB penetrance, with a brain/plasma ratio of 3.18, and good bioavailability (47.4%, Table 2). Following PO administration, the half-life (T1/2) of C458 was 2.6 hours in plasma and 6.8 hours in the brain, with a maximum concentration (Cmax) of 1819.6 ng/mL reached within one hour (Fig. 3c, d; Table 2). BBB penetrance was further confirmed by an efflux ratio (ER = 1.1) in Madin‒Darby canine kidney (MDCKII) cells transfected with the human M*DR1* gene encoding p-glycoprotein (Supplementary Table 2).

To assess target engagement and activation of neuroprotective mechanisms, Western blot (WB) analysis was performed using brain tissues from the PK study. Within one hour of PO administration, C458 activated AMPKα, a key regulator of cellular energy homeostasis, resulting in the downstream inactivation of acetyl-CoA carboxylase 1 (ACC1) (Fig. 3f)^26^. Consistent with AMPK activation, the phosphorylation of Unc-51, similar to Autophagy Activating Kinase 1 (ULK- 1), at S317 and S555 indicated autophagy activation (Fig. 3f). Additionally, C458, like CP2^13–15^, promoted Akt phosphorylation (glucose metabolism) and inactivated Gsk3β and FOXO3A (tau phosphorylation and neuroprotection), engaging multiple neuroprotective mechanisms^27^ (Fig. 3f). These effects persisted for 24 hours post administration. These findings indicate that C458 efficiently crosses the BBB, promptly engages mtCI, and activates a neuroprotective integrated signaling cascade^18^.

To evaluate *in vivo* toxicity, independent groups of 2-month-old male and female WT mice were treated with vehicle or 25 mg/kg or 50 mg/kg C458 via the drinking water *ad libitum*. One month later, the adult and newborn mice were sacrificed, and the liver, spleen, heart, lungs, kidneys, sex organs, and brain were subjected to histopathological examination. No developmental or tissue pathology was observed in C458-treated mice (Supplementary Table 3). Thus, similar to CP2^13^, C458 showed no developmental toxicity or adverse effects after 30 days of treatment at 50 mg/kg/day.

### C458 treatment alleviates the AD-like phenotype in APP/PS1 mice

The preclinical *in vivo* efficacy of C458 was evaluated in two independent cohorts of APP/PS1 mice aged 2.5 to 10.5 months (Fig. 4a). Trial 1 assessed the safety of C458 administration and its impact on behavior, cognitive function, and inflammation, whereas Trial 2 focused on reproducibility and effects on metabolism, brain and peripheral energy homeostasis, synaptic function, and mechanisms of action. Data from tests applied in both trials were combined for analyses.

**Fig. 4:**
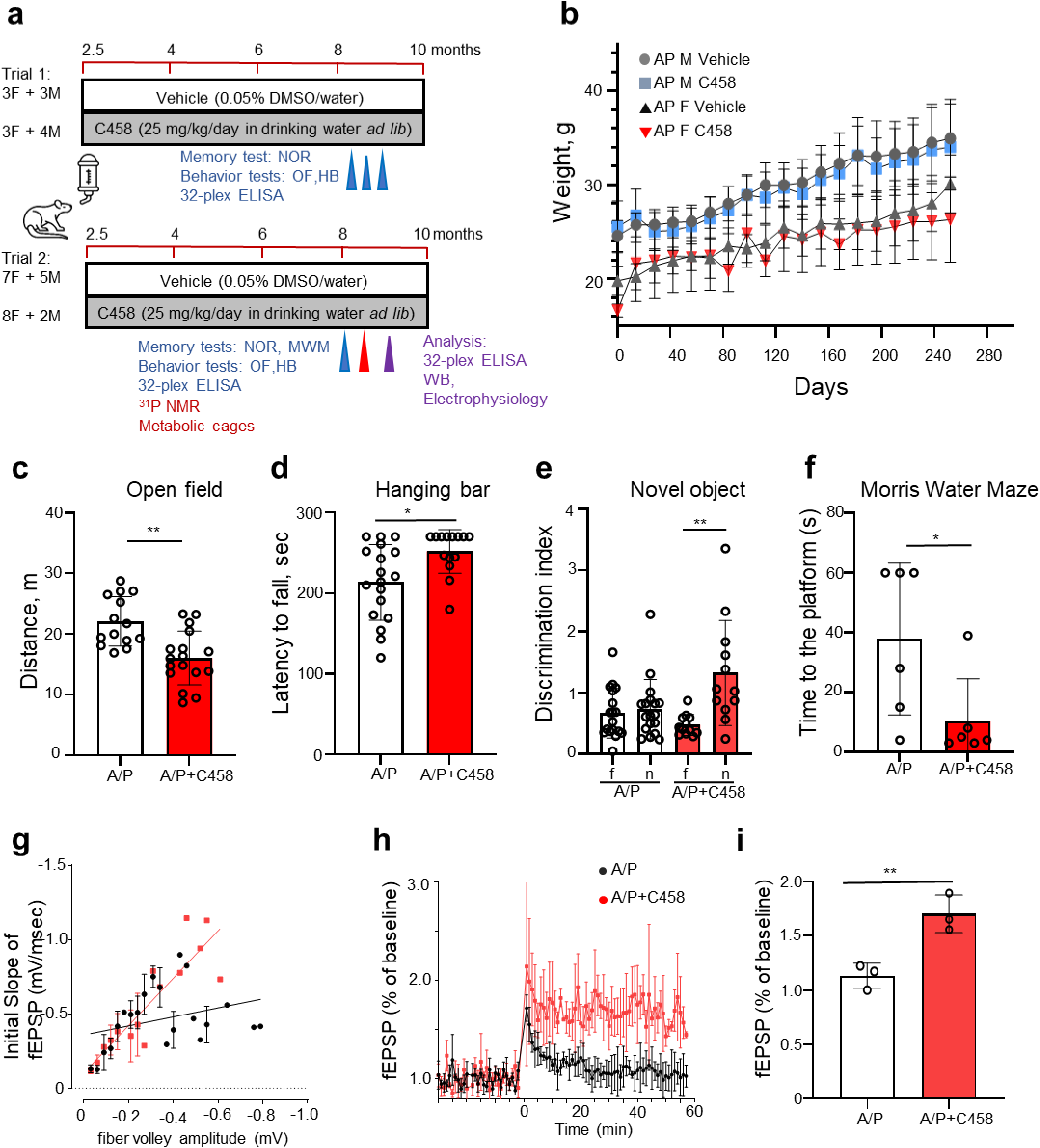
C458 treatment improves motor and cognitive functions and LTP in APP/PS1 mice. **a** Timeline of two trials in which male and female APP/PS1 mice were treated with either vehicle or C458 for 7.5 months, with tests conducted in each trial. **b** Body weights of APP/PS1 mice treated with vehicle or C458 over the study duration (data from Trials 1 and 2 combined). **c** C458 treatment reduces hyperactivity in APP/PS1 mice in the open field test. **d** C458 treatment enhances motor strength and coordination in the hanging bar test. **e, f** C458 treatment improved performance in the novel object recognition test and in the Morris water maze. Data for Trials 1 and 2 are shown in **c - e**, *n* = 11 - 16 mice *per* group. Data from Trial 2 are shown in **f**, *n* = 6 mice *per* group. **g** C458 treatment improved basal synaptic strength. The relationships between the initial slopes of fEPSPs and presynaptic fiber volley amplitudes were significantly greater in the C458-treated APP/PS1 group than in the vehicle-treated APP/PS1 group. **h, i** C458 treatment enhances LTP in APP/PS1 mice; *n* = 2–3 slices from 3 mice *per* group. Representative traces are shown as the mean ± SD *at each* time point. **g-i** Data from Trial 2. The data were analyzed via one-way ANOVA, with the exception of the NOR test, which was analyzed via paired Student’s t tests. \**P* < 0.05, \*\**P* < 0.01, \*\*\**P* < 0.001, \*\*\*\**P* < 0.0001.

As the study was not designed to evaluate sex-specific differences, data from male and female mice were analyzed together. Nontransgenic littermates were not included in this study. APP/PS1 mice received 25 mg/kg/day C458 or vehicle (0.05% DMSO) via *ad libitum* drinking water. The brain and plasma C458 concentrations used in the present study ranged from 10 to 400 nM, which is consistent with mtCI engagement based on the PK data (Supplementary Table 4). C458-treated APP/PS1 mice exhibited no toxicity or side effects and gained weight throughout the study (Fig. 4b).

Compared with vehicle treatment, functional tests revealed significant improvements in motor and cognitive performance in C458-treated APP/PS1 mice. Compared with their vehicle-treated counterparts, the C458-treated APP/PS1 mice demonstrated reduced hyperactivity in the open field test, enhanced motor coordination and muscle strength in the hanging bar test (Fig. 4c, d), improved attention and nonspatial declarative memory in the novel object recognition test, and better spatial memory and learning in the Morris water maze (Fig. 4e, f). Given that synaptic loss strongly correlates with cognitive dysfunction in AD^28,29^, synaptic function was examined to investigate the basis of improved cognitive performance. Extracellular recordings were used to measure field excitatory postsynaptic potentials (fEPSPs) in the CA1 region of acute hippocampal slices (Fig. 4g-i). Compared with vehicle treatment, C458 treatment increased synaptic strength in APP/PS1 mice and significantly improved long-term potentiation (LTP) (Fig. 4h, i). These findings indicate that chronic C458 treatment is safe and significantly improves cognitive and behavioral AD-like phenotypes in APP/PS1 mice.

### C458 treatment enhances metabolic flexibility and energy homeostasis in the brain and periphery

CP2-treated APP/PS1 mice presented improved energy homeostasis in the brain and periphery, as we reported previously^14^. To investigate the effect of C458 on metabolism, indirect calorimetry (CLAMS) was employed. Compared with their vehicle-treated counterparts, APP/PS1 mice treated with C458 presented increased carbohydrate oxidation and metabolic flexibility, as indicated by the respiratory exchange ratio (RER), a critical marker of the ability to adapt to changes in metabolic or energy demands consistent with improved glucose utilization (Fig. 5a-e)^30^. C458 treatment also significantly improved glucose tolerance in APP/PS1 mice (Fig. 5f), suggesting enhanced glucose metabolism.

**Fig. 5:**
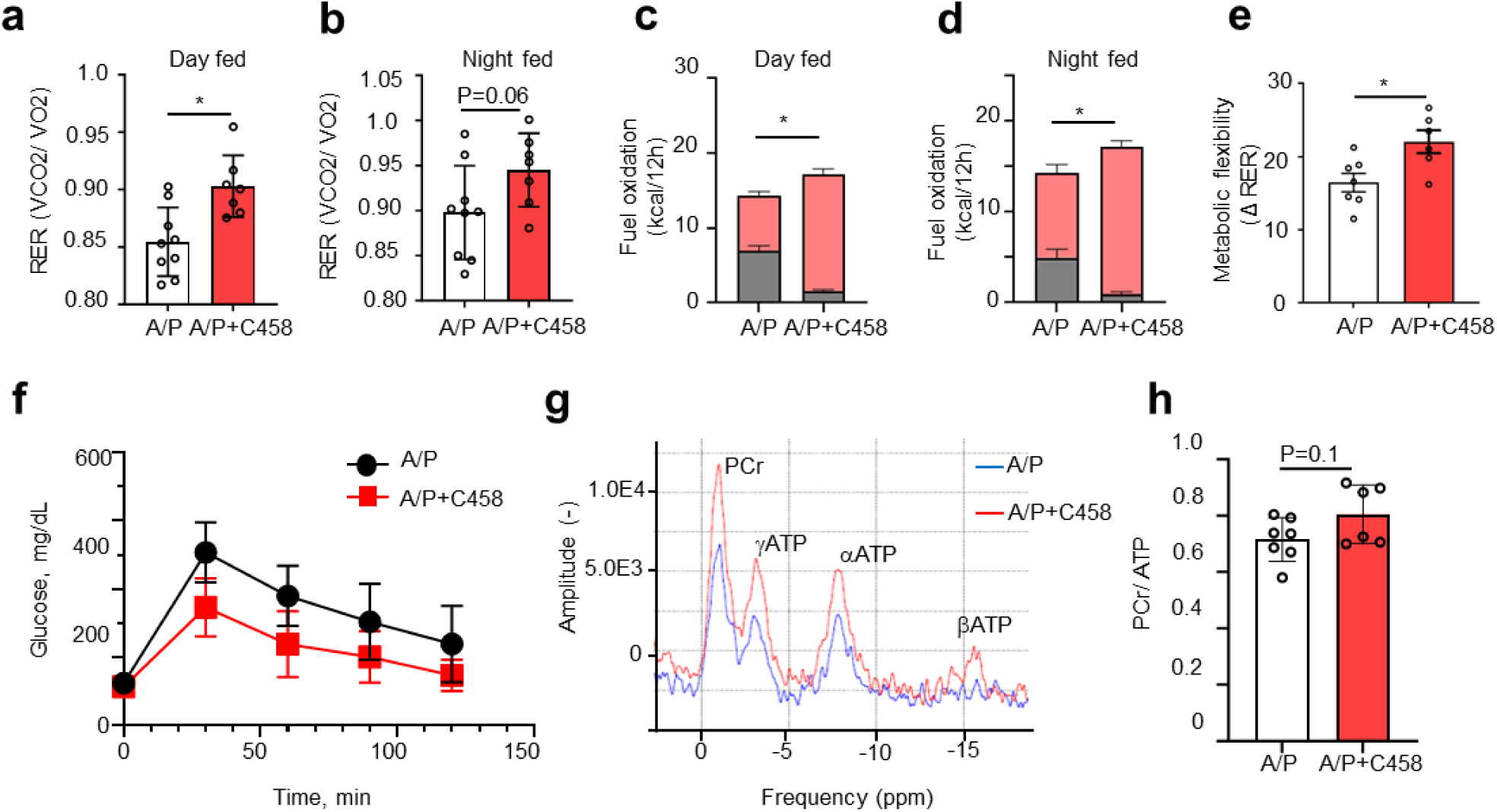
C458 treatment enhances metabolic function and glucose tolerance and preserves ATP in the brains of APP/PS1 mice. a, **b** Changes in the respiratory exchange ratio (RER) over 44 hours during *ad libitum* feeding for all treatment groups. **c, d** C458 treatment increased glucose oxidation during *ad libitum* feeding, as shown by the CLAMS data. The gray bars represent fat consumption; the orange bars represent carbohydrate and protein oxidation. **e** Metabolic flexibility is improved in C458-treated APP/PS1 mice, as evidenced by their ability to switch from carbohydrate metabolism to fat metabolism between the feeding and fasting states. **a–e,** *n* = 7–9 mice *per* group. **f** C458 improves glucose tolerance, as assessed by the intraperitoneal glucose tolerance test (IPGTT). **g** Representative *in vivo* ^31^P-NMR spectra comparing vehicle-treated (blue) and C458-treated (red) APP/PS1 mice. **h** Phosphocreatine/ATP ratio calculated from ^31^P-NMR spectra, demonstrating ATP preservation in C458-treated APP/PS1 mice, *n* = 6–7 mice *per* group. The data in **a, b, f, g, and h** are presented as the means ± SDs; those in c-e are presented as the means ± SEMs. Statistical analysis was performed via one-way ANOVA. \**P* < 0.05, \*\**P* < 0.01, \*\*\**P* < 0.001, \*\*\*\**P* < 0.0001. The data shown are from Trial 2.

Since C458 inhibits mtCI, we examined whether chronic C458 administration affects ATP levels in the brain. We utilized ^31^P-nuclear magnetic resonance (^31^P-NMR) spectroscopy, a noninvasive, *in vivo* method for real-time measurements of key energy metabolites, including phosphocreatine (PCr), inorganic phosphate (Pi), and the α, β, and γ phosphate groups of ATP (Fig. 5g, h)^31^. This translational method is also used to assess energy levels in humans, providing crucial insights into brain energy dynamics.

After 7 months of C458 treatment, APP/PS1 mice tended toward an increased PCr/ATP ratio. In cells, PCr serves as a critical energy reservoir, buffering ATP levels during periods of high energy demand^31^. The observed increase in the PCr/ATP ratio suggests improved maintenance of energy reserves (Fig. 5g, h). The increased PCr/ATP ratio observed in APP/PS1 mice indicates that C458 activates mechanisms that enhance mitochondrial function and energy efficiency within brain cells. Collectively, these findings suggest that C458 treatment promotes metabolic adaptation, thereby preserving or even enhancing energy homeostasis in the brain while also improving glucose metabolism in the periphery, indicating that C458 could help regulate systemic energy homeostasis.

### C458 treatment mitigated oxidative stress and inflammation

AD is associated with increased oxidative stress and ROS generation, altering redox homeostasis^5,32^. Nicotinamide adenine dinucleotide phosphate (NADPH) is a crucial molecule for maintaining the cellular redox balance and functions as an electron donor to reduce glutathione (GSH), a major cellular antioxidant^33^. In AD mouse neurons, NADPH depletion occurs upstream of GSH exhaustion, contributing to oxidative stress and promoting neuronal death^34^. Similarly, the expression of Aβ in MC65 cells induces ROS production, depleting GSH and triggering oxytosis/ferroptosis-induced cell death^21,35^. To assess whether C458 provides protection against oxidative stress following Aβ accumulation in MC65 cells, we challenged MC65 cells with different doses of hydrogen peroxide (H_2_O_2_) after pretreatment with C458 or vehicle for 24 hours (Fig. 6a).

**Fig. 6:**
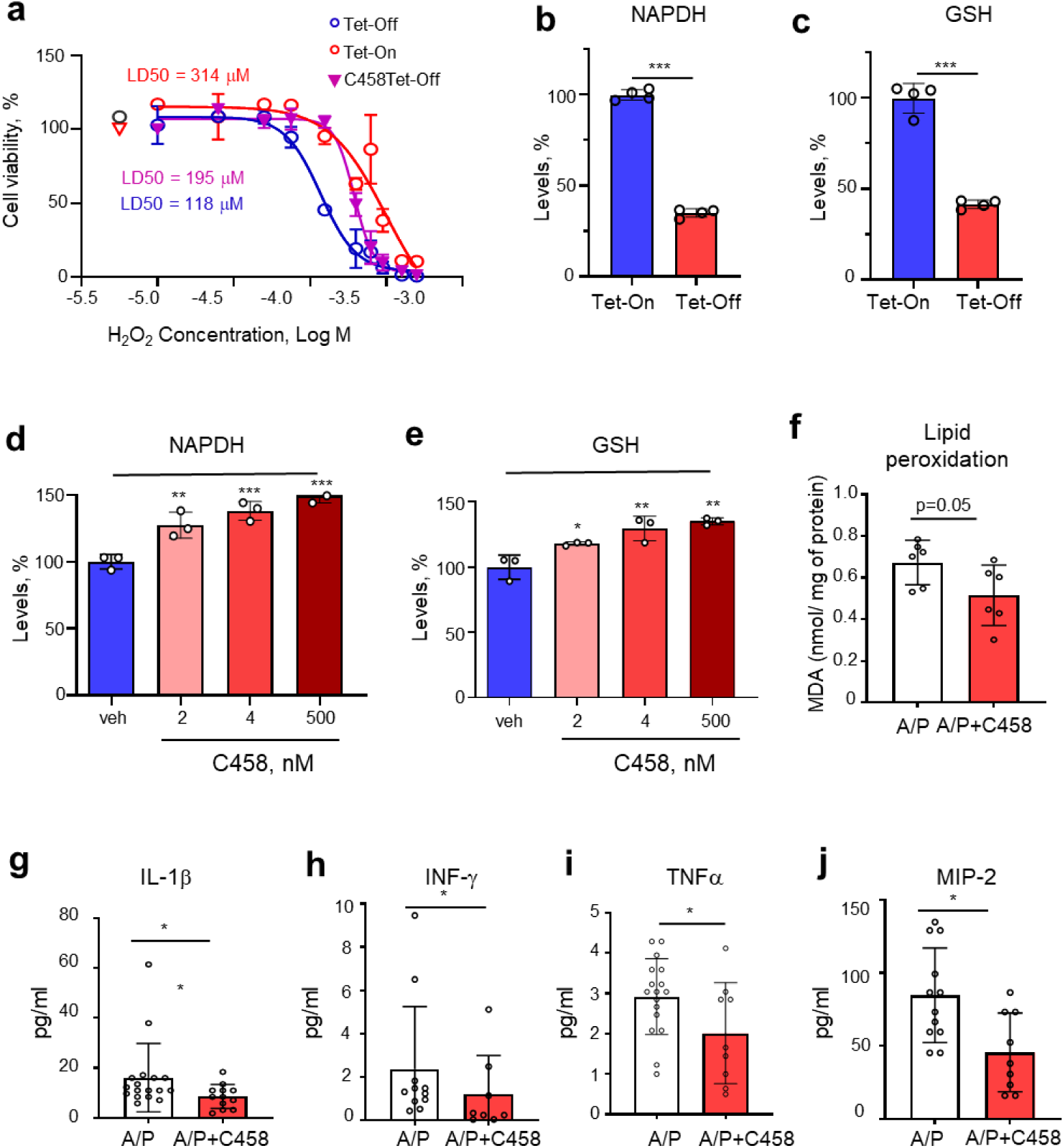
C458 treatment increases resistance to oxidative stress and improves redox balance in MC65 Tet-Off cells. **a** C458 treatment increases resistance to oxidative stress caused by H_2_O_2_ treatment in MC65 cells expressing Aβ (Tet-Off condition). LD50 values were calculated through nonlinear regression via GraphPad Prism software. The data are expressed as the means ± SDs. **b-c** Expression of Aβ (Tet-Off condition) in MC65 cells significantly depletes NAPDH and GSH levels. **d, e** C458 restores NAPDH and GSH levels in MC65 Tet-Off cells. The data are presented as the means ± SD. **f** Levels of malondialdehyde (MDA) were reduced in the brain tissue of C458-treated APP/PS1 mice; *n* = 6 mice *per* group. **g-i** C458 reduces the levels of proinflammatory cytokines and chemokines in the plasma of APP/PS1 mice. *n* = 15–20 mice *per* group. Data from Trials 1 and 2 combined. Statistical analysis was performed via unpaired Student’s *t* tests for comparisons between the Tet-On and Tet-Off conditions and between the vehicle- and C458-treated groups. \**P* < 0.05; \*\**P* < 0.01; \*\*\**P* < 0.001; \*\*\*\**P* < 0.0001.

The expression of Aβ significantly sensitized cells to H_2_O_2_-induced oxidative stress, reducing their survival (Fig. 6a). This was associated with a significant decrease in NADPH and GSH levels compared with those in MC65 Tet-On cells (Fig. 6b, c). C458 treatment restored resistance to H_2_O_2_ to a level comparable to that observed in MC65 Tet-On cells (Fig. 6a). Consistently, C458 significantly increased NADPH and GSH levels in MC65 Tet-Off cells in a dose-dependent manner (Fig. 6d, e).

These findings highlight that restoring NADPH and GSH levels via C458 treatment is crucial for mitigating Aβ-induced oxidative stress and ferroptosis in MC65 cells.

We next measured lipid peroxidation, a well-established marker of oxidative stress, in the brain^36^. After chronic C458 treatment in APP/PS1 mice, the levels of malondialdehyde (MDA), a measure of lipid peroxidation, tended to decrease (Fig. 6f), indicating that C458 treatment effectively mitigates oxidative stress in both *in vitro* and *in vivo* models of AD. Aβ accumulation in MC65 cells depletes GSH, increases ROS and lipid peroxidation, and leads to NF-kB activation and the synthesis of proinflammatory cytokines^21,37^. To determine whether the reduction in oxidative stress after C458 treatment alleviates inflammation^38^, we measured the levels of 32 cytokines and chemokines in the plasma of APP/PS1 mice (Fig. 6g-j). C458 significantly decreased the plasma levels of the proinflammatory cytokines IL-1β, TNFα, and IFN-γ, as well as the chemokine MIP-2, indicating that the treatment alleviates chronic inflammation in APP/PS1 mice (Fig. 6g-j).

### Neuroprotective mechanisms of C458 require AMPK activation

Our previous studies suggested that the efficacy of mtCI inhibitors in AD models requires AMPK activation^13,14^. To confirm that AMPK is essential for C458-dependent neuroprotective mechanisms, we used AMPKα1/α2 knockout mouse embryonic fibroblasts (MEFs) (a generous gift from Dr. B. Violet)^39^. In these MEFs, the absence of AMPK results in constitutive activation of the NF-κB inflammatory pathway, as evidenced by increased degradation of IκBα, a key inhibitor of NF-κB, and subsequent nuclear translocation of the transcription factor p65 NF-κB^40^.

In wild-type (WT) MEFs, C458 activates AMPKα, leading to inactivation of its downstream target ACC and an increase in the levels of antioxidants, including heme oxygenase-1 (HO-1), as well as elevated levels of IκBα (Fig. 7a). These findings indicate that C458 suppresses inflammation through the AMPK-mediated inhibition of NF-κB and enhanced antioxidant capacity. Furthermore, in WT MEFs, C458 stimulates mitochondrial biogenesis, as evidenced by increased levels of peroxisome proliferator-activated receptor gamma coactivator 1-alpha (PGC1α) and OXPHOS complexes I-V (Fig. 7a, Supplementary Fig. 3, Supplementary Fig. 7). In contrast, vehicle-treated AMPKα1/α2-deficient MEFs presented decreased baseline levels of PGC1α, OXPHOS complexes I, II, and V, HO-1, NQO-1, and IκBα (Fig. 7a, Supplementary Fig. 3, Supplementary Fig. 7). In AMPKα1/α2-deficient MEFs, C458 failed to promote mitochondrial biogenesis, antioxidant defenses, or anti-inflammatory responses, demonstrating that AMPK is essential for the activation of neuroprotective adaptive stress response pathways.

**Fig. 7:**
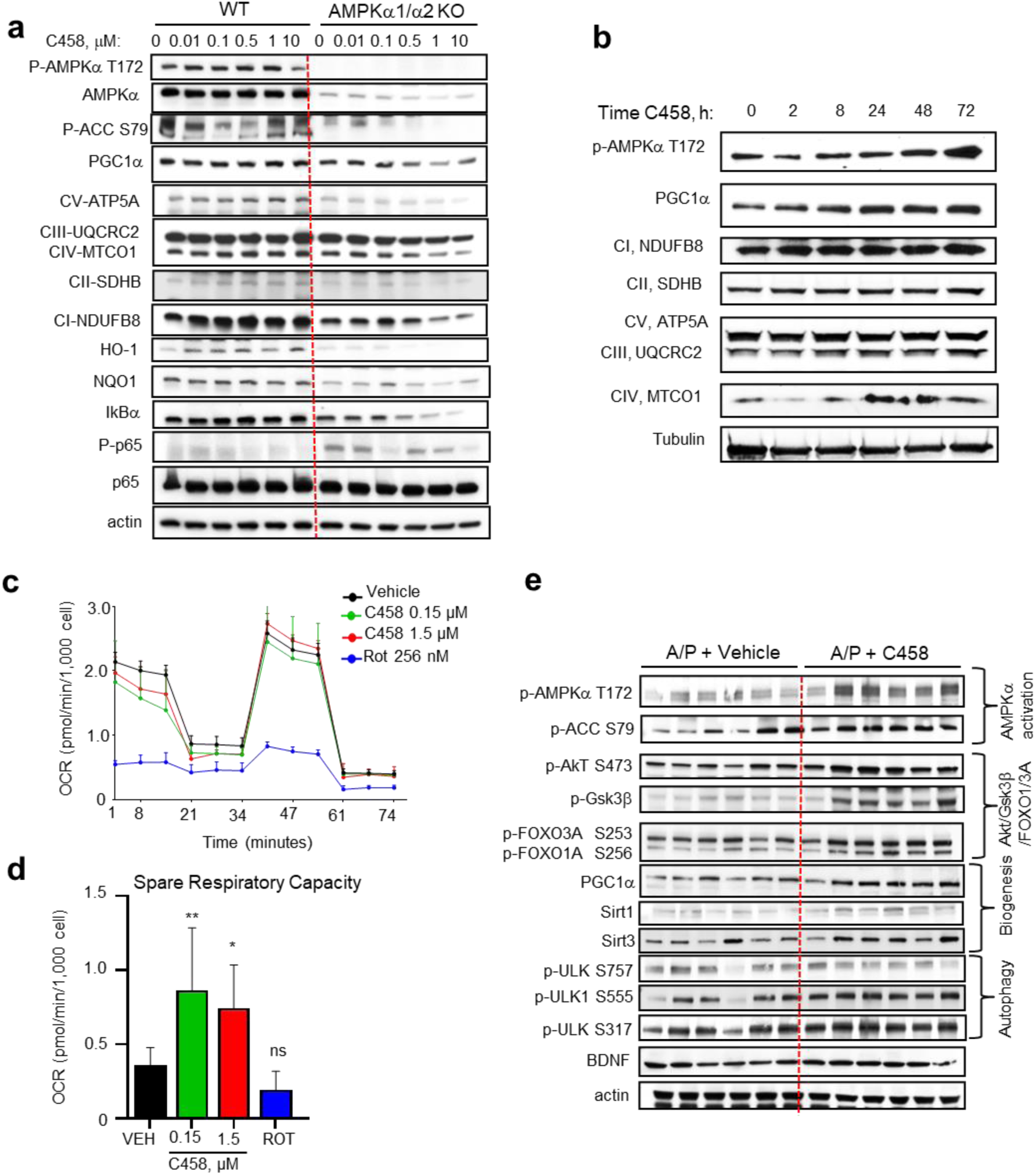
C458 treatment requires AMPK activation, which induces multiple neuroprotective mechanisms *in vivo* and *in vitro*. **a** C458 treatment activated AMPKα and PGC1α, increased the levels of OXPHOS complexes and antioxidants, and mitigated NF-kB signaling in WT MEFs but not in AMPKα1/α2-deficient MEFs. **b** In primary cortical mouse neurons, C458 (2.5 μM) treatment activated AMPKα in a time-dependent manner and induced mitochondrial biogenesis, as evidenced by PGC1α activation and increased levels of OXPHOS complexes I–V. **c** In SH- SY5Y APP_SWE_ cells, C458 treatment decreased the basal OCR while increasing the maximal OCR compared with terminal OCR inhibition via rotenone (256 nM). **d** C458 treatment improved SRC in SH-SY5Y APP_SWE_ cells. The SRC was calculated for the OCR data shown in (**c**). **e** In the brains of APP/PS1 mice (Trial 2; *n* = 6 mice *per* group), C458 treatment activates AMPKα, induces mitochondrial biogenesis via PGC1α activation, reduces GSK3β and FOXO1/3A activity, and enhances autophagy and the levels of BDNF and Sirtuins 1 and 3. The data are presented as the means ± SDs. Statistical analysis was performed via unpaired Student’s *t* tests to compare the vehicle- and C458-treated groups. \**P* < 0.05; \*\**P* < 0.01; \*\*\**P* < 0.001; \*\*\*\**P* < 0.0001.

Consistent with the observed increase in mitochondrial biogenesis and the levels of OXPHOS complexes in MEFs, C458 treatment increased PGC1α and the levels of OXPHOS complexes in primary mouse neurons over a time course of 72 hours (Fig. 7b, Supplementary Fig. 4, Supplementary Fig. 8). Additionally, C458 enhanced bioenergetics in human neuroblastoma SH- SY5Y cells stably transfected with Swedish mutant amyloid precursor protein (APPswe), another cellular AD model characterized by increased Aβ production^41^ (Fig. 7c, d). Specifically, compared with vehicle-treated cells, C458-treated cells presented increased spare respiratory capacity (SRC), an indicator of mitochondrial fitness and the ability to produce energy under stress conditions. Notably, treatment with rotenone, a terminal mtCI inhibitor, did not improve SRC (Fig. 7c, d). This enhanced SRC aligns with the improved energy homeostasis observed following C458 treatment in APP/PS1 mice.

To confirm the *in vivo* neuroprotective mechanisms of C458, WB analysis was performed on brain tissue from APP/PS1 mice treated with vehicle or C458 for 7.5 months (Fig. 7e). C458 treatment reached efficacious concentrations in mouse brains (Supplementary Table 4) and activated key neuroprotective pathways, including the AMPKα, ULK-1, and PGC1α pathways. It also inactivated critical metabolic regulators, such as ACC, Gsk3β, and the transcription factors FOXO1A/FOXO3A (Fig. 7e, Supplementary Fig. 5, Supplementary Fig. 9). We also observed increases in the levels of sirtuin (SIRT) 1 and 3 and brain-derived neurotrophic factor (BDNF), indicating enhanced support for neuronal function. These findings demonstrate that, like CP2, C458 activates multiple neuroprotective mechanisms *in vitro* and *in vivo*, reducing inflammation and augmenting cellular bioenergetics, redox balance, and mitochondrial function.

To explore C458 as a potential drug candidate, a comprehensive safety assessment was conducted via the Cerep Safety Screen 44 panel, which includes various human receptors, neurotransmitters, and ion channels. At a concentration of 10 µM, which is 1,000× greater than the EC_50_ concentration used in efficacy Assay 2 (Fig. 2a), C458 inhibited several human receptors, including hERG (Supplementary Table 5), a critical potassium channel subunit essential for maintaining normal cardiac electrical activity. Owing to the absence of this receptor in mice, this potential cardiac liability cannot be detected in preclinical *in vivo* studies. This highlights the importance of introducing hERG safety assessments earlier in the drug discovery pipeline to guide the development of selective, potent, and safe mtCI inhibitors.

### C458 and CP2 reduce the levels of Aβ and p-Tau in iPSC-derived cerebral organoids from patients with sporadic AD

To assess the translational potential of mtCI inhibitors, we utilized iPSC-derived cerebral organoids from AD patients carrying two *APOE4* alleles, which represent sporadic AD^42^. These well-characterized 3D organoids exhibit an *APOE4/4*-associated phenotype, including elevated levels of Aβ, p-Tau T181, apoptosis, and synaptic loss^42^. Following 48 hours of treatment with CP2 or C458, the organoids were lysed in RIPA buffer, and the levels of Aβ40, Aβ42, p-Tau T181, and total tau were quantified via ELISA^42^ (Supplementary Figure 10). Both the CP2 and C458 treatments significantly reduced the intracellular Aβ42/Aβ40 ratio and the p-Tau/Tau ratio (Fig. 8a-d).

**Fig 8:**
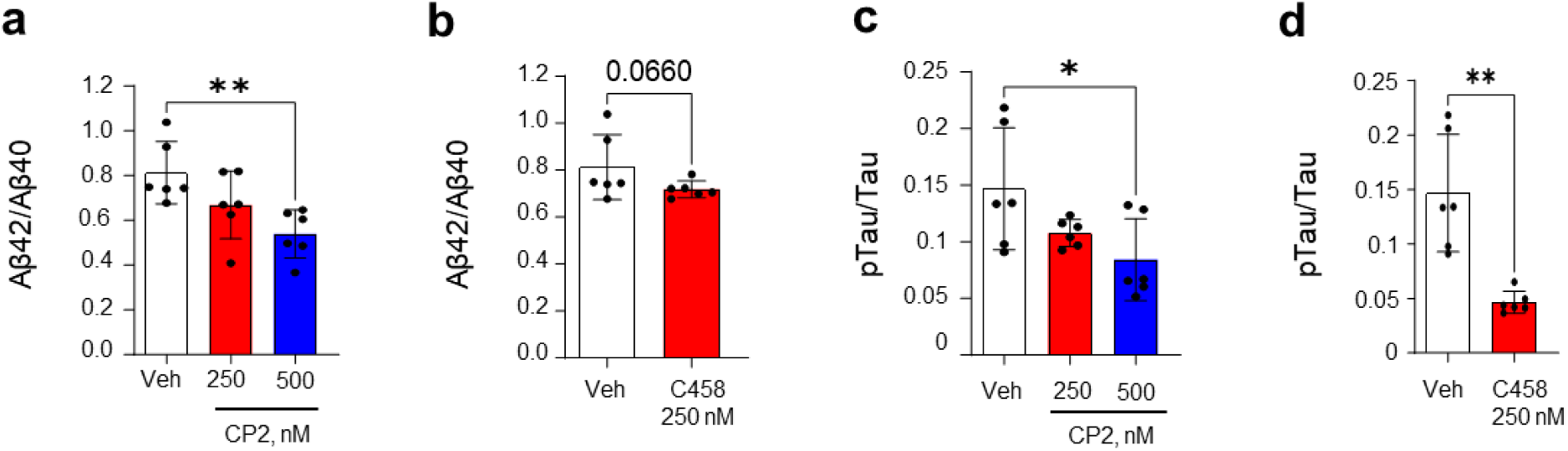
Treatment with the mtCI inhibitors CP2 and C458 reduces the levels of Aβ and pTau T181 in iPSC-derived organoids from patients with sporadic AD. **a-d** The levels of Aβ40, Aβ42, p-Tau T181 and total tau in RIPA fractions from human iPSC-derived organoids treated with vehicle, CP2, or C458 were measured via ELISA. The data were normalized to the total protein concentration of each sample (*n* = 3 organoids *per* treatment, measured in duplicate). The data are expressed as the means ± SD. Statistical analysis was performed via Student’s *t* tests or one-way ANOVA. \**P* < 0.05, \*\**P* < 0.01, \*\*\**P* < 0.001, \*\*\*\**P* < 0.0001.

These results indicate that mild mtCI inhibition mitigates Aβ and Tau pathology in human brain cells, which is consistent with data generated in 3xTgAD and APP/PS1 mice treated with C458 or CP2^14,15^. Furthermore, the reduction in the intracellular Aβ42/Aβ40 ratio is particularly notable given that *APOE4* variants are linked to impaired amyloid clearance and increased Aβ42 accumulation. Collectively, these findings demonstrate that mtCI inhibition ameliorates key AD hallmarks in iPSC-derived cerebral organoids from patients with sporadic AD.

## Discussion

The development of disease-modifying treatments for AD is hindered by an incomplete understanding of the diverse mechanisms that underlie heterogeneous and varied responses to interventions^43,44^. While lecanemab^45^, a disease-modifying anti-amyloid antibody, has been approved by the FDA, its side effects raise concerns with its application. Alternative approaches targeting the underlying mechanisms of AD pathobiology must be considered. Targeting mtCI with small molecules capable of restoring multiple mechanisms essential for AD represents a promising strategy^18^.

We validated the feasibility of using safe and efficacious mtCI inhibitors for AD treatment. Building on the previously characterized mtCI inhibitor CP2, we developed a novel small molecule, C458, with improved drug-like properties; PK; and absorption, distribution, metabolism, and excretion (ADME) profiles. C458 demonstrated strong neuroprotective effects in both *in vitro* and *in vivo* models of AD. C458 reduced Aβ and p-Tau levels in human brain organoids generated from the iPSCs of patients with sporadic AD, emphasizing its translational potential. C458 exhibited potency against Aβ toxicity at nanomolar concentrations, with no detectable cytotoxicity in neural cells up to 10 µM, and safe application in APP/PS1 mice over 7 months, indicating a good safety profile. C458 effectively crosses the BBB and provides neuroprotection in APP/PS1 mice within a therapeutic brain concentration range of 100–1000 nM. The concentration of C458 in the brains of APP/PS mice and the brain/plasma ratio were much greater than those reported for CP2^14^, indicating significantly greater BBB penetration, a lack of excessive efflux, and enhanced stability. Compared with the recently developed mtCI inhibitors for cancer therapy, C458 has an outstanding safety profile. Unlike phenformin and IACS-01759, which can induce lactic acidosis due to strong inhibition of mtCI and enhanced glycolysis^46,47^, C458 mildly inhibits mtCI at efficacious doses. Its mechanism of action leads to a gradual accumulation of NADH, increasing by up to 25% at 1 µM over time (Fig. 2d). This mild and sustained mtCI inhibition results in a modest 10–15% reduction in ATP levels, which is sufficient to activate AMPK via an elevated AMP/ATP ratio^48^. The activation of AMPKα through mtCI inhibition by C458 mirrors the effects of exercise^49^ and calorie restriction^50^, where the degree and timing of inhibition influence the mode of AMPK activation^51^. A single oral dose of C458 induced robust AMPKα activation comparable to that induced by high-intensity exercise (Fig. 3f). Chronic C458 treatment, which was administered *ad libitum* for more than 7.5 months, produced mild and sustained AMPK activation (Fig. 7e), akin to long-duration, moderate-intensity exercise. Both high- and moderate-intensity exercise preferentially activate AMPKα2^52^, suggesting that the energetic stress induced by C458 inhibition of mtCI likely also activates this isoform. Moreover, C458 may selectively activate mitochondria-associated AMPK, the mitoAMPK α2/β2/γ1 isoform, which responds first to energetic stress induced by mtCI inhibition or exercise^53^. This selective activation of mitoAMPK, particularly the regulation of mitophagy and biogenesis, is critical for mitochondrial quality control^53^. Given that the AMPKα2 isoform plays a vital role in regulating long-term synaptic plasticity and memory formation^54^, C458 may preferentially activate the mitoAMPK α2/β2/γ1 isoform in neurons, providing an additional layer of selective targeting of cells with high energy demand.

In contrast to complete mtCI inhibition by rotenone, weak mtCI inhibition by C458 does not increase ROS production or trigger inflammation. In contrast, C458 treatment restored the NADPH/GSH pools, effectively halting Aβ-mediated toxicity and preventing ferroptosis/oxytosis- mediated death in MC65 cells. It also reduces oxidative stress and chronic inflammation in APP/PS1 mice. The increase in the NADPH pool is consistent with the inhibition of ACC1 and ACC2 by AMPKα, which decreases NADPH consumption in fatty acid synthesis while increasing NADPH generation via fatty acid oxidation^55^. Additionally, AMPKα-dependent activation of the Nrf2 and AKT/FOXO3A pathways upregulated the expression of the antioxidant genes HO-1 and NQO-1, along with glutathione biosynthesis, providing an extra layer of antioxidant protection. The inactivation of both FOXO1/3A proteins and Gsk3β via PI3K/AKT activation closely correlates with AMPK activation, further supporting its critical role in the mechanisms of CP2 and C458^56^. In AD, inactivation of GSK3β offers significant neuroprotection by reducing Tau hyperphosphorylation and Aβ production^57^, whereas inactivation of FOXO1/3A prevents neuronal apoptosis, improving insulin signaling and metabolic function^58^. Downregulation of GSK3β, FOXO1A and FOXO3A activity has been shown to reduce inflammation and synaptic dysfunction^57,58^. Thus, targeting diverse pathological processes in AD can be achieved by directly modulating the adaptive mitochondrial stress response via weak inhibition of mtCI with the small molecules CP2 or C458. This approach coordinates multiple pathways to restore mitochondrial function, maintain energy homeostasis, and ultimately provide neuroprotection. Notably, the activation of adaptive mitochondrial stress responses by C458 is critically dependent on AMPKα. In AMPKα1/α2 knockout MEFs, C458 treatment fails to activate the adaptive stress response, exacerbating mitochondrial dysfunction, oxidative stress, and inflammation while preventing cellular adaptation to mild energetic stress.

Activation of AMPKα in response to mild mtCI inhibition and changes in the AMP/ATP ratio^13^, rather than direct activation of AMPKα, appears to be a promising strategy for neuroprotection. The diversity of AMPKα heterotrimeric complexes and their tissue-specific distribution present a challenge in targeting AMPKα activation specifically in the brain or neurons without causing potential adverse effects in other tissues^59^. Direct AMPKα activators act systemically and broadly, bypassing the AMP/ATP-sensing mechanism and activating AMPKα without causing true energetic stress. Systemic and chronic AMPKα activation has been associated with adverse metabolic consequences^60,61^. In contrast, mtCI inhibitors such as C458 offer a more targeted approach, preferentially activating AMPK in mitochondria-rich tissues, such as the brain^13^, suggesting that these inhibitors promote mitochondrial adaptation in select organs without significantly activating AMPKα in tissues that are less dependent on OXPHOS-driven metabolism. Additionally, some mtCI inhibitors, such as metformin^62^ and MitoTam^63^, exhibit specificity because they accumulate in the mitochondrial matrix, further enhancing their targeted action on mitochondria. While we did not test whether C458 accumulates in mitochondria, we cannot rule out this possibility on the basis of its similarity with the mechanism of CP2 action^13^. Although C458 competes with rotenone for binding to the Qd site of mtCI, it only mildly inhibits mtCI, as evidenced by modest NADH accumulation and a gradual decline in the OCR. The neuroprotective action of C458 is blocked by nanomolar concentrations of rotenone, which irreversibly bind to the Qd site of mtCI, suggesting that C458 binds to the Qd site reversibly with low affinity. Interestingly, rotenone at picomolar concentrations reversed age-related gene expression changes, rejuvenated the transcriptome, and extended the lifespan of *N. furzeri*^64^. These findings indicate that low doses of mtCI inhibitors, or weak inhibition, can activate the beneficial stress response. However, complete inhibition of mtCI by higher doses of rotenone induces ATP depletion, severe mitochondrial dysfunction, excessive ROS production, and oxidative stress, ultimately causing neurodegeneration. In contrast, C458 promotes mitochondrial adaptation without generating ROS, restoring energy balance and leading to neuroprotection.

Since AD is a multifactorial disease, various combination therapies^65^ and multitarget^66^ and dual-action^67^ small molecules have been tested to increase treatment efficacy. While these therapies can address multiple pathological mechanisms, such as Aβ accumulation and inflammation, activation of the adaptive stress response via mtCI inhibition offers a more specific and targeted therapeutic approach, which may lead to fewer side effects, greater precision, and strong efficacy in AD, where mitochondrial dysfunction is one of the earliest symptoms and a contributing factor to disease development and progression^4,68^. Given this, mtCI inhibitors could be administered as a single drug or in combination with other medications relatively early in AD progression when mitochondrial function has not yet been severely impaired. This would reduce concerns regarding the additive effect of mtCI inhibition, which could exacerbate mitochondrial dysfunction. Together with data demonstrating the strong safety of metformin in aging populations and our data showing neuroprotection after the administration of CP2 to APP/PS1 mice with reduced mitochondrial function^14,69^, our results in *Ndufs4KO* mice show that CP2 enhances mitochondrial biogenesis and turnover without causing toxicity or affecting lifespan, even when ∼50% of mitochondrial function has been lost^70^.

The dual effect of reducing p-Tau and the Aβ42/Aβ40 ratio in *APOE4*-expressing human organoids makes C458 a particularly attractive compound for addressing major hallmarks of AD in a single therapeutic. This multitarget action is especially significant in AD, where amyloid and p-Tau pathologies are interlinked and often synergistically contribute to neurodegeneration. The ability of C458 to reduce both p-Tau and Aβ levels could slow or prevent early neurodegenerative processes in *APOE4* carriers. This is critical, as *APOE4*-related neurodegeneration tends to occur early in life, and delaying these pathologies could significantly impact cognitive health. Given its ability to alleviate inflammation, increase antioxidant capacity, increase mitochondrial bioenergetics and autophagy, and promote neuroprotective responses, this study highlights the potential of mtCI inhibitors as treatments for age-related neurodegenerative diseases.

## Methods

### Ethics declaration

All experiments with mice were approved by the Mayo Clinic Institutional Animal Care and Use Committee in accordance with the National Institutes of Health’s *Guide for the Care and Use of Laboratory Animals*.___IACUC protocol number A00001186.

### Reagents

CP2 was synthesized by Nanosyn, Inc. (http://www.nanosyn.com), as described previously^16^, and was purified via HPLC. Authentication was performed through NMR spectra to ensure the lack of batch-to-batch variation in purity. CP2 was synthesized as a free base. For *in vitro* experiments, CP2 was prepared as a 10 mM stock solution in DMSO. Stock aliquots of 20 µl were stored at - 80 °C. The following reagents were used: DMSO (Sigma, D2650), hydrogen peroxide 30% solution (Sigma, H1009), MEM nonessential amino acids 100x (Corning, 25-025-CI), penicillin‒ streptomycin (Sigma, PO781), tetracycline hydrochloride (Sigma, T7660), high-glucose DMEM (Thermo Scientific, 11995065), heat-inactivated fetal bovine serum (Sigma, F4135), RPMI-1640 (Corning, MT10041CM), DPBS 1x (Corning, 21-031-CM), sodium pyruvate (Corning, MT25000CI), and phenol red-free Opti-MEM^TM^ I Reduced Serum Medium (Thermo Scientific, 51200038).

### Cells

Human neuroblastoma MC65 cells were a gift from Dr. Bryce Sopher (University of Washington, Seattle, USA). AMPKa1/a2 knockout mouse embryonic fibroblasts (MEFs) were a gift from Dr. Benoit Viollet (Inserm, Paris, France). Human neuroblastoma SH-SY5Y cells stably transfected with Swedish mutant amyloid precursor protein (APPswe) were a gift from Dr. Dr. Cristina Parrado (Karolinska Institute, Sweden). The cells were grown in high-glucose DMEM supplemented with 10% FBS, 1 mM sodium pyruvate and 1x nonessential amino acids. Primary mouse cortical neurons were cultured as described previously ^18^. Neurons from neonatal NTG animals (P1) were isolated and plated from individual pups. All experiments were performed in neurons after 7 days in culture. All the cells were incubated in 5% CO2 at 37 °C unless otherwise noted.

### Antibodies

The following primary antibodies were used: p-AMPK (Thr 172) (1:1000, Cell Signaling Technology, cat. # 2535, RRID:AB_331250), AMPK (1:1000, Cell Signaling Technology, cat. # 2532, RRID:AB_330331), p-acetyl-CoA carboxylase (Ser79) (1:1000, Cell Signaling Technology, cat. #11818, RRID:AB_2687505), p-GSK3β (Ser 9) (1:1000, Cell Signaling Technology, cat. # 9323, RRID:AB_2115201), GSK3β (1:1000, Cell Signaling Technology, cat. # 9832, RRID:AB_10839406), Sirt3 (1:1000, Cell Signaling Technology, cat. # 5490, RRID:AB_10828246), superoxide Dismutase 1 (1:1000, Abcam, cat. # ab16831, RRID:AB_302535), p-Akt (Ser473) (1:1000, Cell Signaling Technology, cat. # 4051, RRID:AB_331158), and p-Akt (Thr308)(1:1000, Cell Signaling Technology, cat. # 4056, RRID:AB_331163), p-FOXO1A (Cell Signaling Technology, cat. # 84192, RRID:AB_2800035), p- FOXO3A (Cell Signaling Technology, cat. # 9466, RRID:AB_2106674), p-ULK1 (Ser757) (1:1000, Cell Signaling Technology, cat. # 14202, RRID:AB_2665508), p-ULK1 (Ser555) (1:1000, Cell Signaling Technology, cat. # 5869, RRID:AB_10707365), p-ULK1 (Ser317) (1:1000, Cell Signaling Technology, cat. # 12753, RRID:AB_2687883), PGC1α (1∶1000, Calbiochem, cat. # KP9803); OXPHOS cocktail (Abcam, ab110413, RRID:AB_2629281), NF-kB Pathway Sampler Kit (Cell Signaling Technology, 9936T, RRID:AB_561197), p-NF-κB p65 (Ser536) (1:1000, Cell Signaling Technology, cat. # 3033, RRID:AB_331284), IκBα (1:1000, Cell Signaling Technology, cat. # 4812, RRID:AB_10694416), HO-1 (1:1000, Cell Signaling Technology, cat. # 70081, RRID:AB_2799772), mouse monoclonal NQO1 (A180) (Santa Cruz, 1:200, sc-32793, RRID:AB_628036), ND1 (USBiological, cat.# 224423), ND5 (USBiological, cat.# 224422), NDUFA9 (Thermo Fisher Scientific Cat# 459100, RRID:AB_10376187), Anti-β-Actin (1:5000, Sigma-Aldrich, cat. # A5316, RRID:AB_476743), tubulin (Rockland, cat.#200-301-880, RRID:AB_2611065) and Complex I immunocapture kit (ab109711). The following secondary antibodies were used: donkey anti-rabbit IgG conjugated with Horseradish Peroxidase (1:10000 dilution, GE Healthcare UK Limited, UK) and sheep anti-mouse IgG conjugated with Horseradish Peroxidase (1:10000 dilution,The following secondary antibodies were used: donkey anti-rabbit IgG conjugated with horseradish peroxidase (1:10000 dilution; GE Healthcare UK Limited, UK) and sheep anti-mouse IgG conjugated with horseradish peroxidase (1:10000 dilution; GE Healthcare UK Limited, UK).

### Rational design of new MTCI inhibitors

C458 was synthesized by Nanosyn, Inc. (http://www.nanosyn.com), and purified via high-performance liquid chromatography (HPLC). The authentication was performed via NMR spectroscopy and HPLC-MS to ensure the lack of batch-to-batch variation in purity (>99%). For *in vitro* experiments, C458 was prepared as a 10 mM stock solution in DMSO. Stock aliquots of 20 µl were stored at -20 °C.

### Synthesis of cis-(N-(pyridin-4-ylmethyl)-2-(3-(m-tolyloxy)cyclohexyl)propan-1-amine) (C458)

A complete step-by-step synthesis of C458 is presented in a protocol described by Trushin S. *et al.*, Synthesis of cis-(N-(pyridin-4-ylmethyl)-2-(3-(m-tolyloxy)cyclohexyl)propan-1-amine) dx.doi.org/10.17504/protocols.io.3byl4wd98vo5/v1, 2025.

### Synthesis of C458 constructs for MTCI pull-down

C458 constructs were synthesized via the C458 route, step 9, using amino alkylpyridines with 2- and 8-atom spacers attached to the NH2 group.

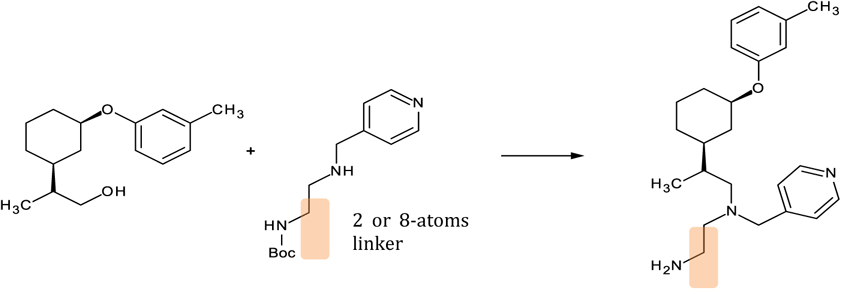

### Treatment of primary neuronal cultures with CP2 and C458

Primary mouse cortical neurons were cultured as previously described^13^. Neurons from neonatal animals (P1) were isolated and plated from individual pups; genotyping was performed prior to the day of the experiment. All experiments were performed in neurons cultured for 7 days. On day 7, the cells were treated with either C458 or CP2 at concentrations ranging from 1–20 μM for 24 hours. As a control, the cells were treated with vehicle (0.01% DMSO). Cell viability was measured after 24 hours of treatment via the MTT assay.

### Mitochondrial isolation and measurement of electron transport chain (ETC) complex activity

Intact brain mitochondria were isolated from mouse brain tissue via differential centrifugation with digitonin treatment^13^. The brain tissue was immersed in ice-cold isolation medium (225 mM mannitol, 75 mM sucrose, 20 mM HEPES-Tris, and 1 mM EGTA, pH 7.4) supplemented with 1 mg/ml BSA. The tissue was homogenized with 40 strokes by the “B” (tight) pestle of a Dounce homogenizer in 10 ml of isolation medium, diluted twofold and transferred into centrifuge tubes. The homogenate was centrifuged at 5,900 × g for 4 min in a refrigerated (4 °C) Beckman centrifuge. The supernatant was centrifuged at 12,000 × g for 10 min, the pellets were resuspended in the same buffer, and 0.02% digitonin was added. The suspension was homogenized briefly with five strokes in a loosely fitted Potter homogenizer and centrifuged again at 12,000 × g for 10 min, then gently resuspended in isolation buffer without BSA and washed once by centrifuging at 12,000 × g for 10 min. The final mitochondrial pellet was resuspended in 0.1 ml of washing buffer and stored on ice. The activity of mtCI was measured spectrophotometrically via a plate reader (SpectraMax M5, Molecular Devices, USA) in 0.2 ml of standard respiration buffer composed of 125 mM sucrose, 25 mM Tris-HCl (pH = 7.5), 0.01 mM EGTA, and 20 mM KCl at 25 °C. The NADH-dependent activity of complex I was assayed as oxidation of 0.15 mM NADH at 340 nm (ε340 nm = 6.22 mM-1 cm-1) in assay buffer supplemented with 10 µM cytochrome c, 40 µg/ml alamethicin, and 1 mM MgCl2 (NADH media). NADH:Q reductase activity was measured in NADH media containing 2 mg/ml BSA, 60 µM decylubiquinone, 1 mM cyanide and 5–15 µg protein per well. The activity of the ETC complexes was measured via Complex I (Cayman Chemicals, cat.#700930), Complex II/III (Cayman Chemicals, cat.# 700950), Complex IV (Cayman Chemicals, cat.# 700990), and Complex V (Cayman Chemicals, cat.# 701000) colorimetric assay kits.

### Pull-down of mtCI with immobilized C458 analogs

Both C458 analogs with 2-atom (458-2) or 8-atom (458-8) linkers were immobilized on Pierce NHS-activated agarose (Thermo Scientific™ Pierce™, cat. # 26197) at two concentrations, 1 mM and 200 μM, in 0.01 M borate buffer (pH 9) for one hour, followed by quenching with 1 M ethanolamine overnight. NHS-activated agarose treated with 1 M ethanolamine overnight served as control beads for nonspecific binding. mtCI was pulled down with 458-2- or 458-8-immobilized agarose beads or immunocaptured with mtCI antibodies (Ab109711, Abcam) following the manufacturer’s protocol. Briefly, freshly isolated mouse liver mitochondrial pellets were resuspended to 5.5 mg/ml in PBS containing protease inhibitors. The mitochondria were lysed with 1% n-dodecyl β-D-maltoside (Sigma, D4641) for 30 minutes on ice and then centrifuged for 10 minutes at 70,000 × g at 4 °C. For mtCI pull-down or immunocapture, the mitochondrial lysate (1 mg in 200 µl) was incubated overnight at 4 °C with 80 µl of solid beads (either 458--2 or 458-- 8) or 25 µl of immunocapturing beads. Ethanolamine-immobilized beads (25 µl) were used as a negative control. After incubation, the beads were washed three times with 1 ml of lysis buffer. After washing, the proteins were eluted from the beads by heating at 95 °C for 5 minutes in 2x Laemmli sample buffer (Bio-Rad, cat. # 1610737). Proteins were resolved by SDS‒PAGE and immunoblotted for the ND2, ND5 and NDUFA9 subunits of mtCI.

### Real-time respirometry

The kinetic injection experiment was performed via the Agilent Seahorse XFe96 Extracellular Flux Analyzer. Prior to the assay, the media was replaced with Agilent Seahorse XF DMEM, pH 7.4 (Agilent, 103575-100), supplemented with 1 mM pyruvate, 2 mM glutamine, and 10 mM glucose at 37 °C for 1 hour and placed in a BioTek Cytation 5 Cell Imaging Multimode Reader (Agilent). During the assay run, the compounds were injected via one of the ports, and the oxygen consumption rate (OCR) was measured every 18 minutes after compound injection for 1 hour.

### Mitochondrial stress test

This assay was performed to determine mitochondrial bioenergetics. Before each assay, the media was exchanged for Agilent Seahorse XF DMEM pH 7.4 (Agilent, 103575-100) supplemented with 1 mM pyruvate, 2 mM glutamine, and 10 mM glucose at 37 °C for 1 hour without CO_2_. Additionally, brightfield images were obtained via a BioTek Cytation 5 Cell Imaging Multimode Reader (Agilent). The oxygen consumption rate (OCR) was analyzed under basal conditions and after treatment with different drugs, including the ATP synthase inhibitor oligomycin A (the optimal dose chosen after the dose‒response optimization assay: 0.5–3 µM oligomycin A), an ETC accelerator ionophore (the optimal dose chosen after the dose‒response optimization assay: 0.5–3 µM FCCP), and an ETC inhibitor mixture (0.5 µM rotenone and 0.5 µM antimycin A). The response to the minimal dose of oligomycin A (2 µM) and FCCP (2 µM), generating the maximal effect, accounts for nonphosphorylating mitochondrial respiration and maximal FCCP-uncoupled respiration, respectively. The response to the rotenone and antimycin A mixture accounts for nonmitochondrial oxygen consumption. Hoechst 3342 dye (final concentration of 5 μM) was injected at the end of the assay, and the mixture was incubated at 37 °C for 15–30 min before fluorescence imaging via XF Imaging and Cell Counting Software for normalization to the cell count. The spare respiratory capacity (SRC) was calculated as the difference between the maximal and basal OCRs.

### Cell viability assays

MC65 cells were treated with 3-(4,5-dimethylthiazol-2-yl)-2,5-diphenyl tetrazolium bromide (MTT) at 0.25 mg/ml, followed by 4 hours of incubation at 37 °C. The solubilization of MTT formazan was performed by the addition of 200 µl of 40 mM HCl in isopropanol (2:1) in 100 µl of media for 10 min on a shaker, and the absorbance was measured at 570 nm. In some cases, cell viability was measured via a CellTiter-Glo® 2.0 assay according to the instructions provided by the manufacturer (Promega, WI).

### *In vitro* safety pharmacology studies

*In vitro* safety pharmacology assays, including C458 binding to human receptors and ion channels, enzyme inhibition, and uptake measurements, were conducted by the Contract Research Organization (CRO) Eurofins (France). C458 was tested at 10 μM (Supplementary Table 5). Compound binding was calculated as the % inhibition of the binding of a radioactively labeled ligand specific for each target. The compound enzyme inhibition effect was calculated as a percentage of the control enzyme activity. In each experiment and if applicable, the respective reference compound was tested concurrently with the test compound, and the data were compared with historical values determined at Eurofins. The experiment was performed in accordance with the Eurofins validation SOP. The results showing an inhibition (or stimulation for assays run under basal conditions) greater than 50% are considered to represent significant effects of the test compounds. The results showing an inhibition (or stimulation) between 25% and 50% are indicative of weak to moderate effects. The results showing an inhibition (or stimulation) lower than 25% are not considered significant and are mostly attributable to variability of the signal around the control level. Low to moderate negative values have no real meaning and are attributable to variability of the signal around the control level. High negative values (≥ 50%) that are sometimes obtained with high concentrations of test compounds are generally attributable to nonspecific effects of the test compounds in the assays. CRO Nanosyn, Inc. (Santa Barbara, CA) was contracted to conduct kinome wide panel (KWP) screening against 250 kinases via the established SOP. C458 was tested at 1 μM and 10 μM concentrations. The data are provided in Supplementary Table 1.

### *In vivo* C458 pharmacokinetic studies

**The** C458 pharmacokinetic profile was determined in C57BL/6J female mice ordered from the Jackson Laboratory. The mice were acclimatized for one week to the new environment prior to the initiation of the experiments. To evaluate C458 concentrations in plasma, mice were injected with a single intravenous dose of 3 mg/kg in 20% PEG 400 and 5% dextrose water via the lateral tail vein via a 0.5 cc tuberculin syringe. Independent cohorts of mice were administered C458 by oral gavage (25 mg/kg in 20% PEG 400 and 5% dextrose water) with a 1 cc tuberculin syringe with a stainless steel 22-gauge straight feeding needle. At 0, 0.08, 0.25, 0.5, 1, 2, 4, 8 and 24 hours after treatment, the mice were anesthetized, and 200 µl of blood was collected from the retro-orbital sinus through a K_2_EDTA-coated capillary into a K_2_EDTA-coated microtainer tube. The plasma was separated via centrifugation at 4 °C (10,000 rpm for 3 min), transferred to a microcentrifuge tube, immediately frozen on dry ice, and stored at -80 °C until analysis. The pharmacokinetic parameters of C458 were estimated via standard noncompartmental analysis.

### C458 quantification via LC‒MS/MS

The method used to quantify C458 in plasma and brain tissue is described in the protocol: Trushin S. *et al*, C458 quantification using LC‒MS/MS, dx.doi.org/10.17504/protocols.io.eq2ly62xmgx9/v1, 2025.

### Studies in mice

Double transgenic APP/PS1 mice were used in the study^14^. The genotypes were determined via PCR as described previously ^14^. All the animals were maintained on a 12–12 h light‒12 h dark cycle, with a regular feeding and cage-cleaning schedule. The mice were randomly assigned to study groups on the basis of their age and genotype. Two trials were conducted within a few months to ensure that mice of the same age and sex were consistently treated. The first trial included three male and three female mice in the vehicle-treated group and four male and three female mice in the C458-treated group. The second trial included five male and seven female mice in the vehicle-treated group and eight female and two male mice in the C458-treated group. Data analysis was conducted on samples combined from both trials. Where possible, analyses were conducted by the investigators blinded to the treatment.

### Chronic C458 treatment in presymptomatic APP/PS1 mice

APP/PS1 male and female mice were given C458 (25 mg/kg/day in 0.05% DMSO dissolved in drinking water *ad libitum*) or vehicle-containing water (0.05% DMSO) starting at 2.5 months of age as we described previously^14^. Water consumption and weight were monitored weekly. Independent groups of mice were continuously treated for 7 months until the age of 10 months. Prior to the beginning of treatment, the mice were subjected to a battery of behavior tests. At the end of the treatment, the mice were subjected to a behavior battery, metabolic cages (CLAMS) and ^31^P-NMR imaging. After the mice were sacrificed, tissue and blood were collected from each mouse. The brain tissues were used for western blot analysis, and the plasma was used for cytokine/chemokine profiling. Three mice from each group were subjected to electrophysiology recording as described below. One female mouse treated with C458 lost 20% of its body weight during the last two weeks of the study. Since this mouse did not demonstrate any signs of motor or behavioral distress, it was not removed from the study.

### Behavior battery

Behavioral tests were carried out in the light phase of the circadian cycle, with at least 24 h between each assessment, as we described previously^14^. More than one paradigm ran within 1 week. However, no more than two separate tests were run on the same day. Behavioral and metabolic tests were performed in the order described in the experimental timeline.

#### Open field test

Spontaneous locomotor activity was measured in brightly lit (500 lux) Plexiglas chambers (41 cm × 41 cm) that automatically recorded the activity via photobeam breaks (Med Associates, Lafayette, IN). The chambers were located in sound-attenuating cubicles and were equipped with two sets of 16 pulse-modulated infrared photobeams to automatically record X–Y ambulatory movements at a 100 ms resolution. Data were collected over a 10-minute trial with 30 second intervals.

#### Hanging bar

Balance and general motor function were assessed using a hanging bar. The mice were lowered onto a parallel rod (D < 0.25 cm) placed 30 cm above a padded surface. The mice were allowed to grab the rod with their forelimbs, after which they were released and scored for success (pass or failure) in holding onto the bar for 30 seconds. The test consisted of a 3-day trial period with three 30-second measurements taken each day. The final latency to fall in sec was presented as an average of nine tests per animal.

#### A novel object recognition (NOR) test

was used to estimate memory deficits. All trials were conducted in an isolated room with dim light in Plexiglas boxes (40 cm × 30 cm). Each mouse was placed in a box for 5 min for the acclimatization period. Thereafter, a mouse was removed, and two similar objects were placed in the box. Objects with various geometric shapes and colors were used in the study. A mouse was returned to the box, and the number of interrogations of each object was automatically recorded by a camera placed above the box for a duration of 10 min. A mouse was removed from the box for 5 min, and one familiar object was replaced with a novel object. Each mouse was returned to the box, and the number of interrogations of novel and familiar objects was recorded for 10 min. Experiments were analyzed via NoldusЕthoVision software. The number of interrogations of the novel object was divided by the number of investigations of the familiar object to generate a discrimination index. Intact recognition memory produces a discrimination index of 1 for the training session and a discrimination index greater than 1 for the test session, which is consistent with greater interrogation of the novel object.

#### Morris water maze

Spatial learning and memory were investigated by measuring the time it took each mouse to locate a platform in opaque water identified with a visual cue above the platform. The path taken to the platform was recorded with a camera attached above the pool. Each mouse was trained to find the platform during four training sessions per day for three consecutive days. For each training session, each mouse was placed in the water facing away from the platform and allowed to swim for up to 60 s to find the platform. Each training session started by placing a mouse in a different quadrant of the tank. If the mouse found the platform before 60 s had passed, the mouse was left on the platform for 30 s before being returned to its cage. If the animal had not found the platform within 60 s, the mouse was manually placed on the platform and left there for 30 s before being returned to its cage. A day of rest was followed by the day of formal testing.

### Inflammatory markers

After 16 hours of fasting, blood from APP/PS1 mice treated with vehicle or C458 was collected via orbital bleeding and centrifuged for 5 min at 5000 × g. Collected plasma was sent for 32-plex cytokine array analysis (Discovery Assay, Eve Technologies Corp. https://www.evetechnologies.com/discovery-assay/). Multiplexing analysis was performed via the Luminex 100 system (Luminex). More than 80% of the targets were within the detectable range (signals from all samples were higher than the lowest standard). Undetectable targets were excluded. Blood samples were run in duplicate.

### Lipid peroxidation assay

The levels of malondialdehyde (MDA), a product of lipid degradation that occurs as a result of oxidative stress, were measured via an MDA assay kit (#MAK085, Sigma Aldrich) in hippocampal brain tissue isolated from APP/PS1 mice treated with vehicle or C458 according to the manufacturer’s instructions.

### Electrophysiology

APP/PS1 mice aged ∼7 months and treated with C458 (*n* = 3 *per* group) or vehicle (*n* = 3 *per* group) were used for electrophysiology analysis. The mice were deeply anesthetized with isoflurane and decapitated. The brain was quickly removed and transferred to cold slicing solution containing artificial cerebrospinal fluid (ACSF), where NaCl was substituted with sucrose to avoid excitotoxicity. Transverse slices (300–350 µm thick) were made via a vibratome (VT-100S, Leica). Slices were incubated in ACSF containing 128 mM NaCl, 2.5 mM KCl, 1.25 mM NaH_2_PO_4_, 26 mM NaHCO_3_, 10 mM glucose, 2 mM CaCl_2_, and 1 mM MgSO_4_ aerated with 95% O_2_/5% CO_2_. Slices were maintained at 32 °C for 13 min and then maintained at room temperature throughout the entire experiment. For electrophysiology recording, each slice (2–3 slices *per* mouse) was transferred to a recording chamber, and ACSF was continuously perfused at a flow rate of 2–3 ml/min. A single recording electrode and a single bipolar stimulation electrode were placed on top of the slice. A boron-doped glass capillary (PG10150, World Precision Instruments) was pulled with a horizontal puller (P-1000, Sutter Instrument) and filled with ACSF for extracellular recording. Under a microscope (FN-1, Nikon), the recording electrode was placed in the CA1 area of the hippocampus. The bipolar stimulation electrode (FHC) was placed at Schaffer collaterals. The distance between the two electrodes was greater than ∼200 µm. To define the half response of the stimulation, various intensities of electrical stimulation were applied (10–500 µA). However, the pulse width was fixed at 60 μs. Once the stimulation parameter was determined to generate half the maximum evoked fEPSP, this stimulation intensity was used for paired-pulse and LTP experiments. For the LTP experiment, test stimulation was applied every 30 sec for 30 min to achieve a stable baseline. Once a stable baseline was achieved, tetanic stimulation (100 Hz for 1 second) was applied three times with 3-second intervals. The initial slope of the fEPSP was used to compare the synaptic strength.

### Comprehensive Laboratory Animal Monitoring System (CLAMS)

CLAMS (Columbus Instruments, Columbus, OH) is a set up allowing automated, noninvasive and simultaneous monitoring of horizontal and vertical activity, feeding and drinking, oxygen consumption and CO_2_ production of an individual mouse. APP/PS1 mice treated with vehicle or C458 (9–10 months old) were individually placed in CLAMS cages. Indirect calorimetry was monitored over 2 days, where the mice were allowed food for 24 hours *ad libitum* (fed state), and for the following 24 hours, the food was removed (fasting state). The mice were maintained at 20–22 °C under a 12:12 hr light–dark cycle. All the mice were acclimatized to CLAMS cages for 3–6 hours before recording. Sample air was passed through an oxygen sensor for determination of the oxygen content. Oxygen consumption was determined by measuring the oxygen concentration in the air entering the chamber compared with that in the air leaving the chamber. The sensor was calibrated against a standard gas mixture containing defined quantities of oxygen, carbon dioxide and nitrogen. Food and water consumption were measured directly. The hourly file displayed measurements for the following parameters: VO_2_ (volume of oxygen consumed, mL/kg/h), VCO_2_ (volume of carbon dioxide produced, mL/kg/h), RER (respiratory exchange ratio), heat (Kcal/h), total energy expenditure (TEE, kcal/h/kg of lean mass), activity energy expenditure (AEE, kcal/h/kg of lean mass), resting total consumed food (REE kcal/h/kg of lean mass), food intake (g/kg of body weight/12 hours), metabolic rate (kcal/h/kg), total activity (all horizontal beam breaks in counts), ambulatory activity (minimum 3 different, consecutive horizontal beam breaks in counts), and rearing activity (all vertical beam breaks in counts). EE, RER, and fatty acid (FA) oxidation were calculated via the following equations: RER = VCO2/VO2; EE(kcal/h) = [3.815+1.232 × RER] × VO2] × 1000; and FA oxidation (kcal/h) = EE X (1-RER/0.3). Daily FA oxidation was calculated from the average 12 hours of hourly FA oxidation. Daily carbohydrate plus protein oxidation was calculated from the average 12-hour hourly EE minus daily FA oxidation. Metabolic flexibility was evaluated from the difference in RER between the daily fed state and fasted state recorded at night according to the following equation: Δ=100% * (RER fed-RER fasted)/RER fed.

### Dual-energy X-ray absorptiometry (DEXA)

The LUNAR PIXImus mouse densitometer (GE Lunar, Madison, WI), a dual-energy supply X-ray machine, was used for measuring skeletal and soft tissue masses for the assessment of skeletal and body composition in C458- or vehicle-treated mice. Live mice were scanned under 1.5–2% isoflurane anesthesia. The mice were individually placed on plastic trays, which were then placed onto the exposure platform of the PIXImus machine to measure body composition. The following parameters were generated: lean mass (in grams), fat mass (in grams), and the percentage of fat mass. These parameters were used to normalize the data generated in CLAMS, including O_2_, VCO_2_, metabolic rate and energy expenditure.

### *In vivo* ^31^P-NMR Spectroscopy

NMR spectra were obtained from 10-month-old APP/PS1 mice treated with C458 (*n* = 6 *per* group) or vehicle for ∼7 months (*n* = 7 *per* group) via an Avance III 300/700 MHz (7/16.4 T) wide-bore NMR spectrometer equipped with microimaging accessories (Bruker, Billerica, MA, USA) with a 25-mm inner diameter dual nucleus (^31^P/^1^H) birdcage coil. For anatomical positioning, a pilot image set of the coronal, sagittal, and axial imaging planes was used. For ^31^P spectroscopy studies, a single pulse acquisition with a pulse width of P1 200 μs (∼30 degrees), spectral width of SW 160 ppm, FID size of TD 16k, FID duration of AQT 0.41 s, waiting time of D1 1 second and number of scans of NS 512 was used. The acquisition time was 12 min. Spectra were processed via TopSpin v3.5 software (BrukerBiospin MRI, Billerica, MA). The integral areas of the spectral peaks corresponding to inorganic phosphate (Pi), phosphocreatine (PCr), and the γ, α, and β phosphates of adenosine triphosphate (αATP, βATP, and γATP) were measured. Since the Pi peaks were not detectable in some mice, the Pi values were uniformly omitted from the analyses. The presence of phosphomonoester (PME) or phosphodiester (PDE) peaks was also recorded. However, the signal‒to‒noise ratios of these peaks are not always adequate for accurate quantification. The levels of PCr and Pi were normalized by the total ATP levels present in that spectrum or by the amount of βATP. The results were consistent between both normalization methods; the data are presented as the ratio of each parameter to total ATP.

### Differentiation of iPSCs into cortical organoids

Cortical organoids were generated via the use of the STEMdiff™ Cerebral Organoid Kit (Stemcell Technologies) according to the manufacturer’s instructions, with slight modifications. On day 0, the iPSCs were dissociated into single-cell suspensions with TrypLE Express (Thermo Fisher Scientific, Waltham, MA, USA) and cultured on U-bottom ultralow-attachment 96-well plates (15,000 cells/well) in embryoid body (EB) formation media (medium A) supplemented with 10 μM Y27632. After the medium was changed to EB formation media on days 2 and 4, the iPSC-derived EBs were transferred onto 96-well low-attachment plates on day 5 and cultured in induction medium (medium B). On day 7, the EBs were transferred onto Matrigel-coated 6-well plates and cultured in expansion medium (medium C + D) for cortical organoid formation for 3 days. On day 10, the culture medium was changed to maturation medium (medium E). After 4 weeks, medium E was replaced with neuronal maturation medium, which was composed of DMEM/F12 + neurobasal medium (1:1) supplemented with N2, B27, BDNF (20 ng/ml), GDNF (20 ng/ml), ascorbic acid (200 μM), and dbcAMP (100 nM) (Sigma‒Aldrich). The organoids were cultured on an orbital shaker, and the medium was changed twice per week until day 60.

### Organoid processing

Organoids were collected 48 hours after treatment with compounds and lysed in RIPA lysis buffer supplemented with protease and phosphatase inhibitor cocktails (Roche). The lysed samples were sonicated at 40% amplitude. The samples were subsequently centrifuged at 21,000 × g for 45 minutes at 4 °C. The total protein concentration in the soluble fraction was quantified via the Pierce BCA Protein Assay Kit.

### ELISA quantification

The levels of Aβ40, Aβ42, total tau, and phospho-Tau were measured via the following ELISA kits: the Human β-Amyloid (1–40) ELISA Kit (Thermo Fisher, KHB3481), the Human β-Amyloid (1–42) ELISA Kit (Thermo Fisher, KHB3544), the Human Tau ELISA Kit (Thermo Fisher, KHB0041), and the Tau (Phospho) [pT181] Human ELISA Kit (Thermo Fisher, KHO0631), according to the manufacturers’ instructions. The samples and detection antibodies were added to the ELISA plates and incubated for the specified duration of time. Following incubation, the wells were washed four times with wash buffer and incubated with an IgG-HRP conjugate for the indicated time. After additional washes, the plates were treated with stabilized chromogen and incubated for 30 minutes at room temperature in the dark. The reaction was stopped with a stop solution, and the absorbance was measured at 450 nm via a microplate reader (BioTek). The results were normalized to the total protein concentration of the cell lysate.

### Western blot analysis

Protein levels in the cortico-hippocampal region of the brains of vehicle- or C458-treated APP/PS1 mice were determined via Western blot analysis. The tissue was homogenized and lysed via RIPA buffer (25 mM Tris-HCl pH 7.6, 150 mM NaCl, 1% NP-40, 1% sodium deoxycholate, 0.1% SDS) containing phosphatase PhosSTOP (Roche, cat. #04906837001) and protease inhibitors (cOmplete, Roche, cat. #11697498001). Total protein lysates (25 μg) were separated on 4–20% Mini-PROTEAN TGX™ Precast Protein Gels (Bio-Rad, 4561093). For the C458 time course study, coronal brain slices that encompassed the cortico-hippocampal region were homogenized, and 30 µg of protein lysates were separated on 4–15% Criterion gels (Bio-Rad, cat. # 5678083) and transferred to Immun-Blot polyvinylidene difluoride membranes (PVDF cat. # 1620177). Total cell lysates were prepared using RIPA buffer. Bands were imaged and quantified via a KwikQuant imager and KwikQuant image analyzer 5.9 (Kindle Biosciences, USA).

### Statistics

All the statistical analyses were performed via GraphPad Prism. The statistical analysis included two-tailed unpaired and paired Student’s t tests (where appropriate) and one-way ANOVA. When *P* values were significant at a level of *P* < 0.05, Fisher’s LSD *post hoc* analysis was applied to determine the differences among groups. The data are presented as the means ± SDs for each group of mice.

## Contributors

E.T. conceived the study, assembled the multidisciplinary team of collaborators, and received funding for the project. S.T., A.S., S.I.M., T.K.O.N., M.O., L.Z. and J.T.D. performed the experiments and analyzed and interpreted the data. A.S. and S.Y.C. conducted the electrophysiology experiments. T.K.O.N., T.N., W.L. and T.K. conducted the experiments on the human organoids, and S.T. and E.T. wrote the manuscript. All the authors edited the manuscript and approved its publication. E.T. and S.T. directly assessed and verified the underlying data presented in the manuscript.

## Acknowledgments

We thank Mayo Clinic Cores for help with ^31^P-NMR and CLAMS data acquisition. We thank Mr. B. Gateno for help with mouse colonies; Ms. R. A. Schoon, Drs. R. A. Kudgus and J. M. Reid for help with PK/PD data acquisition; Drs. K. Greenman, W. Thomas, R. Greenhouse, and O. Isaakova for assisting with compound synthesis; Dr. Z. Liu for assisting with the experiments; Dr. M. Schellenberg for valuable suggestions; and Ms. S. Gochnauer for assisting with manuscript preparation.

## Data Sharing

All data regarding the experiments described in the paper are available upon request to the corresponding author. The C458 structure was deposited in PubChem with CID 134259587.

## Supplementary Figure Legends

**Supplementary Figure 1. Effect of C458 treatment on the activity of complexes II-V, cellular respiration and ATP production. a-c** C458 does not inhibit the enzymatic activity of succinate: cytochrome c reductase (complexes II--III), ferrocytochrome c oxidase (complex IV only), and ATP synthase (complex V). **d** CP2 and C458 decreased ATP levels in primary mouse cortical neurons after 24 hours of treatment. **e** The extracellular acidification rate (ECAR) was measured for 120 min in MC65 Tet-On cells following the injection of different doses of C458 or rotenone. **f** Quantification of the OCR data presented in Fig. 2f. **g** Viability of MC65 Tet-On cells treated with 25 nM or 1 mM rotenone for 72 hours. **h** OCR was measured for 120 min in MC65 Tet-On cells following the injection of different doses of rotenone. The data are the means ± SD. * *P*< 0.05; ** *P* < 0.01; *** *P* < 0.001, **** *P* <0.0001.

**Supplementary Figure 2. C458 analogs protect against Aβ toxicity and bind mtCI. a** Efficacy of C458-2 and C458-8 against Aβ toxicity in MC65 cells. The experiments were performed in triplicate in three independent cultures. **b** mtCI was pulled down from 1 mg of mitochondrial lysate using C458-8 and C458-2 immobilized on agarose beads. Both C458-2 and C458-8 were immobilized on NHS-activated agarose at two concentrations: 1 mM and 0.2 mM. The input was 50 μg of mitochondrial lysate; IC represents the immunocapture of mtCI from 1 mg of mitochondrial lysate with mtCI antibodies (Abcam). mtCI was detected by immunoblotting for the NDUFA9 and ND5 subunits.

**Supplementary Figure 3.** Quantification of the proteins presented on the Western blot in Fig. 7a.

**Supplementary Figure 4.** Quantification of the proteins presented on the Western blot in Fig. 7b.

**Supplementary Figure 5.** Quantification of proteins presented on the Western blot in Fig. 7e

**Supplementary Figure 6.** Original uncropped Western blot images for Fig. 3f.

**Supplementary Figure 7.** Original uncropped Western blot images for Fig. 7a.

**Supplementary Figure 8.** Original uncropped Western blot images for Fig. 7b.

**Supplementary Figure 9.** Original uncropped Western blot images for Fig. 7e.

**Supplementary Figure 10. a** Schematic overview of the procedures for generating human iPSC-derived organoids from AD patients. **b-e** Levels of extracellular Aβ40 (b), Aβ42 (c), intracellular Aβ40 (d) and Aβ42 (e) were measured via ELISA in vehicle- or C458-treated iPSC-derived organoids from AD patients. **f-i** Levels of extracellular Aβ40 (f), Aβ42 (g), intracellular Aβ40 (h) and Aβ42 (i) in vehicle- or CP2-treated iPSC-derived organoids from AD patients were measured via ELISA. **j‒m** Levels of total tau and pTau T181 in the RIPA lysates of 3 cerebral organoids treated with C458 (j‒k) or CP2 (l‒m) were analyzed via ELISA. Lysates of 3 cerebral organoids per iPSC line were analyzed at week 12 in culture. The data are the means ± SD. * *P*< 0.05; ** *P* < 0.01; *** *P* < 0.001; **** *P* <0.0001; ns, not significant.

**Supplementary Table 1. C458 Kinome profiling results.**

**Supplementary Table 2. Permeability assessment of MDR1 (P-gp)-transfected Readyport MDCKII cells for the C458 compound.**

**Supplementary Table 3. Histology results for C57BL/6 adult and newborn mice treated with C458**

**Supplementary Table 4. C458 concentration in the organs of APP/PS1 mice.**

**Supplementary Table 5. Cerep safety screen 44 panel for C458 treatment (10 mM).**

## References

1. Monzel AS, Enriquez JA, Picard M. Multifaceted mitochondria: moving mitochondrial science beyond function and dysfunction. Nat Metab 2023; 5(4): 546–62.

2. Poor TA, Chandel NS. SnapShot: Mitochondrial signaling. Mol Cell 2023; 83(6): 1012–e1.

3. Chen W, Zhao H, Li Y. Mitochondrial dynamics in health and disease: mechanisms and potential targets. Signal Transduct Target Ther 2023; 8(1): 333.

4. Cunnane SC, Trushina E, Morland C, et al. Brain energy rescue: an emerging therapeutic concept for neurodegenerative disorders of ageing. Nat Rev Drug Discov 2020; 19(9): 609–33.

5. Tonnies E, Trushina E. Oxidative Stress, Synaptic Dysfunction, and Alzheimer’s Disease. J Alzheimers Dis 2017; 57(4): 1105–21.

6. Feng J, Bussiere F, Hekimi S. Mitochondrial electron transport is a key determinant of life span in Caenorhabditis elegans. Developmental cell 2001; 1(5): 633–44.

7. Kayser EB, Sedensky MM, Morgan PG, Hoppel CL. Mitochondrial oxidative phosphorylation is defective in the long-lived mutant clk-1. J Biol Chem 2004; 279(52): 54479–86.

8. Dillin A, Hsu AL, Arantes-Oliveira N, et al. Rates of behavior and aging specified by mitochondrial function during development. Science 2002; 298(5602): 2398–401.

9. Salminen A, Kaarniranta K. AMP-activated protein kinase (AMPK) controls the aging process via an integrated signaling network. Ageing Res Rev 2012; 11(2): 230–41.

10. Cameron AR, Logie L, Patel K, et al. Metformin selectively targets redox control of complex I energy transduction. Redox Biol 2018; 14: 187–97.

11. Wang Y, An H, Liu T, et al. Metformin Improves Mitochondrial Respiratory Activity through Activation of AMPK. Cell Rep 2019; 29(6): 1511–23 e5.

12. Effect of intensive blood-glucose control with metformin on complications in overweight patients with type 2 diabetes (UKPDS 34). UK Prospective Diabetes Study (UKPDS) Group. Lancet 1998; 352(9131): 854–65.

13. Zhang L, Zhang S, Maezawa I, et al. Modulation of mitochondrial complex I activity averts cognitive decline in multiple animal models of familial Alzheimer’s Disease. EBioMedicine 2015; 2(4): 294–305.

14. Stojakovic A, Trushin S, Sheu A, et al. Partial inhibition of mitochondrial complex I ameliorates Alzheimer’s disease pathology and cognition in APP/PS1 female mice. Commun Biol 2021; 4(1): 61.

15. Stojakovic A, Chang SY, Nesbitt J, et al. Partial Inhibition of Mitochondrial Complex I Reduces Tau Pathology and Improves Energy Homeostasis and Synaptic Function in 3xTg-AD Mice. J Alzheimers Dis 2021; 79(1): 335–53.

16. Rana S, Hong HS, Barrigan L, Jin LW, Hua DH. Syntheses of tricyclic pyrones and pyridinones and protection of Abeta-peptide induced MC65 neuronal cell death. Bioorg Med Chem Lett 2009; 19(3): 670–4.

17. Panes J, Nguyen TKO, Gao H, et al. Partial Inhibition of Complex I Restores Mitochondrial Morphology and Mitochondria-ER Communication in Hippocampus of APP/PS1 Mice. Cells 2023; 12(8).

18. Trushina E, Trushin S, Hasan MF. Mitochondrial complex I as a therapeutic target for Alzheimer’s disease. Acta Pharm Sin B 2022; 12(2): 483–95.

19. Trushina E, Nguyen TKO, Trushin S. Modulation of Mitochondrial Function as a Therapeutic Strategy for Neurodegenerative Diseases. J Prev Alzheimers Dis 2023; 10(4): 675–85.

20. Sopher BL, Fukuchi K, Smith AC, Leppig KA, Furlong CE, Martin GM. Cytotoxicity mediated by conditional expression of a carboxyl-terminal derivative of the beta-amyloid precursor protein. Brain Res Mol Brain Res 1994; 26(1-2): 207–17.

21. Huang L, McClatchy DB, Maher P, et al. Intracellular amyloid toxicity induces oxytosis/ferroptosis regulated cell death. Cell Death Dis 2020; 11(10): 828.

22. Maezawa I, Hong HS, Wu HC, et al. A novel tricyclic pyrone compound ameliorates cell death associated with intracellular amyloid-beta oligomeric complexes. J Neurochem 2006; 98(1): 57–67.

23. Degli Esposti M. Inhibitors of NADH-ubiquinone reductase: an overview. Biochim Biophys Acta 1998; 1364(2): 222–35.

24. Vercellino I, Sazanov LA. Structure and assembly of the mammalian mitochondrial supercomplex CIII(2)CIV. Nature 2021; 598(7880): 364–7.

25. Kampjut D, Sazanov LA. Structure of respiratory complex I - An emerging blueprint for the mechanism. Curr Opin Struct Biol 2022; 74: 102350.

26. Fullerton MD, Galic S, Marcinko K, et al. Single phosphorylation sites in Acc1 and Acc2 regulate lipid homeostasis and the insulin-sensitizing effects of metformin. Nat Med 2013; 19(12): 1649–54.

27. Cui D, Liu H, Cao L, et al. MST1, a novel therapeutic target for Alzheimer’s disease, regulates mitochondrial homeostasis by mediating mitochondrial DNA transcription and the PI3K-Akt-ROS pathway. J Transl Med 2024; 22(1): 1056.

28. Terry RD, Masliah E, Salmon DP, et al. Physical basis of cognitive alterations in Alzheimer’s disease: synapse loss is the major correlate of cognitive impairment. Ann Neurol 1991; 30(4): 572–80.

29. de Wilde MC, Overk CR, Sijben JW, Masliah E. Meta-analysis of synaptic pathology in Alzheimer’s disease reveals selective molecular vesicular machinery vulnerability. Alzheimers Dement 2016; 12(6): 633–44.

30. Goodpaster BH, Sparks LM. Metabolic Flexibility in Health and Disease. Cell Metab 2017; 25(5): 1027–36.

31. Jett S, Boneu C, Zarate C, et al. Systematic review of (31)P-magnetic resonance spectroscopy studies of brain high energy phosphates and membrane phospholipids in aging and Alzheimer’s disease. Front Aging Neurosci 2023; 15: 1183228.

32. Cheignon C, Tomas M, Bonnefont-Rousselot D, Faller P, Hureau C, Collin F. Oxidative stress and the amyloid beta peptide in Alzheimer’s disease. Redox Biol 2018; 14: 450–64.

33. Hothersall JS, Gordge M, Noronha-Dutra AA. Inhibition of NADPH supply by 6- aminonicotinamide: effect on glutathione, nitric oxide and superoxide in J774 cells. FEBS Lett 1998; 434(1- 2): 97–100.

34. Ghosh D, Levault KR, Brewer GJ. Relative importance of redox buffers GSH and NAD(P)H in age-related neurodegeneration and Alzheimer disease-like mouse neurons. Aging Cell 2014; 13(4): 631–40.

35. Woltjer RL, Nghiem W, Maezawa I, et al. Role of glutathione in intracellular amyloid-alpha precursor protein/carboxy-terminal fragment aggregation and associated cytotoxicity. J Neurochem 2005; 93(4): 1047–56.

36. Pratico D, Uryu K, Leight S, Trojanoswki JQ, Lee VM. Increased lipid peroxidation precedes amyloid plaque formation in an animal model of Alzheimer amyloidosis. J Neurosci 2001; 21(12): 4183–7.

37. Currais A, Quehenberger O, A MA, Daugherty D, Maher P, Schubert D. Amyloid proteotoxicity initiates an inflammatory response blocked by cannabinoids. NPJ Aging Mech Dis 2016; 2: 16012.

38. Morgan MJ, Liu ZG. Crosstalk of reactive oxygen species and NF-kappaB signaling. Cell Res 2011; 21(1): 103–15.

39. Laderoute KR, Amin K, Calaoagan JM, et al. 5’-AMP-activated protein kinase (AMPK) is induced by low-oxygen and glucose deprivation conditions found in solid-tumor microenvironments. Mol Cell Biol 2006; 26(14): 5336–47.

40. Morizane Y, Thanos A, Takeuchi K, et al. AMP-activated protein kinase suppresses matrix metalloproteinase-9 expression in mouse embryonic fibroblasts. J Biol Chem 2011; 286(18): 16030–8.

41. Belyaev ND, Kellett KA, Beckett C, et al. The transcriptionally active amyloid precursor protein (APP) intracellular domain is preferentially produced from the 695 isoform of APP in a beta-secretase-dependent pathway. J Biol Chem 2010; 285(53): 41443–54.

42. Zhao J, Fu Y, Yamazaki Y, et al. APOE4 exacerbates synapse loss and neurodegeneration in Alzheimer’s disease patient iPSC-derived cerebral organoids. Nat Commun 2020; 11(1): 5540.

43. Talwar P, Sinha J, Grover S, et al. Dissecting Complex and Multifactorial Nature of Alzheimer’s Disease Pathogenesis: a Clinical, Genomic, and Systems Biology Perspective. Mol Neurobiol 2016; 53(7): 4833–64.

44. Deming Y, Dumitrescu L, Barnes LL, et al. Sex-specific genetic predictors of Alzheimer’s disease biomarkers. Acta Neuropathol 2018.

45. van Dyck CH, Swanson CJ, Aisen P, et al. Lecanemab in Early Alzheimer’s Disease. N Engl J Med 2023; 388(1): 9–21.

46. Kolata GB. The phenformin ban: is the drug an imminent hazard? Science 1979; 203(4385): 1094–6.

47. Yap TA, Daver N, Mahendra M, et al. Complex I inhibitor of oxidative phosphorylation in advanced solid tumors and acute myeloid leukemia: phase I trials. Nat Med 2023; 29(1): 115–26.

48. Krebs H. The Croonian Lecture, 1963. Gluconeogenesis. Proc R Soc Lond B Biol Sci 1964; 159: 545–64.

49. Richter EA, Ruderman NB. AMPK and the biochemistry of exercise: implications for human health and disease. Biochem J 2009; 418(2): 261–75.

50. Green CL, Lamming DW, Fontana L. Molecular mechanisms of dietary restriction promoting health and longevity. Nat Rev Mol Cell Biol 2022; 23(1): 56–73.

51. Gurd BJ, Menezes ES, Arhen BB, Islam H. Impacts of altered exercise volume, intensity, and duration on the activation of AMPK and CaMKII and increases in PGC-1alpha mRNA. Semin Cell Dev Biol 2023; 143: 17–27.

52. Sriwijitkamol A, Coletta DK, Wajcberg E, et al. Effect of acute exercise on AMPK signaling in skeletal muscle of subjects with type 2 diabetes: a time-course and dose-response study. Diabetes 2007; 56(3): 836–48.

53. Drake JC, Wilson RJ, Laker RC, et al. Mitochondria-localized AMPK responds to local energetics and contributes to exercise and energetic stress-induced mitophagy. Proc Natl Acad Sci U S A 2021; 118(37).

54. Yang W, Zhou X, Zimmermann HR, Ma T. Brain-specific suppression of AMPKalpha2 isoform impairs cognition and hippocampal LTP by PERK-mediated eIF2alpha phosphorylation. Mol Psychiatry 2021; 26(6): 1880–97.

55. Jeon SM, Chandel NS, Hay N. AMPK regulates NADPH homeostasis to promote tumour cell survival during energy stress. Nature 2012; 485(7400): 661–5.

56. Han F, Li CF, Cai Z, et al. The critical role of AMPK in driving Akt activation under stress, tumorigenesis and drug resistance. Nat Commun 2018; 9(1): 4728.

57. Lauretti E, Dincer O, Pratico D. Glycogen synthase kinase-3 signaling in Alzheimer’s disease. Biochim Biophys Acta Mol Cell Res 2020; 1867(5): 118664.

58. Lee S, Dong HH. FoxO integration of insulin signaling with glucose and lipid metabolism. J Endocrinol 2017; 233(2): R67–R79.

59. Olivier S, Foretz M, Viollet B. Promise and challenges for direct small molecule AMPK activators. Biochem Pharmacol 2018; 153: 147–58.

60. Myers RW, Guan HP, Ehrhart J, et al. Systemic pan-AMPK activator MK-8722 improves glucose homeostasis but induces cardiac hypertrophy. Science 2017; 357(6350): 507–11.

61. Yavari A, Stocker CJ, Ghaffari S, et al. Chronic Activation of gamma2 AMPK Induces Obesity and Reduces beta Cell Function. Cell Metab 2016; 23(5): 821–36.

62. Bridges HR, Jones AJ, Pollak MN, Hirst J. Effects of metformin and other biguanides on oxidative phosphorylation in mitochondria. Biochem J 2014; 462(3): 475–87.

63. Rohlenova K, Sachaphibulkij K, Stursa J, et al. Selective Disruption of Respiratory Supercomplexes as a New Strategy to Suppress Her2(high) Breast Cancer. Antioxid Redox Signal 2017; 26(2): 84–103.

64. Baumgart M, Priebe S, Groth M, et al. Longitudinal RNA-Seq Analysis of Vertebrate Aging Identifies Mitochondrial Complex I as a Small-Molecule-Sensitive Modifier of Lifespan. Cell Syst 2016; 2(2): 122–32.

65. Cummings JL, Osse AML, Kinney JW, Cammann D, Chen J. Alzheimer’s Disease: Combination Therapies and Clinical Trials for Combination Therapy Development. CNS Drugs 2024; 38(8): 613–24.

66. Guo Q, Wu G, Huang F, et al. Novel small molecular compound 2JY-OBZ4 alleviates AD pathology in cell models via regulating multiple targets. Aging (Albany NY) 2022; 14(19): 8077–94.

67. Al Assi A, Posty S, Lamarche F, et al. A novel inhibitor of the mitochondrial respiratory complex I with uncoupling properties exerts potent antitumor activity. Cell Death Dis 2024; 15(5): 311.

68. Zhang XX, Wei M, Wang HR, Hu YZ, Sun HM, Jia JJ. Mitochondrial dysfunction gene expression, DNA methylation, and inflammatory cytokines interaction activate Alzheimer’s disease: a multi-omics Mendelian randomization study. J Transl Med 2024; 22(1): 893.

69. Trushina E, Nemutlu E, Zhang S, et al. Defects in Mitochondrial Dynamics and Metabolomic Signatures of Evolving Energetic Stress in Mouse Models of Familial Alzheimer’s Disease. PLoS One 2012; 7(2).

70. Gao H, Jensen, K., Nesbitt, J., Ostroot, M., Baloni, P., Funk, C., Trushina, E. Ndufs4 knockout induces transcriptomic signatures of Alzheimer’s Diseases that are partially reversed by mitochondrial complex I inhibitor. BioRxiv 2024.

