## Supplementary Figure 1 for "Therapeutic assessment of a novel mitochondrial complex I inhibitor in *in vitro* and *in vivo* models of Alzheimer’s disease"

### Slide 1
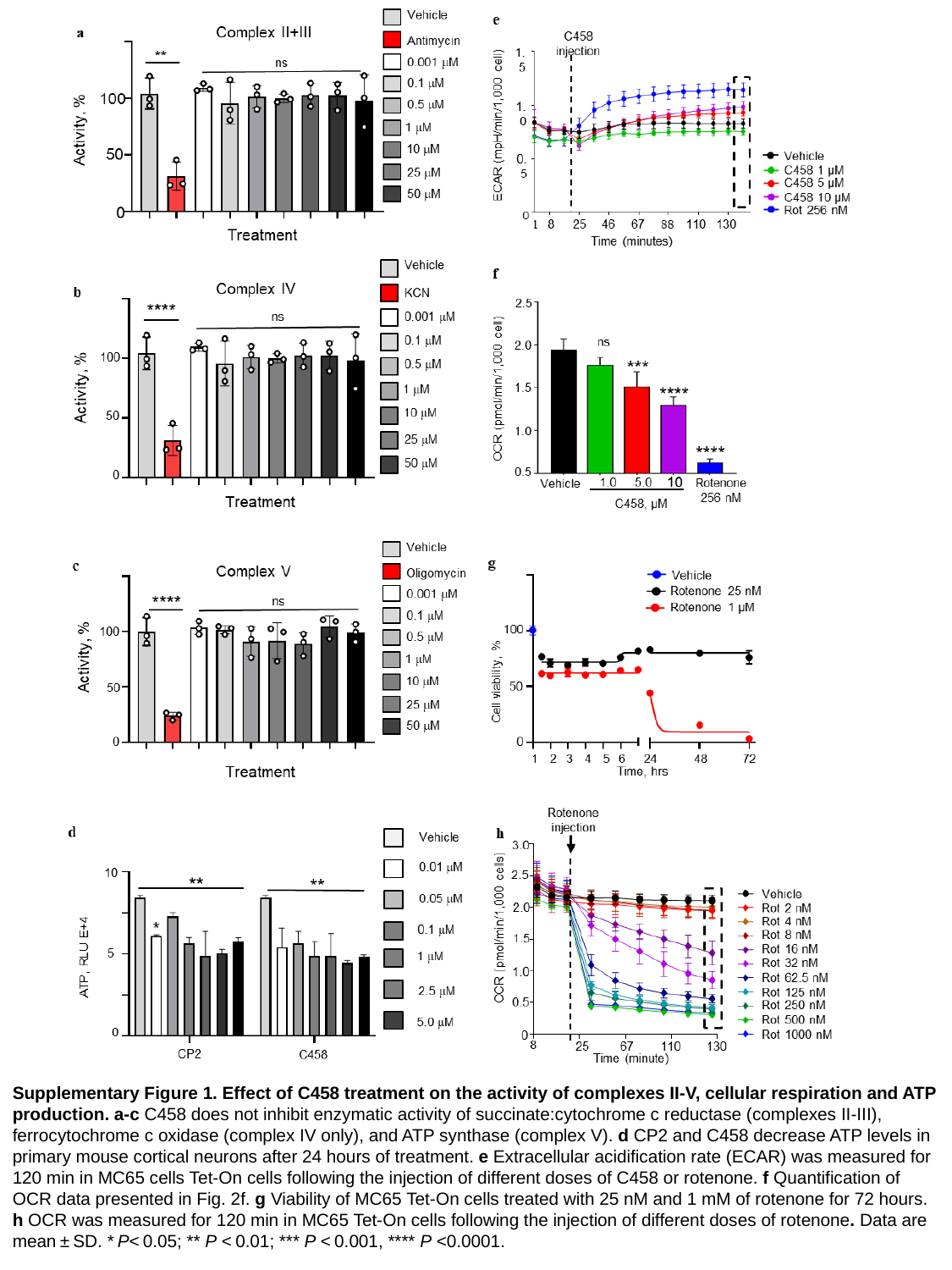

Supplementary Figure 1. Effect of C458 treatment on the activity of complexes II-V, cellular respiration and ATP production. a-c C458 does not inhibit enzymatic activity of succinate:cytochrome c reductase (complexes II-III), ferrocytochrome c oxidase (complex IV only), and ATP synthase (complex V). d CP2 and C458 decrease ATP levels in primary mouse cortical neurons after 24 hours of treatment. e Extracellular acidification rate (ECAR) was measured for 120 min in MC65 cells Tet-On cells following the injection of different doses of C458 or rotenone. f Quantification of OCR data presented in Fig. 2f. g Viability of MC65 Tet-On cells treated with 25 nM and 1 mM of rotenone for 72 hours. h OCR was measured for 120 min in MC65 Tet-On cells following the injection of different doses of rotenone. Data are mean ± SD. * P< 0.05; ** P < 0.01; *** P < 0.001, **** P <0.0001.
