## Supplementary Table 1 for "Therapeutic assessment of a novel mitochondrial complex I inhibitor in *in vitro* and *in vivo* models of Alzheimer’s disease"

### Slide 1
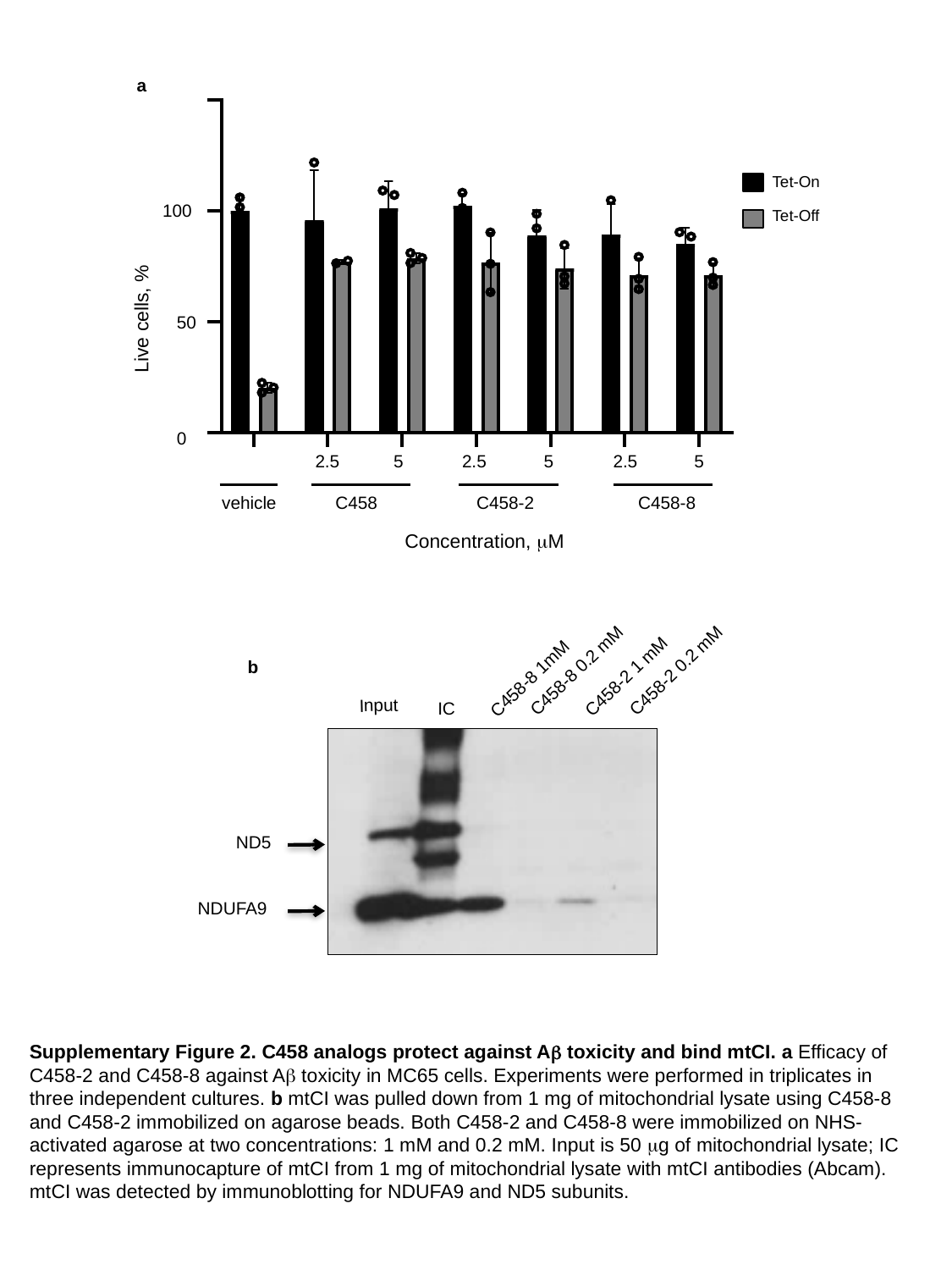

a
Tet-On
100
Tet-Off
Live cells, %
50
0
2.5
5
2.5
5
2.5
5
vehicle
C458
C458-2
C458-8
Concentration, μM
b
 C458-8 0.2 mM
 C458-2 0.2 mM
 C458-2 1 mM
 C458-8 1mM
Input
IC
ND5
NDUFA9
Supplementary Figure 2. C458 analogs protect against Aβ toxicity and bind mtCI. a Efficacy of C458-2 and C458-8 against Aβ toxicity in MC65 cells. Experiments were performed in triplicates in three independent cultures. b mtCI was pulled down from 1 mg of mitochondrial lysate using C458-8 and C458-2 immobilized on agarose beads. Both C458-2 and C458-8 were immobilized on NHS-activated agarose at two concentrations: 1 mM and 0.2 mM. Input is 50 μg of mitochondrial lysate; IC represents immunocapture of mtCI from 1 mg of mitochondrial lysate with mtCI antibodies (Abcam). mtCI was detected by immunoblotting for NDUFA9 and ND5 subunits.
