## Supplementary Figure 2 for "Therapeutic assessment of a novel mitochondrial complex I inhibitor in *in vitro* and *in vivo* models of Alzheimer’s disease"

### Slide 1
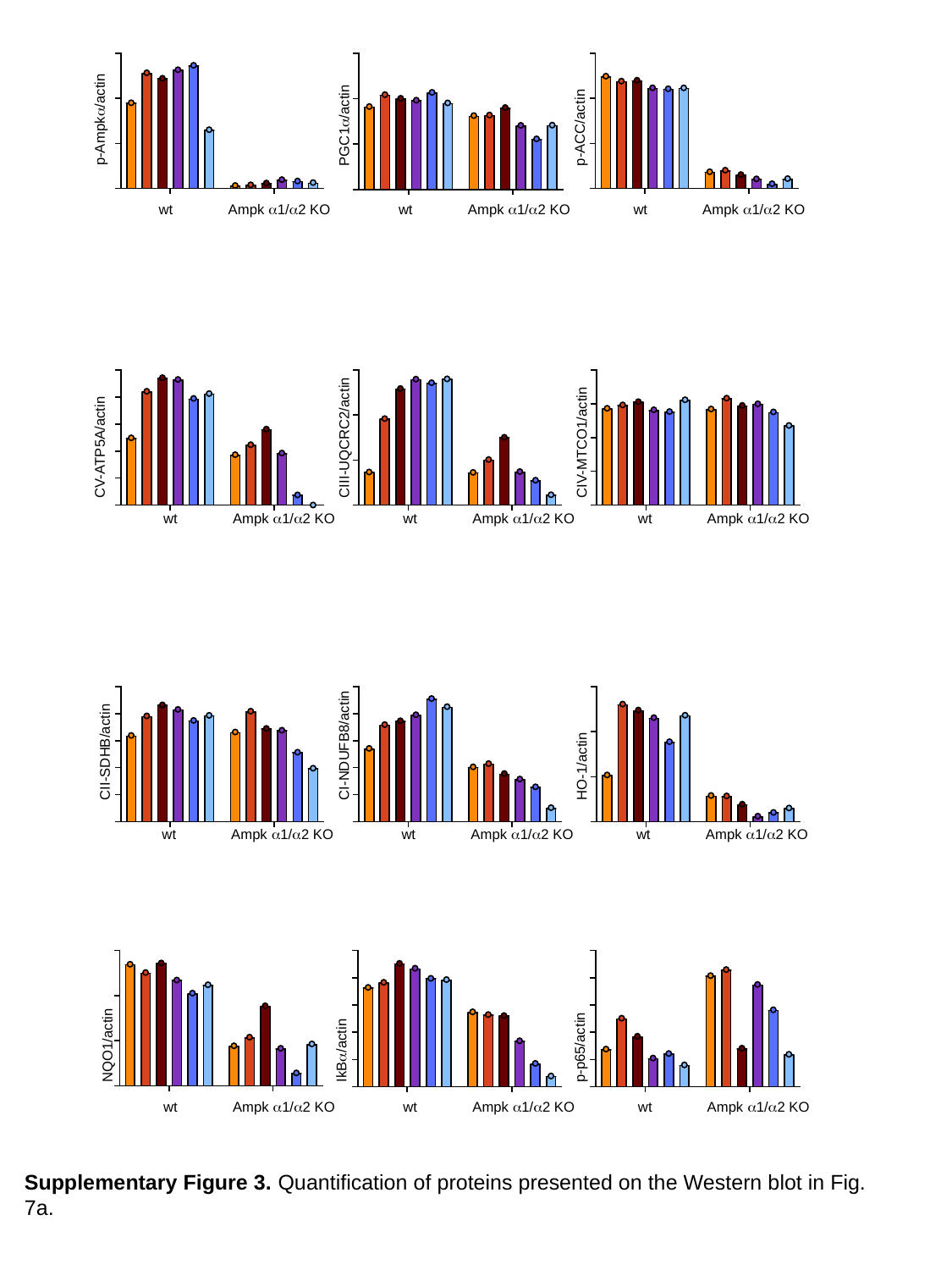

p-Ampkα/actin
PGC1α/actin
p-ACC/actin
wt
Ampk α1/α2 KO
wt
Ampk α1/α2 KO
wt
Ampk α1/α2 KO
CV-ATP5A/actin
CIII-UQCRC2/actin
CIV-MTCO1/actin
wt
Ampk α1/α2 KO
wt
Ampk α1/α2 KO
wt
Ampk α1/α2 KO
CI-NDUFB8/actin
CII-SDHB/actin
HO-1/actin
wt
Ampk α1/α2 KO
wt
Ampk α1/α2 KO
wt
Ampk α1/α2 KO
NQO1/actin
p-p65/actin
IkBα/actin
wt
Ampk α1/α2 KO
wt
Ampk α1/α2 KO
wt
Ampk α1/α2 KO
Supplementary Figure 3. Quantification of proteins presented on the Western blot in Fig. 7a.
