## Supplementary Figure 3 for "Therapeutic assessment of a novel mitochondrial complex I inhibitor in *in vitro* and *in vivo* models of Alzheimer’s disease"

### Slide 1
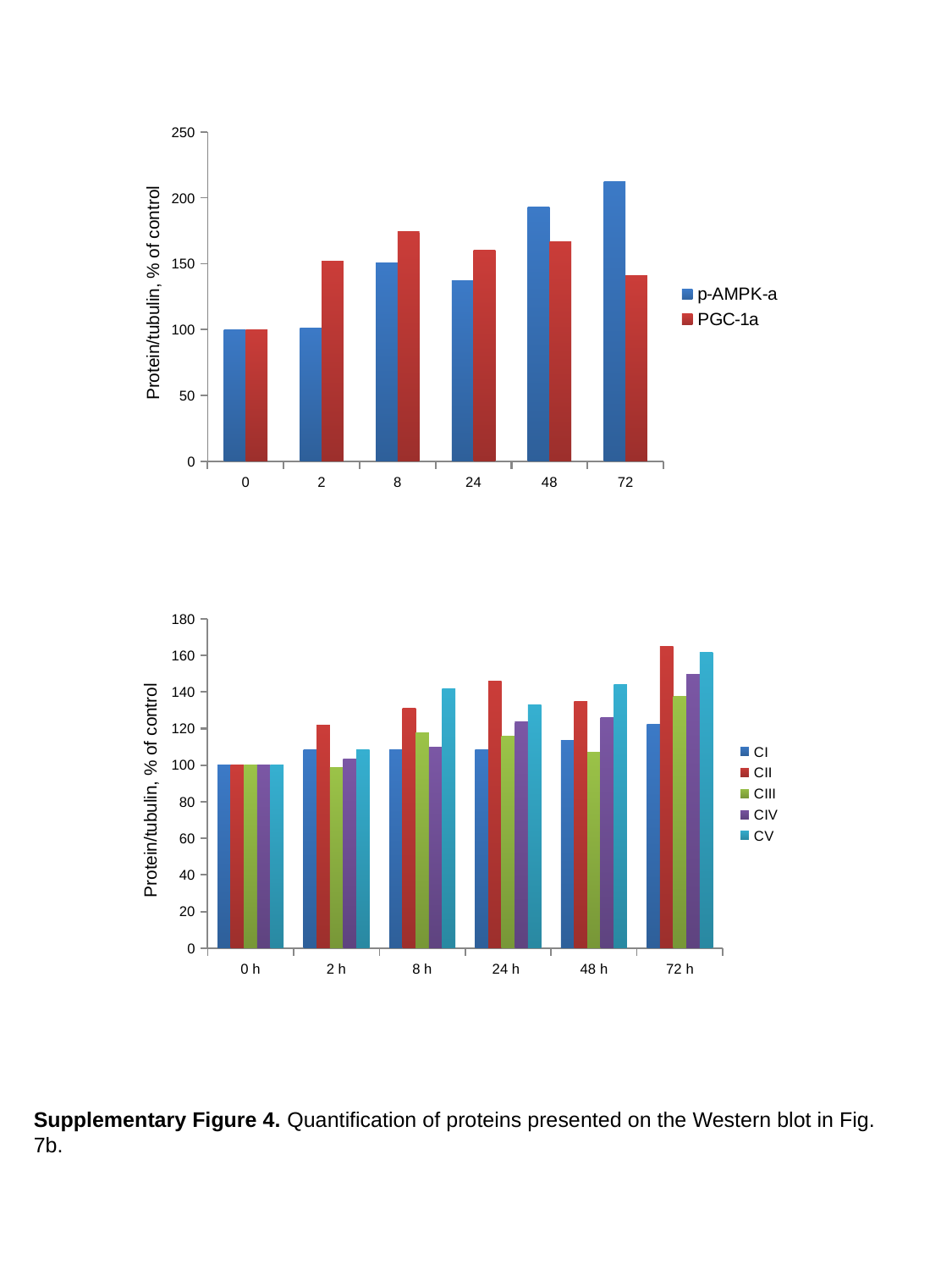

#### Chart
| Category | p-AMPK-a | PGC-1a |
|---|---|---|
| 0 | 100.00165968671413 | 100.01759643821788 |
| 2 | 101.03057266469382 | 151.9397506139899 |
| 8 | 150.34189839020237 | 174.26762642919468 |
| 24 | 136.93846098060683 | 160.33375155674392 |
| 48 | 193.35169962611428 | 166.9217599961859 |
| 72 | 212.24835996134254 | 141.1489273003691 |Protein/tubulin, % of control
#### Chart
| Category | CI | CII | CIII | CIV | CV |
|---|---|---|---|---|---|
| 0 h | 100.0 | 100.0 | 100.0 | 100.0 | 100.0 |
| 2 h | 108.42105263157895 | 121.8801874303965 | 98.70224793362725 | 103.26086956521738 | 108.65289731922591 |
| 8 h | 108.42105263157895 | 131.2601122060894 | 117.71832015364639 | 109.78260869565217 | 141.96453045337427 |
| 24 h | 108.42105263157895 | 145.9109915740371 | 115.83039803432162 | 123.50543478260869 | 132.8408530234088 |
| 48 h | 113.68421052631578 | 135.00661889853123 | 106.88453972178809 | 125.95108695652173 | 143.95318239131626 |
| 72 h | 122.10526315789474 | 165.01019100250048 | 137.3611078712667 | 149.72826086956522 | 161.42755331396305 |Protein/tubulin, % of control
Supplementary Figure 4. Quantification of proteins presented on the Western blot in Fig. 7b.
