## Supplementary Figure 4 for "Therapeutic assessment of a novel mitochondrial complex I inhibitor in *in vitro* and *in vivo* models of Alzheimer’s disease"

### Slide 1
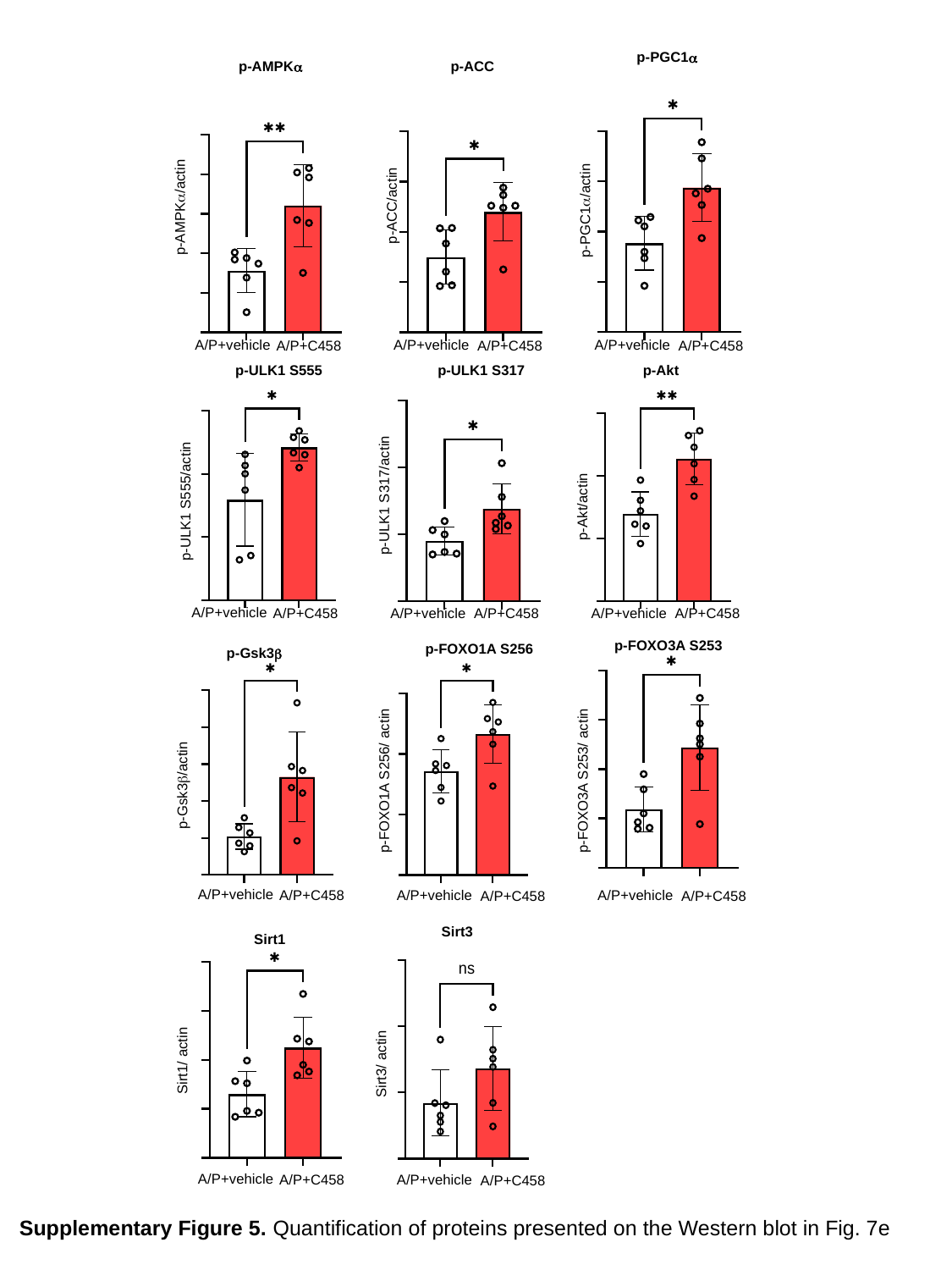

p-PGC1α
p-AMPKα
p-ACC
p-ACC/actin
p-AMPKα/actin
p-PGC1α/actin
A/P+vehicle
A/P+C458
A/P+vehicle
A/P+C458
A/P+vehicle
A/P+C458
p-ULK1 S555
p-Akt
p-ULK1 S317
p-ULK1 S317/actin
p-Akt/actin
p-ULK1 S555/actin
A/P+vehicle
A/P+C458
A/P+vehicle
A/P+C458
A/P+vehicle
A/P+C458
p-FOXO3A S253
p-FOXO1A S256
p-Gsk3β
p-FOXO3A S253/ actin
p-Gsk3β/actin
p-FOXO1A S256/ actin
A/P+vehicle
A/P+C458
A/P+vehicle
A/P+C458
A/P+vehicle
A/P+C458
Sirt3
Sirt1
Sirt1/ actin
Sirt3/ actin
A/P+vehicle
A/P+C458
A/P+vehicle
A/P+C458
Supplementary Figure 5. Quantification of proteins presented on the Western blot in Fig. 7e
