## Supplementary figures and images for "Therapeutic assessment of a novel mitochondrial complex I inhibitor in *in vitro* and *in vivo* models of Alzheimer’s disease"

### Supplementary Figure 5

#### Slide 1
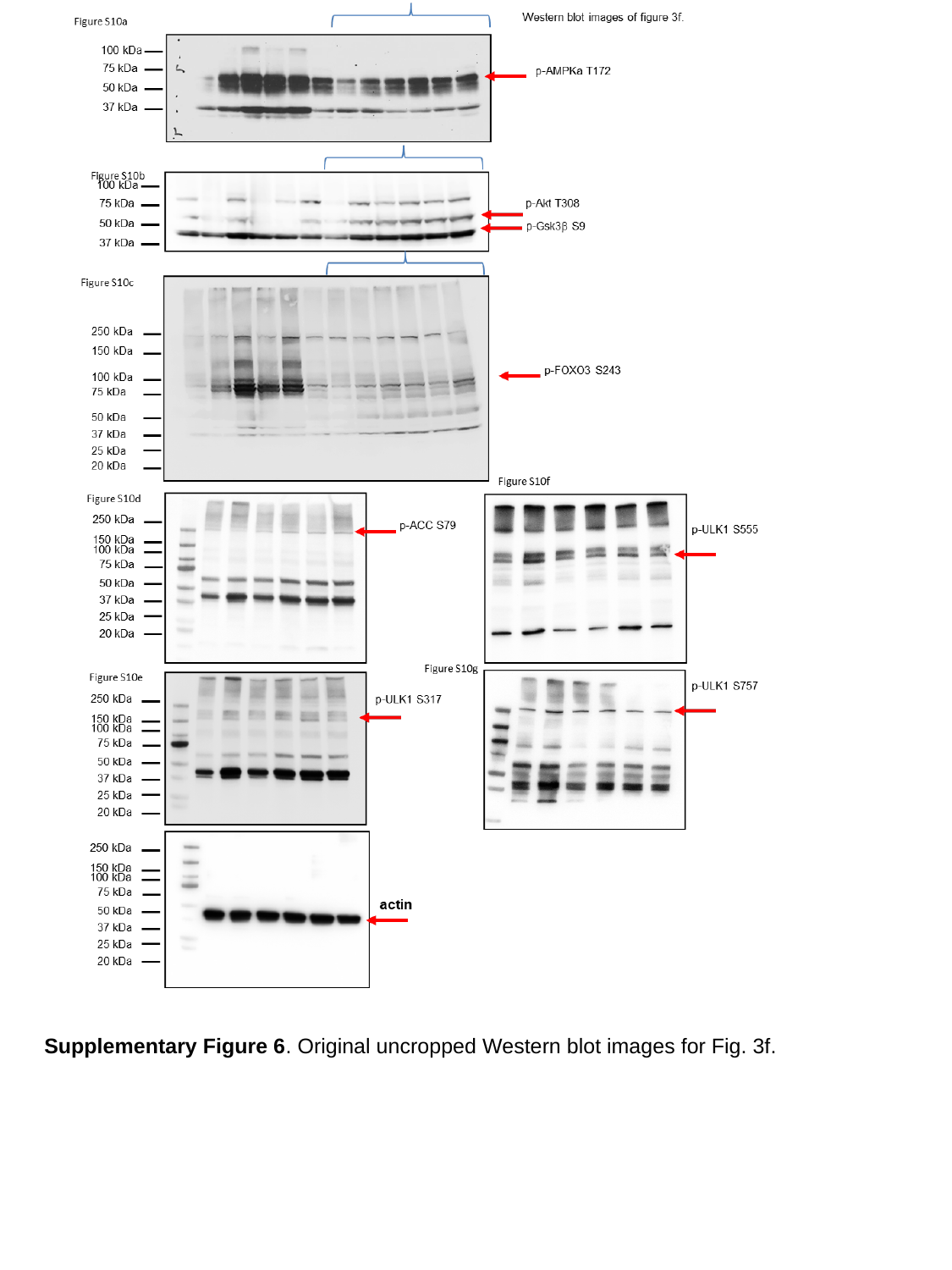

Supplementary Figure 6. Original uncropped Western blot images for Fig. 3f.
