## Supplementary Figure 6 for "Therapeutic assessment of a novel mitochondrial complex I inhibitor in *in vitro* and *in vivo* models of Alzheimer’s disease"

### Slide 1
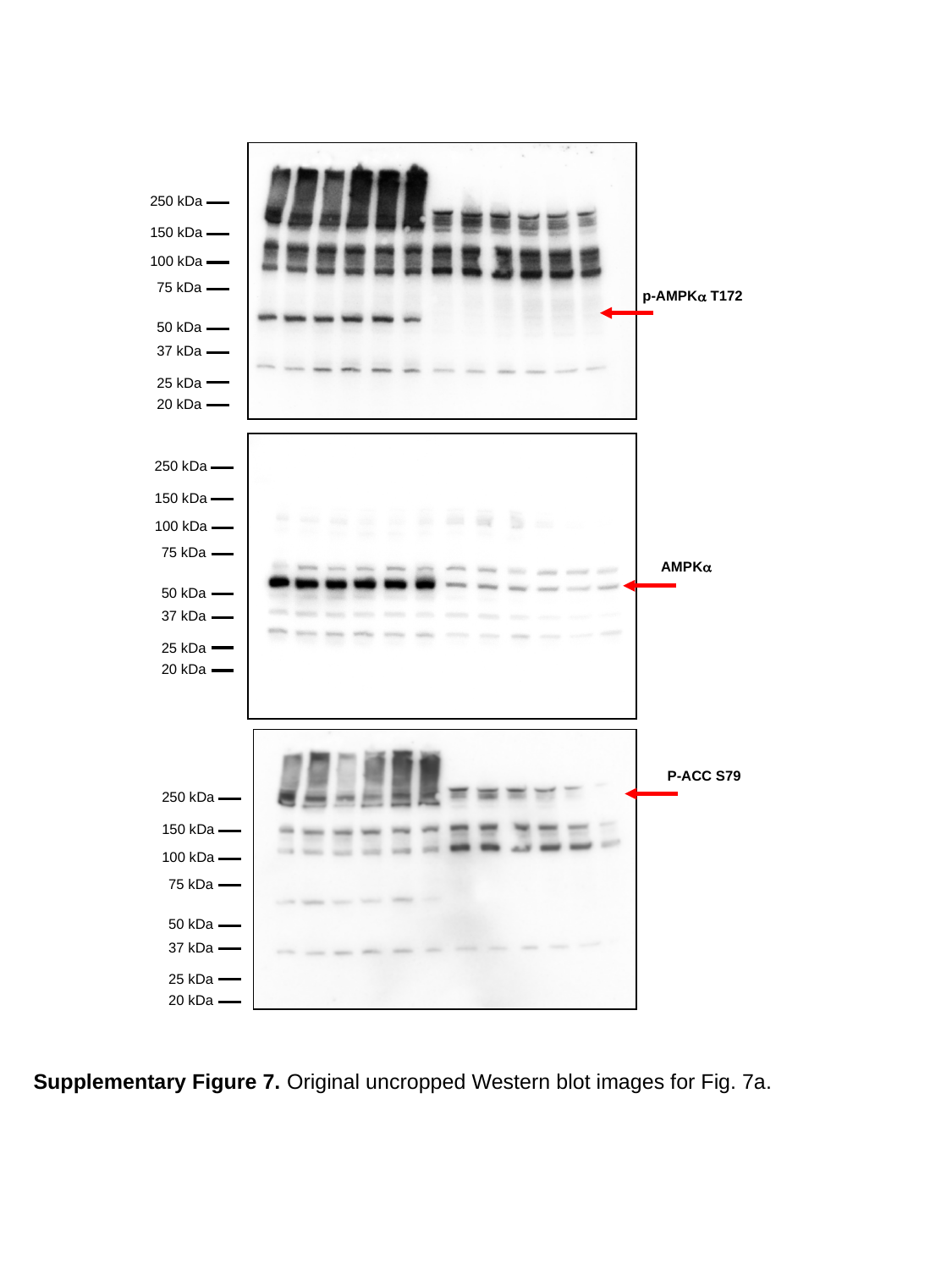

250 kDa
150 kDa
100 kDa
75 kDa
50 kDa
37 kDa
25 kDa
20 kDa
p-AMPKα T172
250 kDa
150 kDa
100 kDa
75 kDa
50 kDa
37 kDa
25 kDa
20 kDa
AMPKα
P-ACC S79
250 kDa
150 kDa
100 kDa
75 kDa
50 kDa
37 kDa
25 kDa
20 kDa
Supplementary Figure 7. Original uncropped Western blot images for Fig. 7a.

### Slide 2
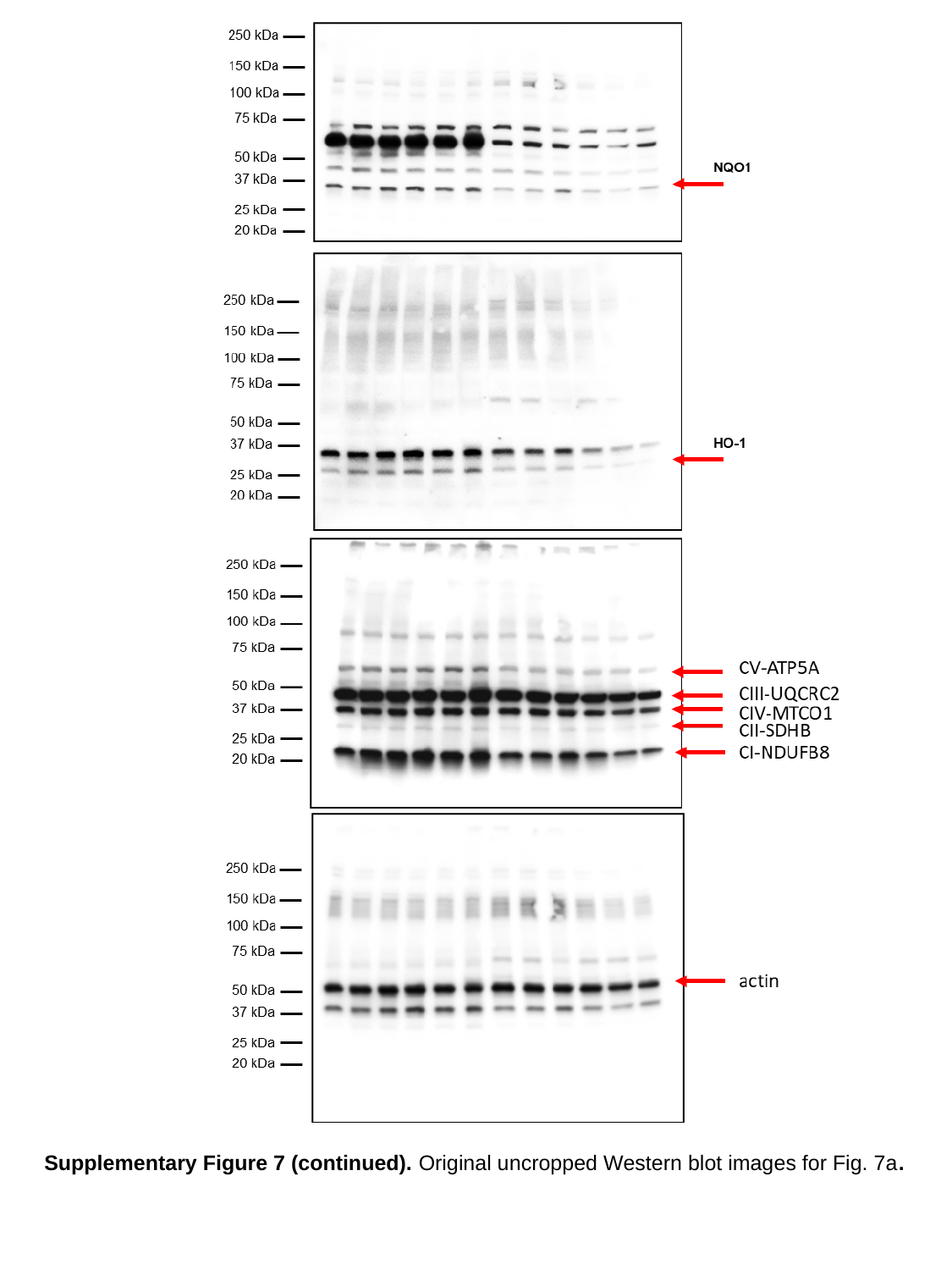

Supplementary Figure 7 (continued). Original uncropped Western blot images for Fig. 7a.

### Slide 3
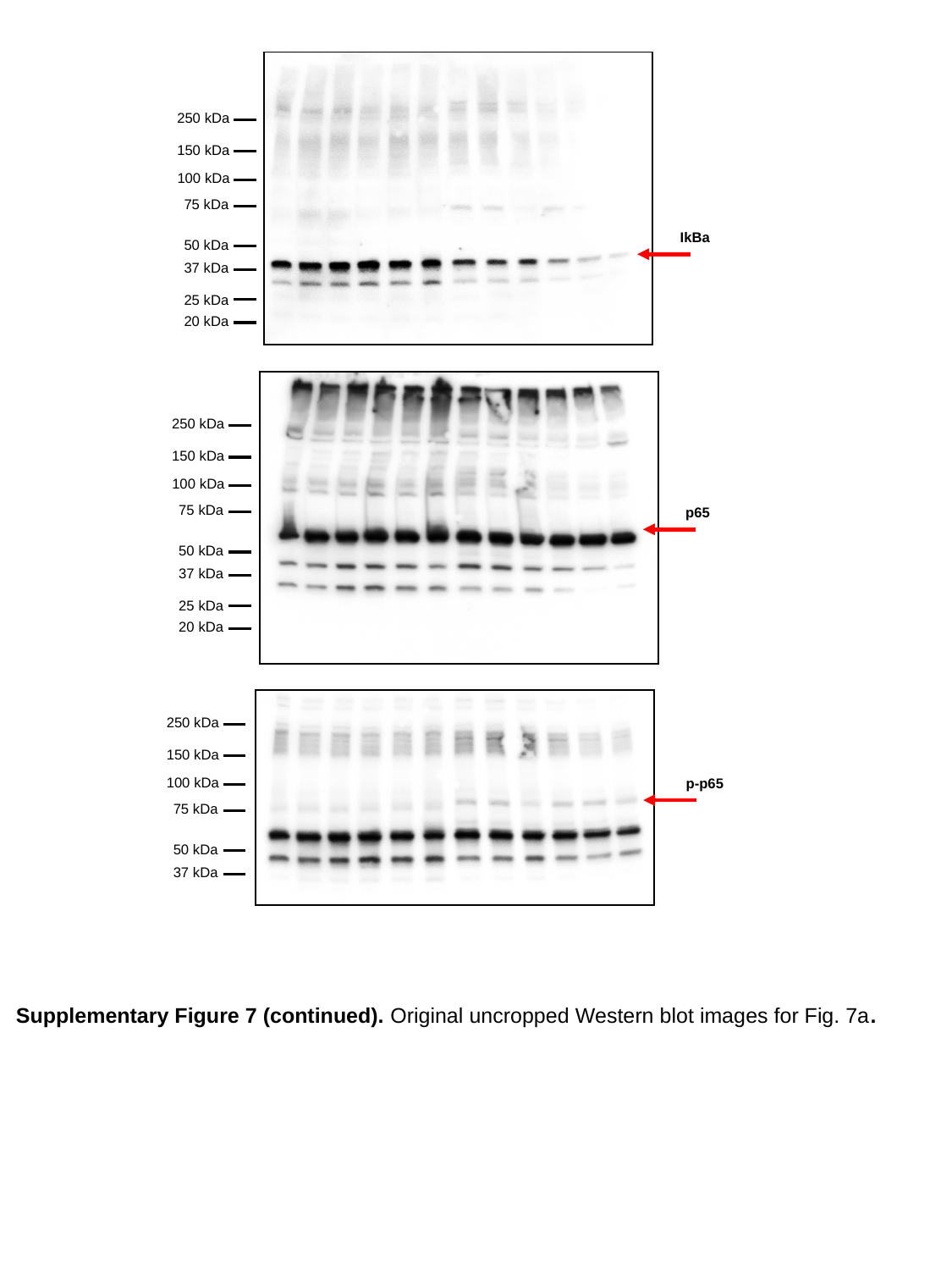

250 kDa
150 kDa
100 kDa
75 kDa
50 kDa
37 kDa
25 kDa
20 kDa
IkBa
250 kDa
150 kDa
100 kDa
75 kDa
50 kDa
37 kDa
25 kDa
20 kDa
p65
250 kDa
150 kDa
100 kDa
75 kDa
50 kDa
37 kDa
p-p65
Supplementary Figure 7 (continued). Original uncropped Western blot images for Fig. 7a.
