## Supplementary Figure 7 for "Therapeutic assessment of a novel mitochondrial complex I inhibitor in *in vitro* and *in vivo* models of Alzheimer’s disease"

### Slide 1
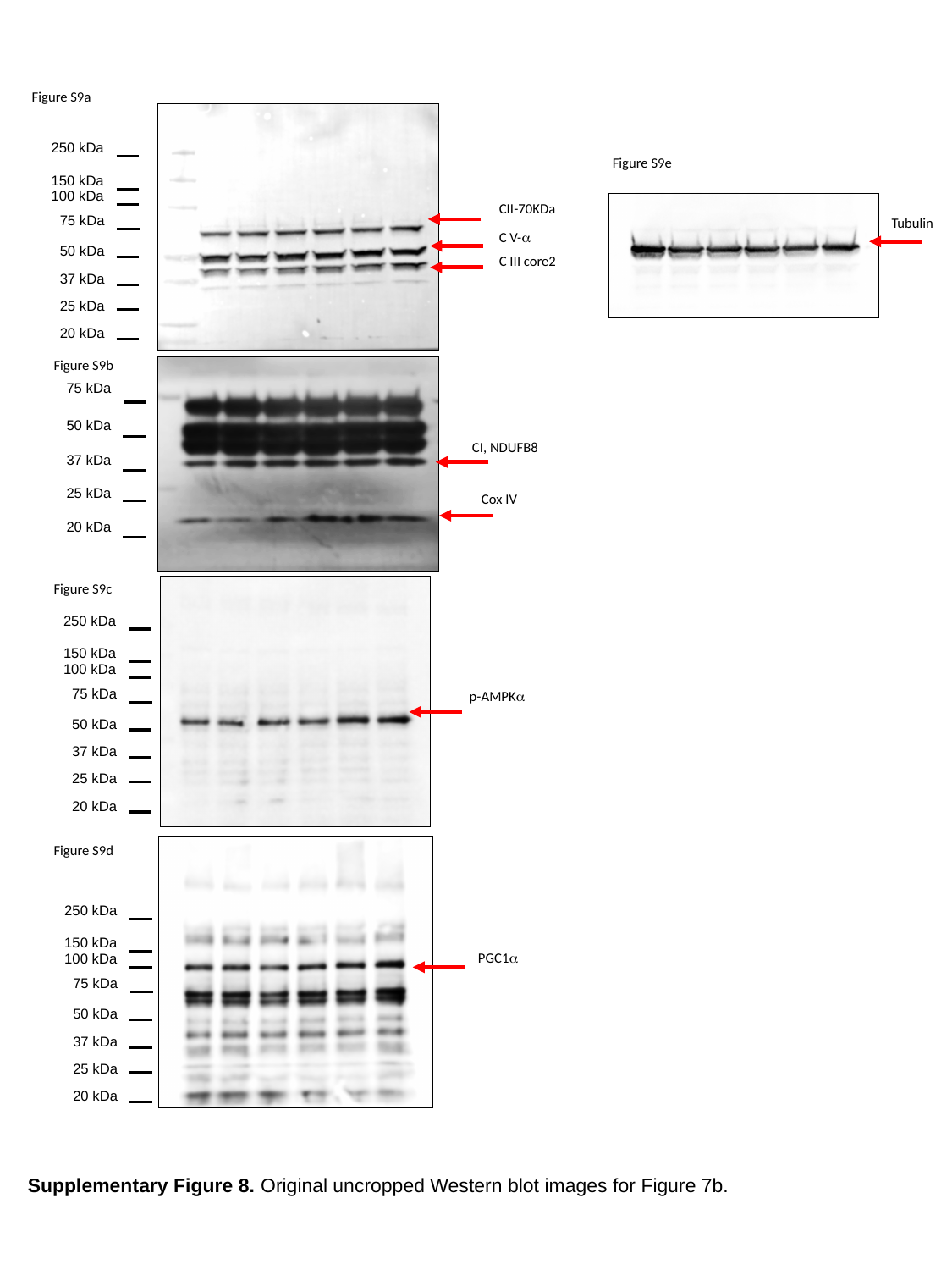

Figure S9a
250 kDa
150 kDa
100 kDa
75 kDa
50 kDa
37 kDa
25 kDa
20 kDa
Figure S9e
CII-70KDa
Tubulin
C V-α
C III core2
Figure S9b
75 kDa
50 kDa
37 kDa
25 kDa
20 kDa
CI, NDUFB8
Cox IV
Figure S9c
250 kDa
150 kDa
100 kDa
75 kDa
50 kDa
37 kDa
25 kDa
20 kDa
p-AMPKα
Figure S9d
250 kDa
150 kDa
100 kDa
75 kDa
50 kDa
37 kDa
25 kDa
20 kDa
PGC1α
Supplementary Figure 8. Original uncropped Western blot images for Figure 7b.
