## Supplementary Figure 8 for "Therapeutic assessment of a novel mitochondrial complex I inhibitor in *in vitro* and *in vivo* models of Alzheimer’s disease"

### Slide 1
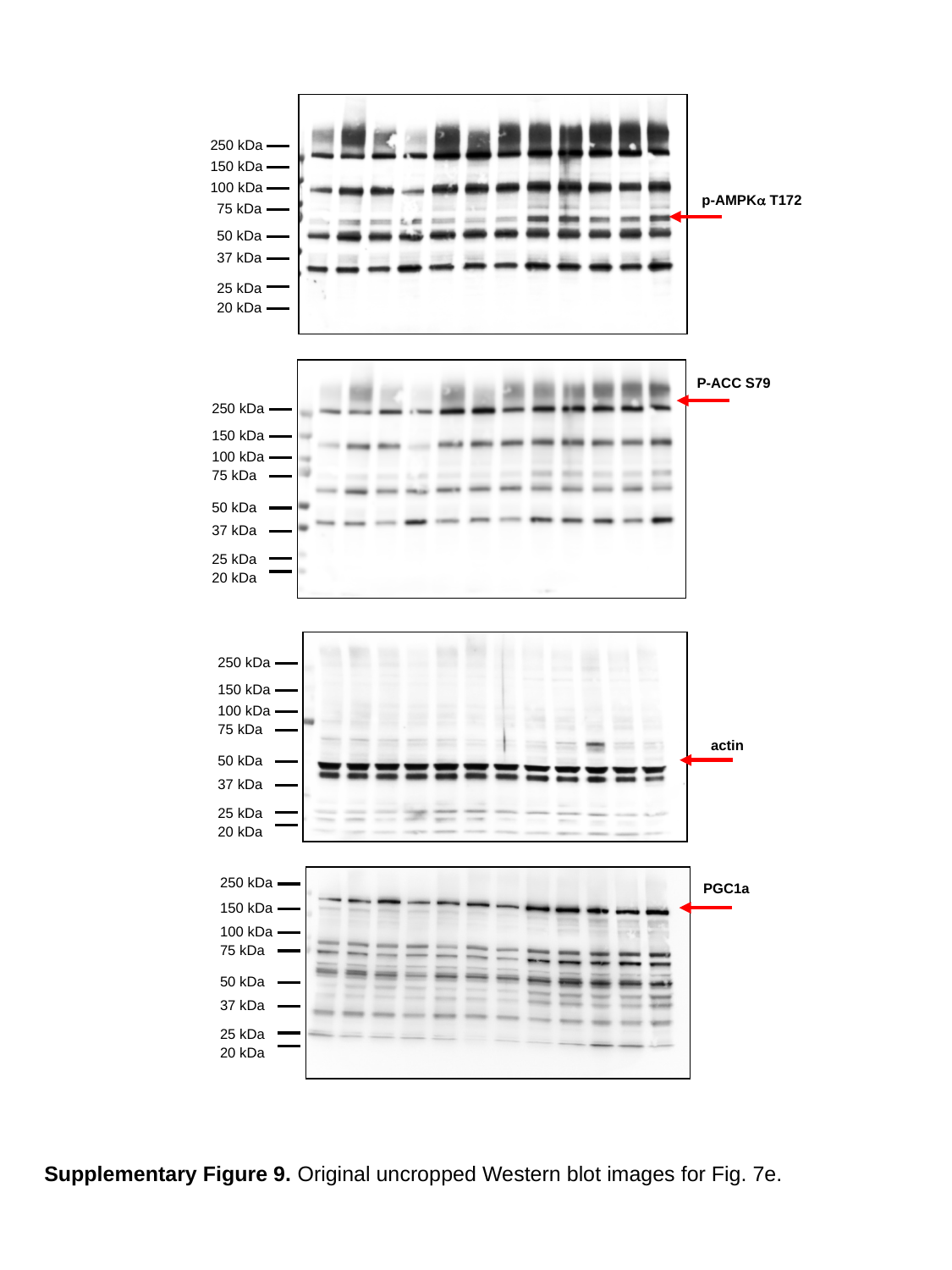

250 kDa
150 kDa
100 kDa
75 kDa
50 kDa
37 kDa
25 kDa
20 kDa
p-AMPKα T172
P-ACC S79
250 kDa
150 kDa
100 kDa
75 kDa
50 kDa
37 kDa
25 kDa
20 kDa
250 kDa
150 kDa
100 kDa
75 kDa
50 kDa
37 kDa
25 kDa
20 kDa
actin
250 kDa
150 kDa
100 kDa
75 kDa
50 kDa
37 kDa
25 kDa
20 kDa
PGC1a
Supplementary Figure 9. Original uncropped Western blot images for Fig. 7e.

### Slide 2
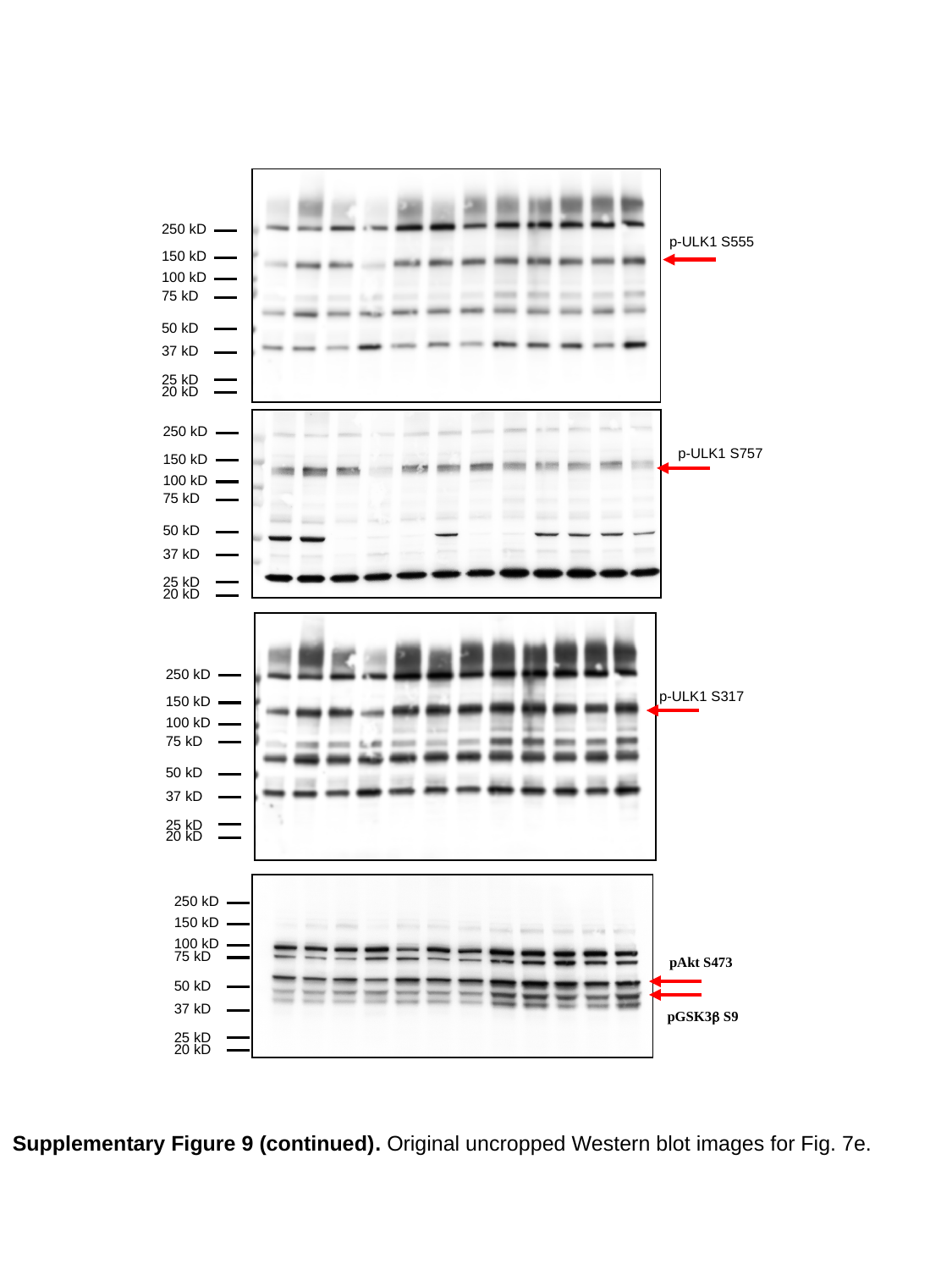

250 kD
150 kD
100 kD
75 kD
50 kD
37 kD
25 kD
20 kD
p-ULK1 S555
250 kD
150 kD
100 kD
75 kD
50 kD
37 kD
25 kD
20 kD
p-ULK1 S757
250 kD
150 kD
100 kD
75 kD
50 kD
37 kD
25 kD
20 kD
p-ULK1 S317
250 kD
150 kD
100 kD
75 kD
50 kD
37 kD
25 kD
20 kD
pAkt S473
pGSK3β S9
Supplementary Figure 9 (continued). Original uncropped Western blot images for Fig. 7e.

### Slide 3
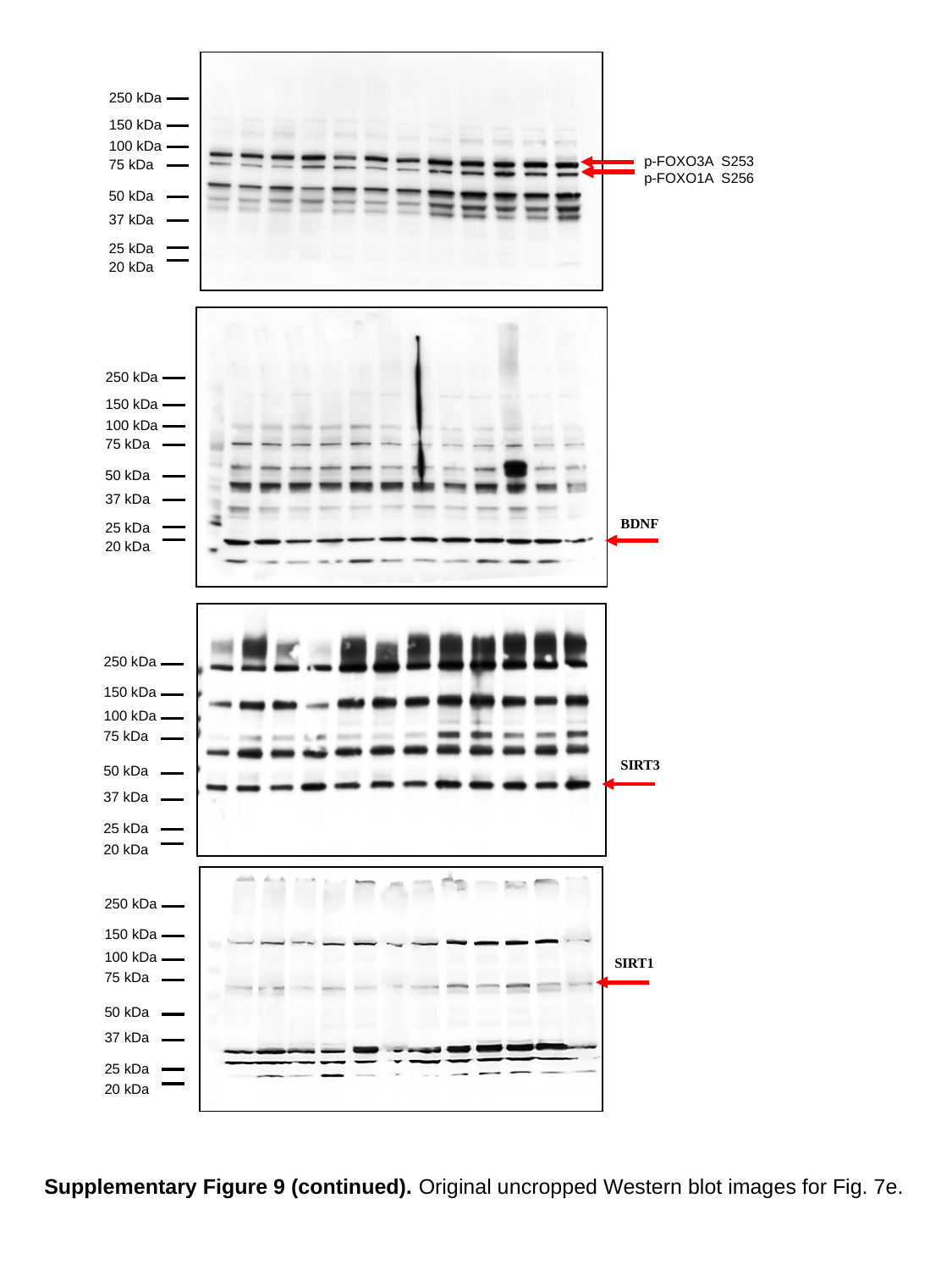

250 kDa
150 kDa
100 kDa
75 kDa
50 kDa
37 kDa
25 kDa
20 kDa
p-FOXO3A S253
p-FOXO1A S256
250 kDa
150 kDa
100 kDa
75 kDa
50 kDa
37 kDa
25 kDa
20 kDa
BDNF
250 kDa
150 kDa
100 kDa
75 kDa
50 kDa
37 kDa
25 kDa
20 kDa
SIRT3
250 kDa
150 kDa
100 kDa
75 kDa
50 kDa
37 kDa
25 kDa
20 kDa
SIRT1
Supplementary Figure 9 (continued). Original uncropped Western blot images for Fig. 7e.
