## Supplementary Figure 9 for "Therapeutic assessment of a novel mitochondrial complex I inhibitor in *in vitro* and *in vivo* models of Alzheimer’s disease"

### Slide 1
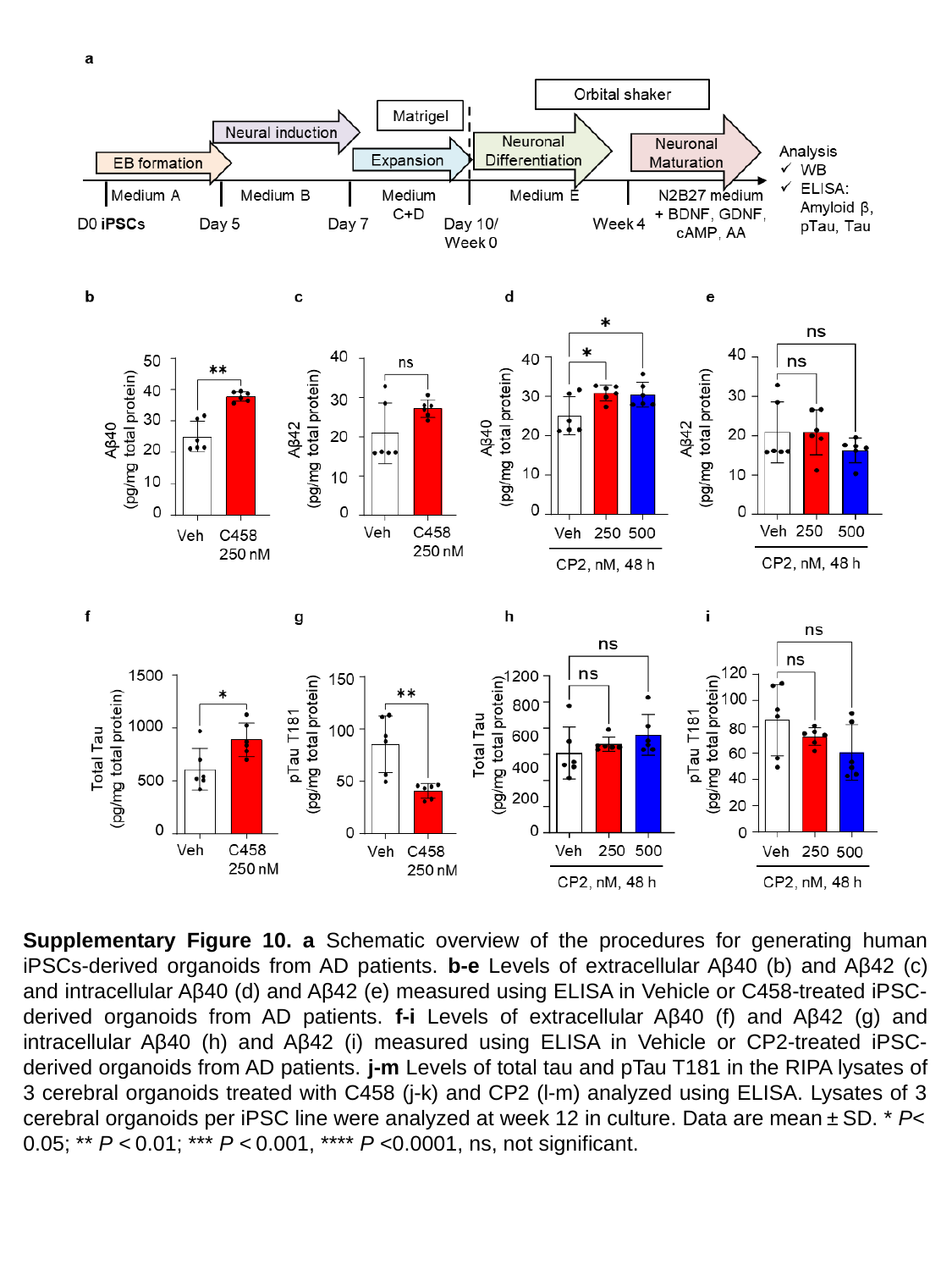

Supplementary Figure 10. a Schematic overview of the procedures for generating human iPSCs-derived organoids from AD patients. b-e Levels of extracellular Aβ40 (b) and Aβ42 (c) and intracellular Aβ40 (d) and Aβ42 (e) measured using ELISA in Vehicle or C458-treated iPSC-derived organoids from AD patients. f-i Levels of extracellular Aβ40 (f) and Aβ42 (g) and intracellular Aβ40 (h) and Aβ42 (i) measured using ELISA in Vehicle or CP2-treated iPSC-derived organoids from AD patients. j-m Levels of total tau and pTau T181 in the RIPA lysates of 3 cerebral organoids treated with C458 (j-k) and CP2 (l-m) analyzed using ELISA. Lysates of 3 cerebral organoids per iPSC line were analyzed at week 12 in culture. Data are mean ± SD. * P< 0.05; ** P < 0.01; *** P < 0.001, **** P <0.0001, ns, not significant.
