## Supplementary Figure 10 for "Therapeutic assessment of a novel mitochondrial complex I inhibitor in *in vitro* and *in vivo* models of Alzheimer’s disease"

### Slide 1
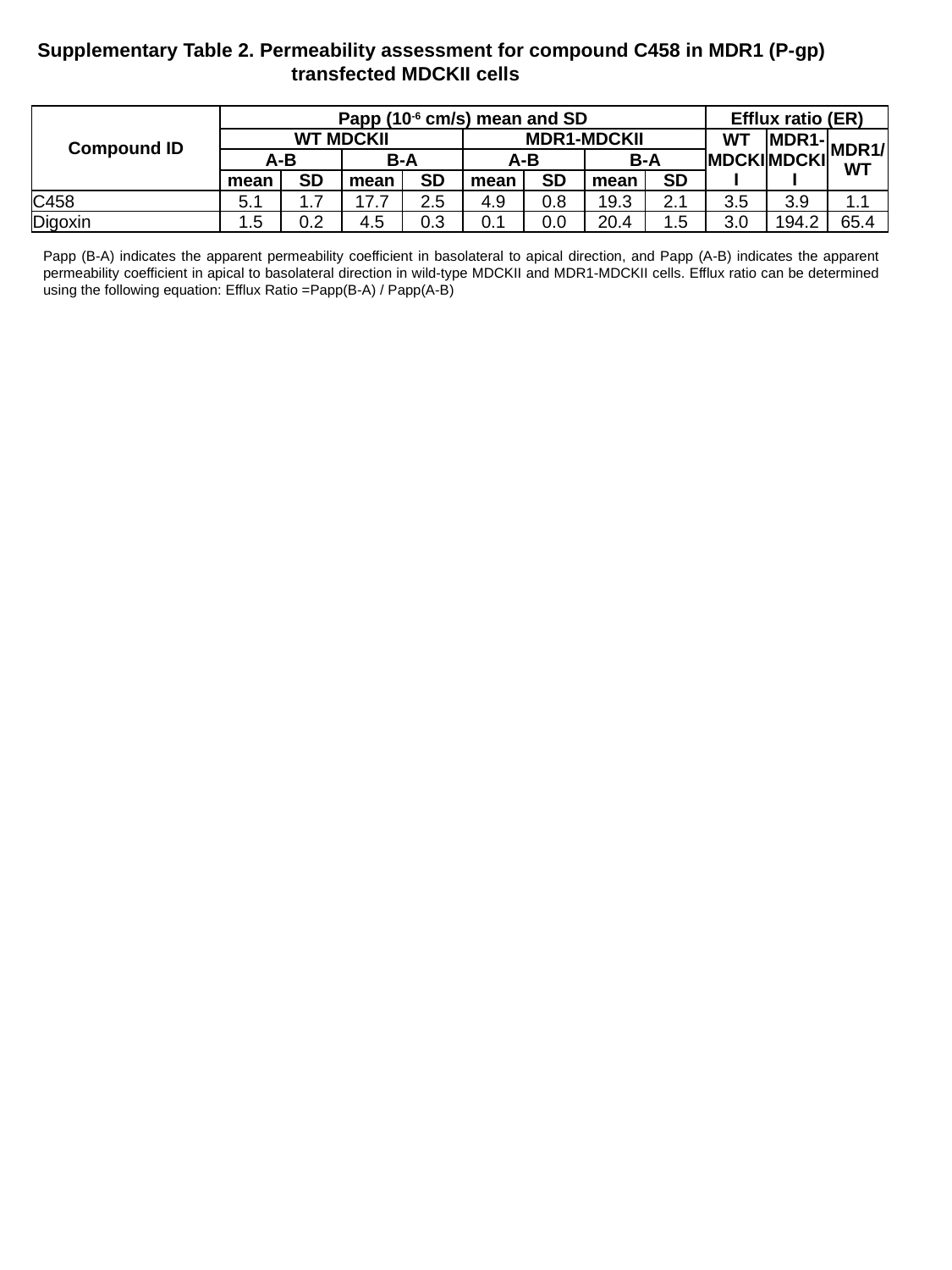

Supplementary Table 2. Permeability assessment for compound C458 in MDR1 (P-gp) 		transfected MDCKII cells
| Compound ID | Papp (10^-6^ cm/s) mean and SD | | | | | | | | Efflux ratio (ER) | | |
| --- | --- | --- | --- | --- | --- | --- | --- | --- | --- | --- | --- |
| | WT MDCKII | | | | MDR1-MDCKII | | | | WT MDCKII | MDR1-MDCKII | MDR1/ WT |
| | A-B | | B-A | | A-B | | B-A | | | | |
| | mean | SD | mean | SD | mean | SD | mean | SD | | | |
| C458 | 5.1 | 1.7 | 17.7 | 2.5 | 4.9 | 0.8 | 19.3 | 2.1 | 3.5 | 3.9 | 1.1 |
| Digoxin | 1.5 | 0.2 | 4.5 | 0.3 | 0.1 | 0.0 | 20.4 | 1.5 | 3.0 | 194.2 | 65.4 |
Papp (B-A) indicates the apparent permeability coefficient in basolateral to apical direction, and Papp (A-B) indicates the apparent permeability coefficient in apical to basolateral direction in wild-type MDCKII and MDR1-MDCKII cells. Efflux ratio can be determined using the following equation: Efflux Ratio =Papp(B-A) / Papp(A-B)
