## Supplementary Table 2 for "Therapeutic assessment of a novel mitochondrial complex I inhibitor in *in vitro* and *in vivo* models of Alzheimer’s disease"

### Slide 1
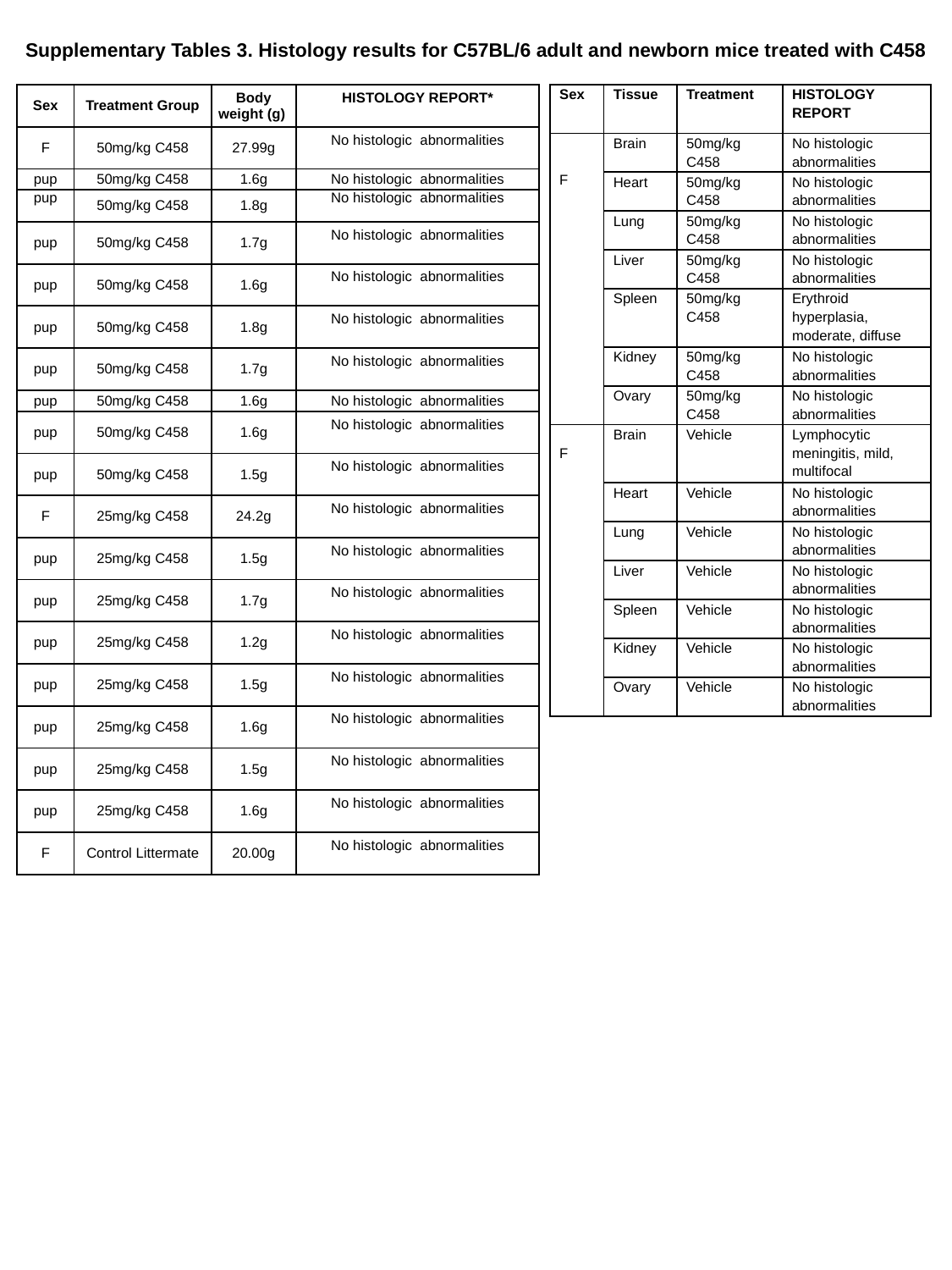

Supplementary Tables 3. Histology results for C57BL/6 adult and newborn mice treated with C458
| Sex | Treatment Group | Body weight (g) | HISTOLOGY REPORT\* |
| --- | --- | --- | --- |
| F | 50mg/kg C458 | 27.99g | No histologic abnormalities |
| pup | 50mg/kg C458 | 1.6g | No histologic abnormalities |
| pup | 50mg/kg C458 | 1.8g | No histologic abnormalities |
| pup | 50mg/kg C458 | 1.7g | No histologic abnormalities |
| pup | 50mg/kg C458 | 1.6g | No histologic abnormalities |
| pup | 50mg/kg C458 | 1.8g | No histologic abnormalities |
| pup | 50mg/kg C458 | 1.7g | No histologic abnormalities |
| pup | 50mg/kg C458 | 1.6g | No histologic abnormalities |
| pup | 50mg/kg C458 | 1.6g | No histologic abnormalities |
| pup | 50mg/kg C458 | 1.5g | No histologic abnormalities |
| F | 25mg/kg C458 | 24.2g | No histologic abnormalities |
| pup | 25mg/kg C458 | 1.5g | No histologic abnormalities |
| pup | 25mg/kg C458 | 1.7g | No histologic abnormalities |
| pup | 25mg/kg C458 | 1.2g | No histologic abnormalities |
| pup | 25mg/kg C458 | 1.5g | No histologic abnormalities |
| pup | 25mg/kg C458 | 1.6g | No histologic abnormalities |
| pup | 25mg/kg C458 | 1.5g | No histologic abnormalities |
| pup | 25mg/kg C458 | 1.6g | No histologic abnormalities |
| F | Control Littermate | 20.00g | No histologic abnormalities |
| Sex | Tissue | Treatment | HISTOLOGY REPORT |
| --- | --- | --- | --- |
| F | Brain | 50mg/kg C458 | No histologic abnormalities |
| | Heart | 50mg/kg C458 | No histologic abnormalities |
| | Lung | 50mg/kg C458 | No histologic abnormalities |
| | Liver | 50mg/kg C458 | No histologic abnormalities |
| | Spleen | 50mg/kg C458 | Erythroid hyperplasia, moderate, diffuse |
| | Kidney | 50mg/kg C458 | No histologic abnormalities |
| | Ovary | 50mg/kg C458 | No histologic abnormalities |
| F | Brain | Vehicle | Lymphocytic meningitis, mild, multifocal |
| | Heart | Vehicle | No histologic abnormalities |
| | Lung | Vehicle | No histologic abnormalities |
| | Liver | Vehicle | No histologic abnormalities |
| | Spleen | Vehicle | No histologic abnormalities |
| | Kidney | Vehicle | No histologic abnormalities |
| | Ovary | Vehicle | No histologic abnormalities |
