## Supplementary Table 3 for "Therapeutic assessment of a novel mitochondrial complex I inhibitor in *in vitro* and *in vivo* models of Alzheimer’s disease"

### Slide 1
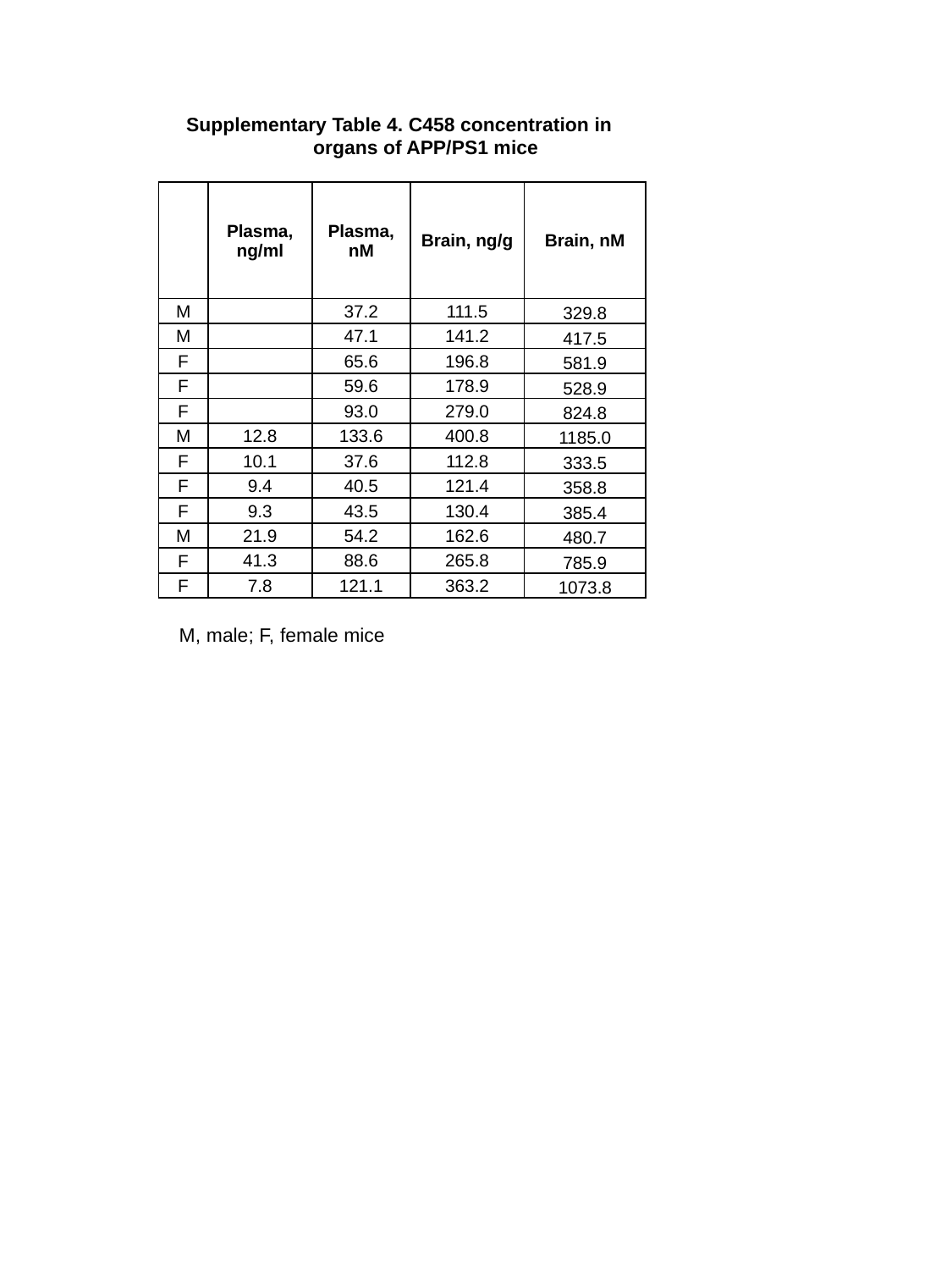

Supplementary Table 4. C458 concentration in 	organs of APP/PS1 mice
| | Plasma, ng/ml | Plasma, nM | Brain, ng/g | Brain, nM |
| --- | --- | --- | --- | --- |
| M | | 37.2 | 111.5 | 329.8 |
| M | | 47.1 | 141.2 | 417.5 |
| F | | 65.6 | 196.8 | 581.9 |
| F | | 59.6 | 178.9 | 528.9 |
| F | | 93.0 | 279.0 | 824.8 |
| M | 12.8 | 133.6 | 400.8 | 1185.0 |
| F | 10.1 | 37.6 | 112.8 | 333.5 |
| F | 9.4 | 40.5 | 121.4 | 358.8 |
| F | 9.3 | 43.5 | 130.4 | 385.4 |
| M | 21.9 | 54.2 | 162.6 | 480.7 |
| F | 41.3 | 88.6 | 265.8 | 785.9 |
| F | 7.8 | 121.1 | 363.2 | 1073.8 |
M, male; F, female mice
