## Supplementary Table 4 for "Therapeutic assessment of a novel mitochondrial complex I inhibitor in *in vitro* and *in vivo* models of Alzheimer’s disease"

### Slide 1
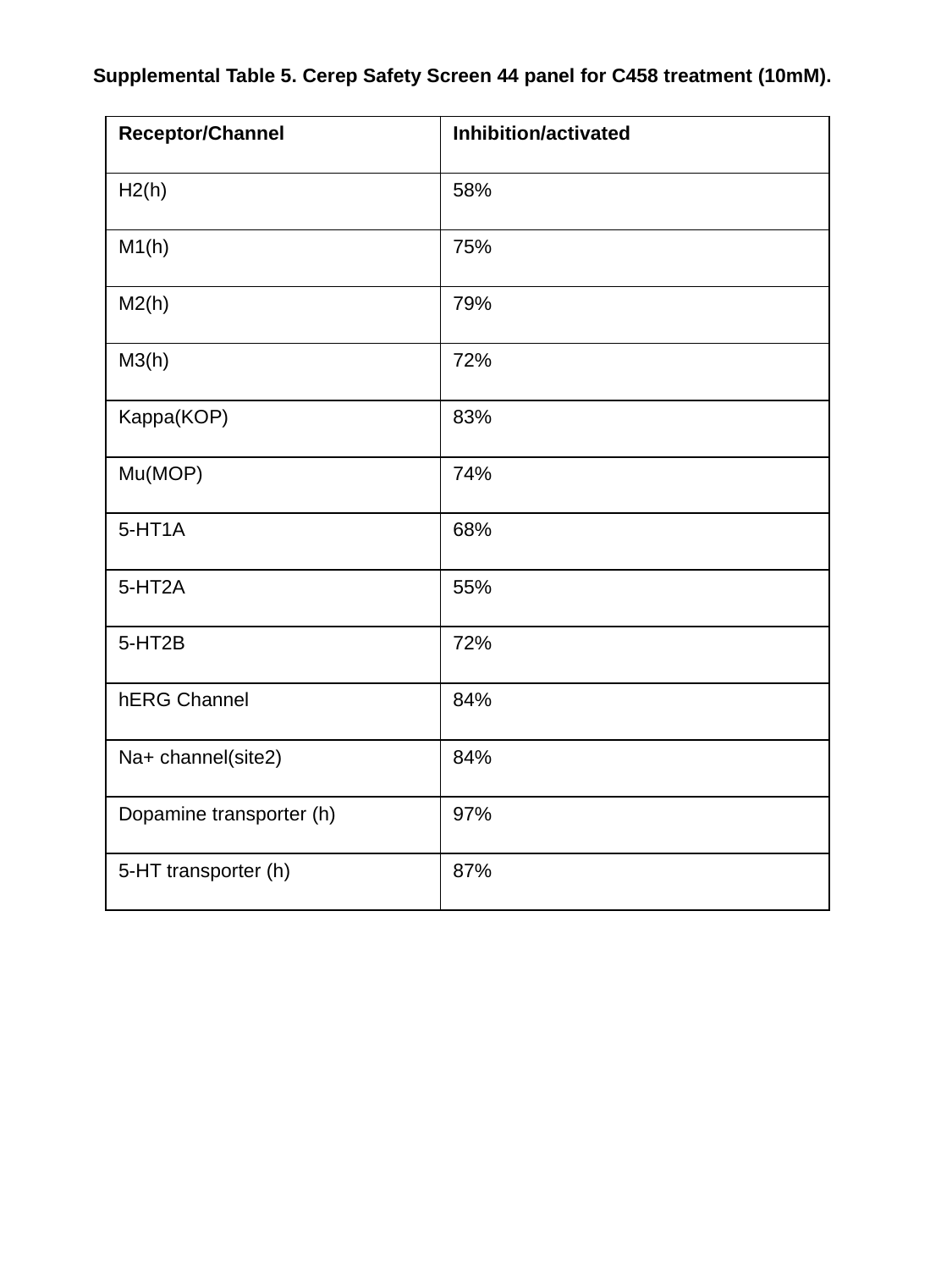

Supplemental Table 5. Cerep Safety Screen 44 panel for C458 treatment (10mM).
| Receptor/Channel | Inhibition/activated |
| --- | --- |
| H2(h) | 58% |
| M1(h) | 75% |
| M2(h) | 79% |
| M3(h) | 72% |
| Kappa(KOP) | 83% |
| Mu(MOP) | 74% |
| 5-HT1A | 68% |
| 5-HT2A | 55% |
| 5-HT2B | 72% |
| hERG Channel | 84% |
| Na+ channel(site2) | 84% |
| Dopamine transporter (h) | 97% |
| 5-HT transporter (h) | 87% |
