## Supplementary Table 5 for "Therapeutic assessment of a novel mitochondrial complex I inhibitor in *in vitro* and *in vivo* models of Alzheimer’s disease"

### Slide 1
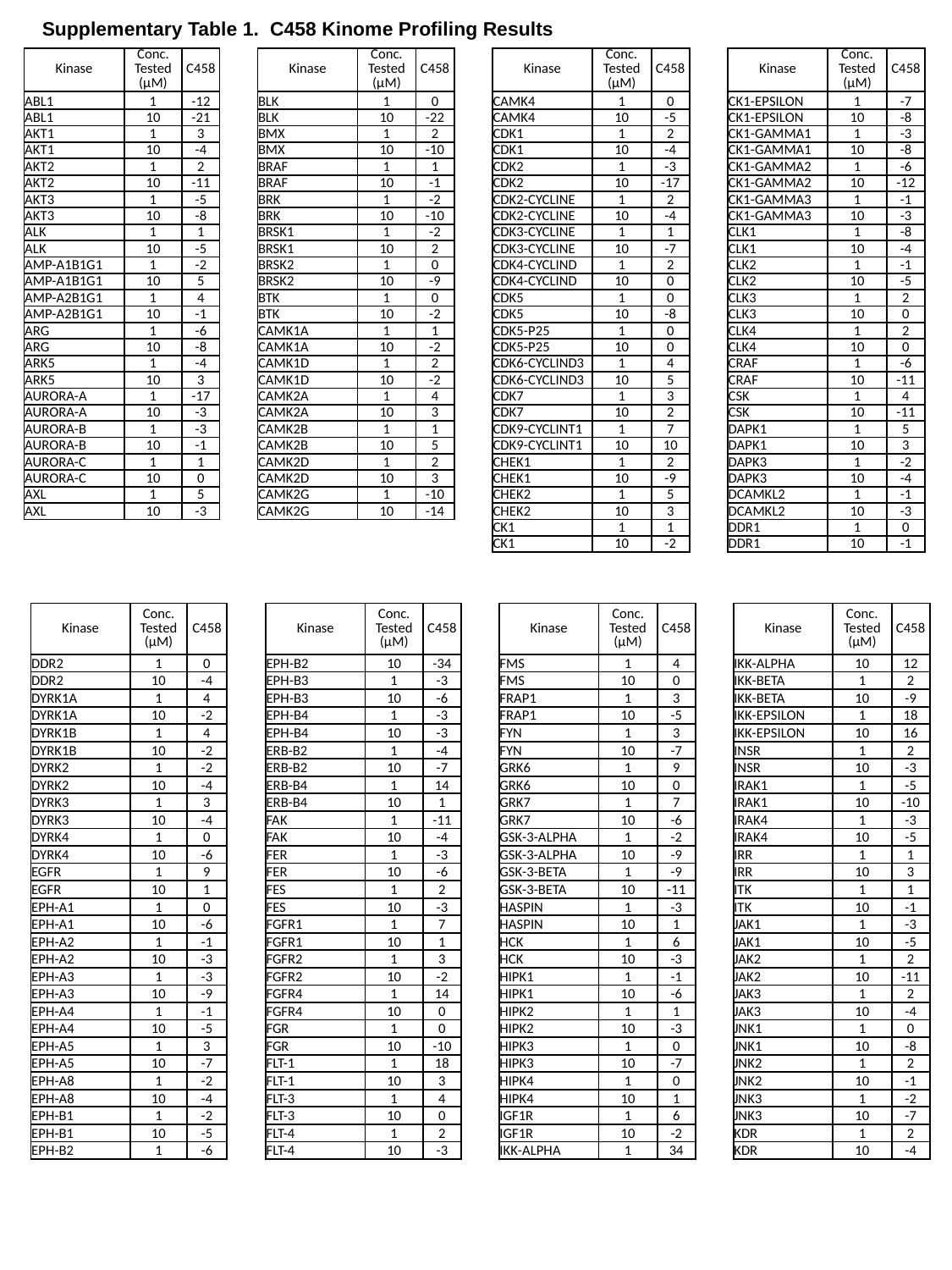

Supplementary Table 1. C458 Kinome Profiling Results
| Kinase | Conc. Tested (μM) | C458 | | Kinase | Conc. Tested (μM) | C458 | | Kinase | Conc. Tested (μM) | C458 | | Kinase | Conc. Tested (μM) | C458 |
| --- | --- | --- | --- | --- | --- | --- | --- | --- | --- | --- | --- | --- | --- | --- |
| ABL1 | 1 | -12 | | BLK | 1 | 0 | | CAMK4 | 1 | 0 | | CK1-EPSILON | 1 | -7 |
| ABL1 | 10 | -21 | | BLK | 10 | -22 | | CAMK4 | 10 | -5 | | CK1-EPSILON | 10 | -8 |
| AKT1 | 1 | 3 | | BMX | 1 | 2 | | CDK1 | 1 | 2 | | CK1-GAMMA1 | 1 | -3 |
| AKT1 | 10 | -4 | | BMX | 10 | -10 | | CDK1 | 10 | -4 | | CK1-GAMMA1 | 10 | -8 |
| AKT2 | 1 | 2 | | BRAF | 1 | 1 | | CDK2 | 1 | -3 | | CK1-GAMMA2 | 1 | -6 |
| AKT2 | 10 | -11 | | BRAF | 10 | -1 | | CDK2 | 10 | -17 | | CK1-GAMMA2 | 10 | -12 |
| AKT3 | 1 | -5 | | BRK | 1 | -2 | | CDK2-CYCLINE | 1 | 2 | | CK1-GAMMA3 | 1 | -1 |
| AKT3 | 10 | -8 | | BRK | 10 | -10 | | CDK2-CYCLINE | 10 | -4 | | CK1-GAMMA3 | 10 | -3 |
| ALK | 1 | 1 | | BRSK1 | 1 | -2 | | CDK3-CYCLINE | 1 | 1 | | CLK1 | 1 | -8 |
| ALK | 10 | -5 | | BRSK1 | 10 | 2 | | CDK3-CYCLINE | 10 | -7 | | CLK1 | 10 | -4 |
| AMP-A1B1G1 | 1 | -2 | | BRSK2 | 1 | 0 | | CDK4-CYCLIND | 1 | 2 | | CLK2 | 1 | -1 |
| AMP-A1B1G1 | 10 | 5 | | BRSK2 | 10 | -9 | | CDK4-CYCLIND | 10 | 0 | | CLK2 | 10 | -5 |
| AMP-A2B1G1 | 1 | 4 | | BTK | 1 | 0 | | CDK5 | 1 | 0 | | CLK3 | 1 | 2 |
| AMP-A2B1G1 | 10 | -1 | | BTK | 10 | -2 | | CDK5 | 10 | -8 | | CLK3 | 10 | 0 |
| ARG | 1 | -6 | | CAMK1A | 1 | 1 | | CDK5-P25 | 1 | 0 | | CLK4 | 1 | 2 |
| ARG | 10 | -8 | | CAMK1A | 10 | -2 | | CDK5-P25 | 10 | 0 | | CLK4 | 10 | 0 |
| ARK5 | 1 | -4 | | CAMK1D | 1 | 2 | | CDK6-CYCLIND3 | 1 | 4 | | CRAF | 1 | -6 |
| ARK5 | 10 | 3 | | CAMK1D | 10 | -2 | | CDK6-CYCLIND3 | 10 | 5 | | CRAF | 10 | -11 |
| AURORA-A | 1 | -17 | | CAMK2A | 1 | 4 | | CDK7 | 1 | 3 | | CSK | 1 | 4 |
| AURORA-A | 10 | -3 | | CAMK2A | 10 | 3 | | CDK7 | 10 | 2 | | CSK | 10 | -11 |
| AURORA-B | 1 | -3 | | CAMK2B | 1 | 1 | | CDK9-CYCLINT1 | 1 | 7 | | DAPK1 | 1 | 5 |
| AURORA-B | 10 | -1 | | CAMK2B | 10 | 5 | | CDK9-CYCLINT1 | 10 | 10 | | DAPK1 | 10 | 3 |
| AURORA-C | 1 | 1 | | CAMK2D | 1 | 2 | | CHEK1 | 1 | 2 | | DAPK3 | 1 | -2 |
| AURORA-C | 10 | 0 | | CAMK2D | 10 | 3 | | CHEK1 | 10 | -9 | | DAPK3 | 10 | -4 |
| AXL | 1 | 5 | | CAMK2G | 1 | -10 | | CHEK2 | 1 | 5 | | DCAMKL2 | 1 | -1 |
| AXL | 10 | -3 | | CAMK2G | 10 | -14 | | CHEK2 | 10 | 3 | | DCAMKL2 | 10 | -3 |
| | | | | | | | | CK1 | 1 | 1 | | DDR1 | 1 | 0 |
| | | | | | | | | CK1 | 10 | -2 | | DDR1 | 10 | -1 |
| Kinase | Conc. Tested (μM) | C458 | | Kinase | Conc. Tested (μM) | C458 | | Kinase | Conc. Tested (μM) | C458 | | Kinase | Conc. Tested (μM) | C458 |
| --- | --- | --- | --- | --- | --- | --- | --- | --- | --- | --- | --- | --- | --- | --- |
| DDR2 | 1 | 0 | | EPH-B2 | 10 | -34 | | FMS | 1 | 4 | | IKK-ALPHA | 10 | 12 |
| DDR2 | 10 | -4 | | EPH-B3 | 1 | -3 | | FMS | 10 | 0 | | IKK-BETA | 1 | 2 |
| DYRK1A | 1 | 4 | | EPH-B3 | 10 | -6 | | FRAP1 | 1 | 3 | | IKK-BETA | 10 | -9 |
| DYRK1A | 10 | -2 | | EPH-B4 | 1 | -3 | | FRAP1 | 10 | -5 | | IKK-EPSILON | 1 | 18 |
| DYRK1B | 1 | 4 | | EPH-B4 | 10 | -3 | | FYN | 1 | 3 | | IKK-EPSILON | 10 | 16 |
| DYRK1B | 10 | -2 | | ERB-B2 | 1 | -4 | | FYN | 10 | -7 | | INSR | 1 | 2 |
| DYRK2 | 1 | -2 | | ERB-B2 | 10 | -7 | | GRK6 | 1 | 9 | | INSR | 10 | -3 |
| DYRK2 | 10 | -4 | | ERB-B4 | 1 | 14 | | GRK6 | 10 | 0 | | IRAK1 | 1 | -5 |
| DYRK3 | 1 | 3 | | ERB-B4 | 10 | 1 | | GRK7 | 1 | 7 | | IRAK1 | 10 | -10 |
| DYRK3 | 10 | -4 | | FAK | 1 | -11 | | GRK7 | 10 | -6 | | IRAK4 | 1 | -3 |
| DYRK4 | 1 | 0 | | FAK | 10 | -4 | | GSK-3-ALPHA | 1 | -2 | | IRAK4 | 10 | -5 |
| DYRK4 | 10 | -6 | | FER | 1 | -3 | | GSK-3-ALPHA | 10 | -9 | | IRR | 1 | 1 |
| EGFR | 1 | 9 | | FER | 10 | -6 | | GSK-3-BETA | 1 | -9 | | IRR | 10 | 3 |
| EGFR | 10 | 1 | | FES | 1 | 2 | | GSK-3-BETA | 10 | -11 | | ITK | 1 | 1 |
| EPH-A1 | 1 | 0 | | FES | 10 | -3 | | HASPIN | 1 | -3 | | ITK | 10 | -1 |
| EPH-A1 | 10 | -6 | | FGFR1 | 1 | 7 | | HASPIN | 10 | 1 | | JAK1 | 1 | -3 |
| EPH-A2 | 1 | -1 | | FGFR1 | 10 | 1 | | HCK | 1 | 6 | | JAK1 | 10 | -5 |
| EPH-A2 | 10 | -3 | | FGFR2 | 1 | 3 | | HCK | 10 | -3 | | JAK2 | 1 | 2 |
| EPH-A3 | 1 | -3 | | FGFR2 | 10 | -2 | | HIPK1 | 1 | -1 | | JAK2 | 10 | -11 |
| EPH-A3 | 10 | -9 | | FGFR4 | 1 | 14 | | HIPK1 | 10 | -6 | | JAK3 | 1 | 2 |
| EPH-A4 | 1 | -1 | | FGFR4 | 10 | 0 | | HIPK2 | 1 | 1 | | JAK3 | 10 | -4 |
| EPH-A4 | 10 | -5 | | FGR | 1 | 0 | | HIPK2 | 10 | -3 | | JNK1 | 1 | 0 |
| EPH-A5 | 1 | 3 | | FGR | 10 | -10 | | HIPK3 | 1 | 0 | | JNK1 | 10 | -8 |
| EPH-A5 | 10 | -7 | | FLT-1 | 1 | 18 | | HIPK3 | 10 | -7 | | JNK2 | 1 | 2 |
| EPH-A8 | 1 | -2 | | FLT-1 | 10 | 3 | | HIPK4 | 1 | 0 | | JNK2 | 10 | -1 |
| EPH-A8 | 10 | -4 | | FLT-3 | 1 | 4 | | HIPK4 | 10 | 1 | | JNK3 | 1 | -2 |
| EPH-B1 | 1 | -2 | | FLT-3 | 10 | 0 | | IGF1R | 1 | 6 | | JNK3 | 10 | -7 |
| EPH-B1 | 10 | -5 | | FLT-4 | 1 | 2 | | IGF1R | 10 | -2 | | KDR | 1 | 2 |
| EPH-B2 | 1 | -6 | | FLT-4 | 10 | -3 | | IKK-ALPHA | 1 | 34 | | KDR | 10 | -4 |

### Slide 2
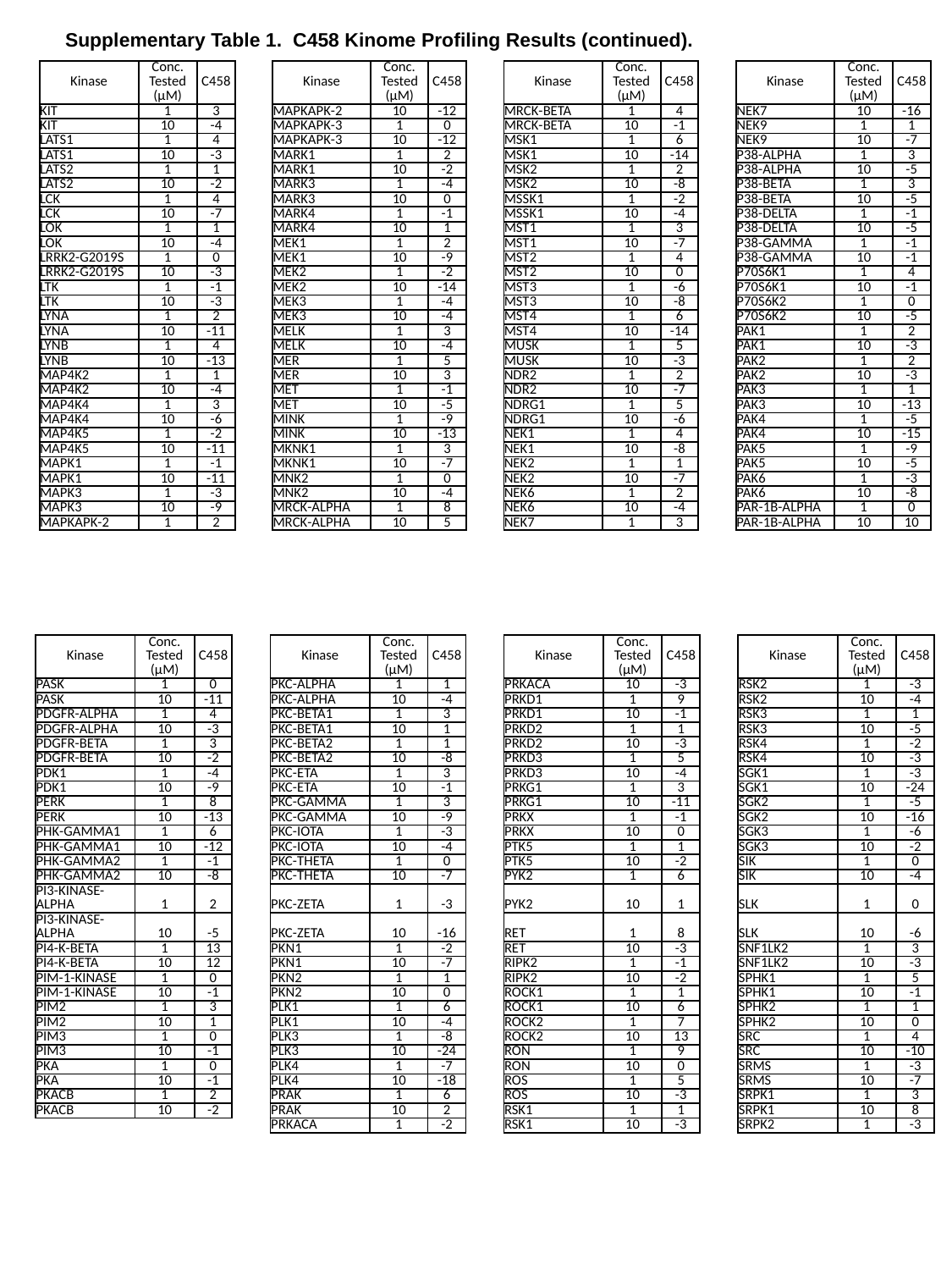

Supplementary Table 1. C458 Kinome Profiling Results (continued).
| Kinase | Conc. Tested (μM) | C458 | | Kinase | Conc. Tested (μM) | C458 | | Kinase | Conc. Tested (μM) | C458 | | Kinase | Conc. Tested (μM) | C458 |
| --- | --- | --- | --- | --- | --- | --- | --- | --- | --- | --- | --- | --- | --- | --- |
| KIT | 1 | 3 | | MAPKAPK-2 | 10 | -12 | | MRCK-BETA | 1 | 4 | | NEK7 | 10 | -16 |
| KIT | 10 | -4 | | MAPKAPK-3 | 1 | 0 | | MRCK-BETA | 10 | -1 | | NEK9 | 1 | 1 |
| LATS1 | 1 | 4 | | MAPKAPK-3 | 10 | -12 | | MSK1 | 1 | 6 | | NEK9 | 10 | -7 |
| LATS1 | 10 | -3 | | MARK1 | 1 | 2 | | MSK1 | 10 | -14 | | P38-ALPHA | 1 | 3 |
| LATS2 | 1 | 1 | | MARK1 | 10 | -2 | | MSK2 | 1 | 2 | | P38-ALPHA | 10 | -5 |
| LATS2 | 10 | -2 | | MARK3 | 1 | -4 | | MSK2 | 10 | -8 | | P38-BETA | 1 | 3 |
| LCK | 1 | 4 | | MARK3 | 10 | 0 | | MSSK1 | 1 | -2 | | P38-BETA | 10 | -5 |
| LCK | 10 | -7 | | MARK4 | 1 | -1 | | MSSK1 | 10 | -4 | | P38-DELTA | 1 | -1 |
| LOK | 1 | 1 | | MARK4 | 10 | 1 | | MST1 | 1 | 3 | | P38-DELTA | 10 | -5 |
| LOK | 10 | -4 | | MEK1 | 1 | 2 | | MST1 | 10 | -7 | | P38-GAMMA | 1 | -1 |
| LRRK2-G2019S | 1 | 0 | | MEK1 | 10 | -9 | | MST2 | 1 | 4 | | P38-GAMMA | 10 | -1 |
| LRRK2-G2019S | 10 | -3 | | MEK2 | 1 | -2 | | MST2 | 10 | 0 | | P70S6K1 | 1 | 4 |
| LTK | 1 | -1 | | MEK2 | 10 | -14 | | MST3 | 1 | -6 | | P70S6K1 | 10 | -1 |
| LTK | 10 | -3 | | MEK3 | 1 | -4 | | MST3 | 10 | -8 | | P70S6K2 | 1 | 0 |
| LYNA | 1 | 2 | | MEK3 | 10 | -4 | | MST4 | 1 | 6 | | P70S6K2 | 10 | -5 |
| LYNA | 10 | -11 | | MELK | 1 | 3 | | MST4 | 10 | -14 | | PAK1 | 1 | 2 |
| LYNB | 1 | 4 | | MELK | 10 | -4 | | MUSK | 1 | 5 | | PAK1 | 10 | -3 |
| LYNB | 10 | -13 | | MER | 1 | 5 | | MUSK | 10 | -3 | | PAK2 | 1 | 2 |
| MAP4K2 | 1 | 1 | | MER | 10 | 3 | | NDR2 | 1 | 2 | | PAK2 | 10 | -3 |
| MAP4K2 | 10 | -4 | | MET | 1 | -1 | | NDR2 | 10 | -7 | | PAK3 | 1 | 1 |
| MAP4K4 | 1 | 3 | | MET | 10 | -5 | | NDRG1 | 1 | 5 | | PAK3 | 10 | -13 |
| MAP4K4 | 10 | -6 | | MINK | 1 | -9 | | NDRG1 | 10 | -6 | | PAK4 | 1 | -5 |
| MAP4K5 | 1 | -2 | | MINK | 10 | -13 | | NEK1 | 1 | 4 | | PAK4 | 10 | -15 |
| MAP4K5 | 10 | -11 | | MKNK1 | 1 | 3 | | NEK1 | 10 | -8 | | PAK5 | 1 | -9 |
| MAPK1 | 1 | -1 | | MKNK1 | 10 | -7 | | NEK2 | 1 | 1 | | PAK5 | 10 | -5 |
| MAPK1 | 10 | -11 | | MNK2 | 1 | 0 | | NEK2 | 10 | -7 | | PAK6 | 1 | -3 |
| MAPK3 | 1 | -3 | | MNK2 | 10 | -4 | | NEK6 | 1 | 2 | | PAK6 | 10 | -8 |
| MAPK3 | 10 | -9 | | MRCK-ALPHA | 1 | 8 | | NEK6 | 10 | -4 | | PAR-1B-ALPHA | 1 | 0 |
| MAPKAPK-2 | 1 | 2 | | MRCK-ALPHA | 10 | 5 | | NEK7 | 1 | 3 | | PAR-1B-ALPHA | 10 | 10 |
| Kinase | Conc. Tested (μM) | C458 | | Kinase | Conc. Tested (μM) | C458 | | Kinase | Conc. Tested (μM) | C458 | | Kinase | Conc. Tested (μM) | C458 |
| --- | --- | --- | --- | --- | --- | --- | --- | --- | --- | --- | --- | --- | --- | --- |
| PASK | 1 | 0 | | PKC-ALPHA | 1 | 1 | | PRKACA | 10 | -3 | | RSK2 | 1 | -3 |
| PASK | 10 | -11 | | PKC-ALPHA | 10 | -4 | | PRKD1 | 1 | 9 | | RSK2 | 10 | -4 |
| PDGFR-ALPHA | 1 | 4 | | PKC-BETA1 | 1 | 3 | | PRKD1 | 10 | -1 | | RSK3 | 1 | 1 |
| PDGFR-ALPHA | 10 | -3 | | PKC-BETA1 | 10 | 1 | | PRKD2 | 1 | 1 | | RSK3 | 10 | -5 |
| PDGFR-BETA | 1 | 3 | | PKC-BETA2 | 1 | 1 | | PRKD2 | 10 | -3 | | RSK4 | 1 | -2 |
| PDGFR-BETA | 10 | -2 | | PKC-BETA2 | 10 | -8 | | PRKD3 | 1 | 5 | | RSK4 | 10 | -3 |
| PDK1 | 1 | -4 | | PKC-ETA | 1 | 3 | | PRKD3 | 10 | -4 | | SGK1 | 1 | -3 |
| PDK1 | 10 | -9 | | PKC-ETA | 10 | -1 | | PRKG1 | 1 | 3 | | SGK1 | 10 | -24 |
| PERK | 1 | 8 | | PKC-GAMMA | 1 | 3 | | PRKG1 | 10 | -11 | | SGK2 | 1 | -5 |
| PERK | 10 | -13 | | PKC-GAMMA | 10 | -9 | | PRKX | 1 | -1 | | SGK2 | 10 | -16 |
| PHK-GAMMA1 | 1 | 6 | | PKC-IOTA | 1 | -3 | | PRKX | 10 | 0 | | SGK3 | 1 | -6 |
| PHK-GAMMA1 | 10 | -12 | | PKC-IOTA | 10 | -4 | | PTK5 | 1 | 1 | | SGK3 | 10 | -2 |
| PHK-GAMMA2 | 1 | -1 | | PKC-THETA | 1 | 0 | | PTK5 | 10 | -2 | | SIK | 1 | 0 |
| PHK-GAMMA2 | 10 | -8 | | PKC-THETA | 10 | -7 | | PYK2 | 1 | 6 | | SIK | 10 | -4 |
| PI3-KINASE-ALPHA | 1 | 2 | | PKC-ZETA | 1 | -3 | | PYK2 | 10 | 1 | | SLK | 1 | 0 |
| PI3-KINASE-ALPHA | 10 | -5 | | PKC-ZETA | 10 | -16 | | RET | 1 | 8 | | SLK | 10 | -6 |
| PI4-K-BETA | 1 | 13 | | PKN1 | 1 | -2 | | RET | 10 | -3 | | SNF1LK2 | 1 | 3 |
| PI4-K-BETA | 10 | 12 | | PKN1 | 10 | -7 | | RIPK2 | 1 | -1 | | SNF1LK2 | 10 | -3 |
| PIM-1-KINASE | 1 | 0 | | PKN2 | 1 | 1 | | RIPK2 | 10 | -2 | | SPHK1 | 1 | 5 |
| PIM-1-KINASE | 10 | -1 | | PKN2 | 10 | 0 | | ROCK1 | 1 | 1 | | SPHK1 | 10 | -1 |
| PIM2 | 1 | 3 | | PLK1 | 1 | 6 | | ROCK1 | 10 | 6 | | SPHK2 | 1 | 1 |
| PIM2 | 10 | 1 | | PLK1 | 10 | -4 | | ROCK2 | 1 | 7 | | SPHK2 | 10 | 0 |
| PIM3 | 1 | 0 | | PLK3 | 1 | -8 | | ROCK2 | 10 | 13 | | SRC | 1 | 4 |
| PIM3 | 10 | -1 | | PLK3 | 10 | -24 | | RON | 1 | 9 | | SRC | 10 | -10 |
| PKA | 1 | 0 | | PLK4 | 1 | -7 | | RON | 10 | 0 | | SRMS | 1 | -3 |
| PKA | 10 | -1 | | PLK4 | 10 | -18 | | ROS | 1 | 5 | | SRMS | 10 | -7 |
| PKACB | 1 | 2 | | PRAK | 1 | 6 | | ROS | 10 | -3 | | SRPK1 | 1 | 3 |
| PKACB | 10 | -2 | | PRAK | 10 | 2 | | RSK1 | 1 | 1 | | SRPK1 | 10 | 8 |
| | | | | PRKACA | 1 | -2 | | RSK1 | 10 | -3 | | SRPK2 | 1 | -3 |

### Slide 3
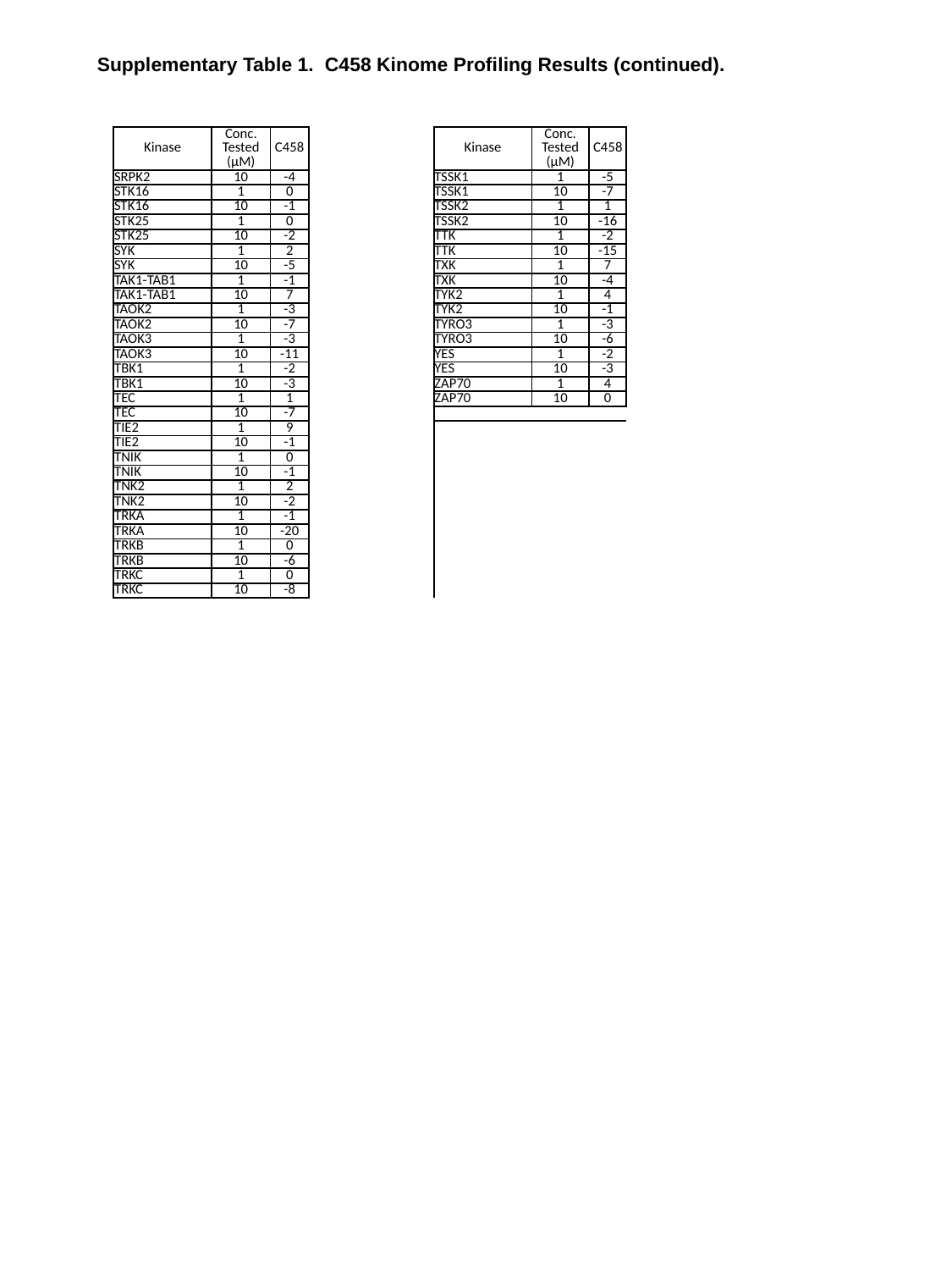

Supplementary Table 1. C458 Kinome Profiling Results (continued).
| Kinase | Conc. Tested (μM) | C458 | | Kinase | Conc. Tested (μM) | C458 |
| --- | --- | --- | --- | --- | --- | --- |
| SRPK2 | 10 | -4 | | TSSK1 | 1 | -5 |
| STK16 | 1 | 0 | | TSSK1 | 10 | -7 |
| STK16 | 10 | -1 | | TSSK2 | 1 | 1 |
| STK25 | 1 | 0 | | TSSK2 | 10 | -16 |
| STK25 | 10 | -2 | | TTK | 1 | -2 |
| SYK | 1 | 2 | | TTK | 10 | -15 |
| SYK | 10 | -5 | | TXK | 1 | 7 |
| TAK1-TAB1 | 1 | -1 | | TXK | 10 | -4 |
| TAK1-TAB1 | 10 | 7 | | TYK2 | 1 | 4 |
| TAOK2 | 1 | -3 | | TYK2 | 10 | -1 |
| TAOK2 | 10 | -7 | | TYRO3 | 1 | -3 |
| TAOK3 | 1 | -3 | | TYRO3 | 10 | -6 |
| TAOK3 | 10 | -11 | | YES | 1 | -2 |
| TBK1 | 1 | -2 | | YES | 10 | -3 |
| TBK1 | 10 | -3 | | ZAP70 | 1 | 4 |
| TEC | 1 | 1 | | ZAP70 | 10 | 0 |
| TEC | 10 | -7 | | | | |
| TIE2 | 1 | 9 | | | | |
| TIE2 | 10 | -1 | | | | |
| TNIK | 1 | 0 | | | | |
| TNIK | 10 | -1 | | | | |
| TNK2 | 1 | 2 | | | | |
| TNK2 | 10 | -2 | | | | |
| TRKA | 1 | -1 | | | | |
| TRKA | 10 | -20 | | | | |
| TRKB | 1 | 0 | | | | |
| TRKB | 10 | -6 | | | | |
| TRKC | 1 | 0 | | | | |
| TRKC | 10 | -8 | | | | |
